# REPEATED MARINE-TO-FRESHWATER FISH TRANSITIONS REVEAL PALEOENVIRONMENTAL MODULATION OF ADAPTIVE RADIATION

**DOI:** 10.64898/2026.05.26.728014

**Authors:** Aline P. M. Medeiros, Melissa Rincon-Sandoval, Aaron Davis, Aintzane Santaquiteria, Christine E. Thacker, Joshua P. Egan, Jonathan Kim, Alfred Ko’ou, Dahiana Arcila, William B. Ludt, Lily C. Hughes, Devin Bloom, Ricardo Betancur-R

**Author notes:** These authors contributed equally to this work (Aline P. M. Medeiros, Melissa Rincon-Sandoval, Ricardo Betancur-R). Corresponding author: Ricardo Betancur; Scripps Institution of Oceanography, University of California San Diego, 8622 Kennel Way, La Jolla, CA 92037, USA.

## Abstract

Tropical rivers in Australia and New Guinea (Sahul) provide a rare natural experiment in vertebrate evolution: unlike other continental systems, their freshwater ichthyofaunas are composed almost entirely of marine-derived lineages rather than primary freshwater fishes. This unique biogeographic setting enables replicated tests of why some marine-to-freshwater transitions give rise to extensive adaptive radiations whereas others remain species-poor, and whether these outcomes reflect ecological opportunity or temporally structured paleoenvironmental constraints. Using a densely sampled, time-calibrated phylogenomic framework spanning 2,303 teleost species, we identified a likely range of 27–34 marine-to-freshwater transitions during the Cenozoic, including a pronounced Middle Miocene peak (16–11 Ma). Although ecological opportunity in Sahul rivers enabled repeated colonization in the absence of dominant primary freshwater incumbents, younger freshwater lineages nevertheless diversify faster than older ones, contradicting the expectation that early arrivers should undergo elevated diversification when accessing vacant niche space. Although some colonizations coincide with bursts of speciation consistent with adaptive radiation, many yielded few species despite long residence times. Functional trait analyses likewise revealed no consistent relationship between colonization timing, ecological breadth, or diversification rate, although expanded functional space characterizes previously proposed Sahul adaptive radiations. Comparisons with paleoenvironmental curves indicate that colonization success correlates with sea-level minima and low-oxygen conditions, suggesting that Earth history dynamics modulated when ecological opportunity was accessible. Our results show that although ecological opportunity enabled repeated freshwater invasions into the Sahul region, diversification outcomes are governed by the interaction of paleoenvironmental dynamics and possibly lineage-specific traits, generating stark asymmetries in freshwater radiations.

**Significance statement:** Tropical rivers in Australia and New Guinea host one of the most unusual continental freshwater fish assemblages on Earth, composed almost entirely of marine-derived lineages. This system allows asking why some colonizing lineages diversify dramatically while others remain species-poor on a continental scale. Using large-scale phylogenomic and functional trait data, we show that early arrival alone does not predict diversification success. Instead, the lineages that radiate most successfully are those whose arrival coincides with windows of paleoenvironmental opportunity created by sea-level and oxygen fluctuations. These results reveal that the fates of colonizing lineages are shaped not only by ecological opportunity, but also by Earth-history dynamics that govern when, where, and how species can invade and diversify.

## Introduction

Tropical rivers in Sahul, the continental landmass encompassing Australia and New Guinea, represent a natural laboratory for studying diversification dynamics in the absence of dominant incumbents (i.e., resident freshwater lineages already established in the habitat when a new colonizer arrives). Much as marsupials radiated in the region in the absence of placental mammals, Sahul freshwater systems lack most of the primary freshwater fish clades that dominate rivers in South America, Africa, and Eurasia (e.g., characins, cyprinids, knifefishes, and most catfishes) (1, 2). Although the fossil record indicates that primary freshwater fishes were more diverse in the past, extant tropical freshwater faunas in Sahul now include only a few relict primary lineages, such as lungfishes and bonytongues, leaving large portions of the ecological space occupied by freshwater fishes on other continents weakly represented or absent today (3). Extinct relatives of these relicts, together with other ancient freshwater fish lineages (e.g., early ray-finned and stem teleost lineages), were progressively lost from Sahul between the Late Cretaceous and the Paleogene, so that primary freshwater fish communities were already depleted before the major Cenozoic assembly of the modern freshwater fauna (4, 5). Sahul riverine systems were subsequently populated largely through repeated colonizations by marine-derived teleost fishes, which today account for more than 97% of the region’s extant freshwater ichthyofauna, spanning ∼54 families and ca. 620 species (Fig. 1) (3–6).

**Figure 1.**
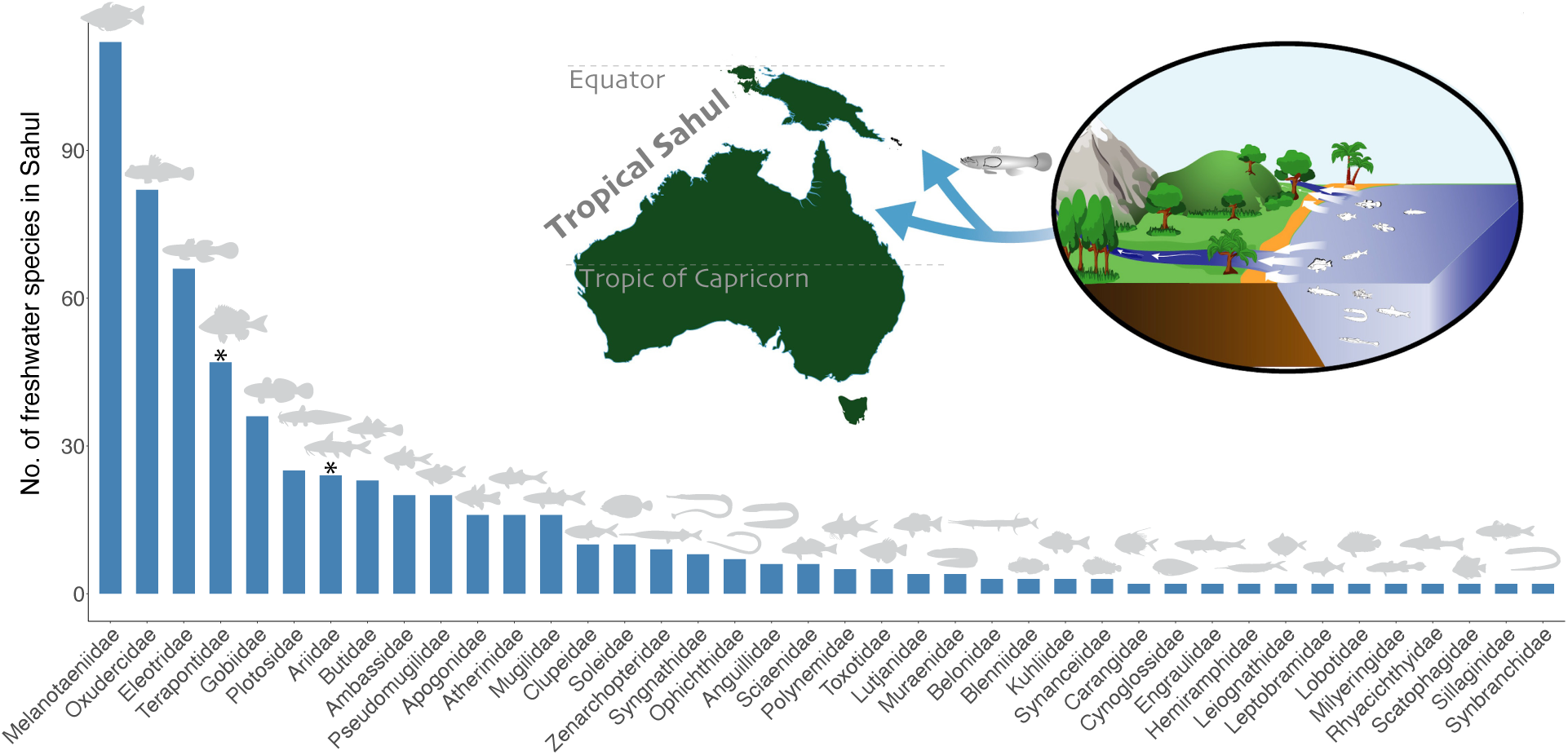
Diversity of marine-derived freshwater teleost fish clades in tropical rivers of Australia and New Guinea (Tropical Sahul; ca. 620 species). Bars indicate the number of species recorded from freshwater habitats for each family represented by more than one species in Sahul, including peripheral species that occur in freshwater, brackish and marine environments (22). Asterisks denote families previously shown to exhibit signatures consistent with adaptive radiations in Sahul freshwater based on phylogenetic comparative analyses (11, 23). Primary freshwater fishes (e.g., Osteoglossidae; 2 species) and temperate freshwater groups (e.g., Galaxiidae) fall outside the tropical, marine-derived scope of this study and are therefore not shown.

Despite similar ancestral ecologies and colonization routes, the evolutionary outcomes of marine-to-freshwater transitions in Sahul have been remarkably uneven (Fig. 1). Some groups, such as rainbowfishes (Melanotaeniidae), gudgeons (Eleotridae), grunters (Terapontidae), and sea catfishes (Ariidae), have diversified extensively, occupying a broad functional spectrum of hydrological regimes and trophic axes (7, 9–12), whereas other clades, including needlefishes (Belonidae), halfbeaks (Hemiramphidae), and croakers (Sciaenidae), contain relatively few species and limited ecological diversity despite comparable or longer occupancy of freshwater habitats (5, 11, 12). These transitions unfolded against a backdrop of major long-term environmental change in Australia, including progressive continental aridification beginning near the Eocene-Oligocene transition (∼33 Ma), which profoundly reshaped freshwater availability and drainage structure in Sahul (15, 16).

Classic macroevolutionary theory attributes such disparities in diversity to ecological opportunity, whereby colonization of species-poor environments with underutilized resources can promote rapid taxonomic and functional diversification (13, 14). Under this framework, two related predictions follow. First, arrival order should matter: lineages that colonized Sahul freshwater earlier (“early arrivers”) are expected to experience higher diversification rates than lineages that colonize freshwater later (“late comers”) and encounter established incumbents (7). Second, ecological opportunity also predicts expansion into novel regions of functional space, such that early arrivers should occupy broader and more disparate functional trait space than late comers, whose opportunities are constrained by competition with incumbents. Together, these predictions contrast two scenarios: under priority effects, late comers occupy trait space already filled by early arrivers; under open-ended radiation, each invasion accesses new functional space (19).

An alternative, which we call the paleoenvironmental opportunity hypothesis, proposes that diversification success reflects the dynamic geological history of the Sahul region, including tectonic uplift, fluctuating sea levels, and drainage reorganizations that repeatedly connected and fragmented coastal and inland habitats (3, 6, 20, 21). Under this scenario, episodes of colonization and diversification are expected to track periods when changing seaways or drainage connections enabled invasion and subsequent isolation, meaning that diversification dynamics are governed less by arrival order and more by the timing of paleoenvironmental change. These hypotheses are not mutually exclusive: ecological opportunity may itself have been contingent upon the temporal windows generated by paleoenvironmental restructuring.

Taken together, the repeated origins of freshwater colonization among distantly related marine lineages provide an exceptional opportunity to evaluate how ecological and paleoenvironmental opportunity shape diversification in a replicated natural experiment. Here, we evaluate these alternative expectations using a newly constructed, time-calibrated phylogenomic framework for 2,303 teleost species. By integrating ancestral habitat reconstruction, diversification models, and comparative analyses of broad ecological trait space, we assess whether diversification in Sahul freshwater fishes is better explained by arrival order and incumbency effects or by temporally structured paleoenvironmental windows that modulated access to, and isolation in, river systems. Specifically, we reconstruct the timing and frequency of marine-to-freshwater transitions, relate diversification rates to colonization history, and test whether patterns of functional diversification and colonization pulses align with major episodes of geological and environmental restructuring in Sahul.

## Results

### A densely sampled phylogenomic framework reveals a Middle Miocene peak in marine-to-freshwater colonizations in Sahul

We constructed a set of 30 densely sampled, time-calibrated phylogenies of teleost fishes to determine when and in which lineages marine-to-freshwater transitions into Sahul occurred. This framework integrates a phylogenomic backbone, expanded taxonomic sampling with legacy markers, and grafting of subclade trees from recent studies (Tables S1–S6), placing up to 2,303 species in the full trees used for diversification analyses. These include focal Sahul freshwater (356 of ca. 620 species; 57%) clades and their marine close relatives (477 species). Divergence times were estimated using secondary calibrations (Tables S4–S5), yielding a consistent temporal scale for comparison across clades (Fig. 2a). Using the set of 30 trees, including two master trees estimated through concatenation (RAxML (24)) and coalescent (ASTRAL (25)) approaches, we reconstructed ancestral habitats in BioGeoBEARS (26) based on habitat assignments compiled from FishBase (hereafter FB; (27)) and the Catalog of Fishes (hereafter CoF; (8)). Because these databases differ in the classification of freshwater, marine, and euryhaline species, we accounted for these differences by generating alternative habitat datasets, including a “Fresh+” dataset (including brackish species) and a “Fresh Only” dataset (excluding brackish species), and reconciling conflicting records through manual curation based on the primary literature and expert input (Tables S7–S13). Reconstructions were performed on pruned versions of these trees, retaining 1,677 species, including all marine and brackish taxa but only freshwater species occurring in Sahul (Fig. 2a), thereby restricting inferred marine-to-freshwater transitions to Sahul river systems. Across all datasets, the BAYAREALIKE+J model was consistently preferred (Tables S20–S23). We summarized ancestral range estimates from all 30 trees by overlaying average probabilities across compatible nodes on the master trees under the best-fit model (26). We then estimated the timing and frequency of independent colonization events by averaging 100 stochastic histories per analysis on the master trees under the same model (following Xing and Ree (28)).

**Figure 2.**
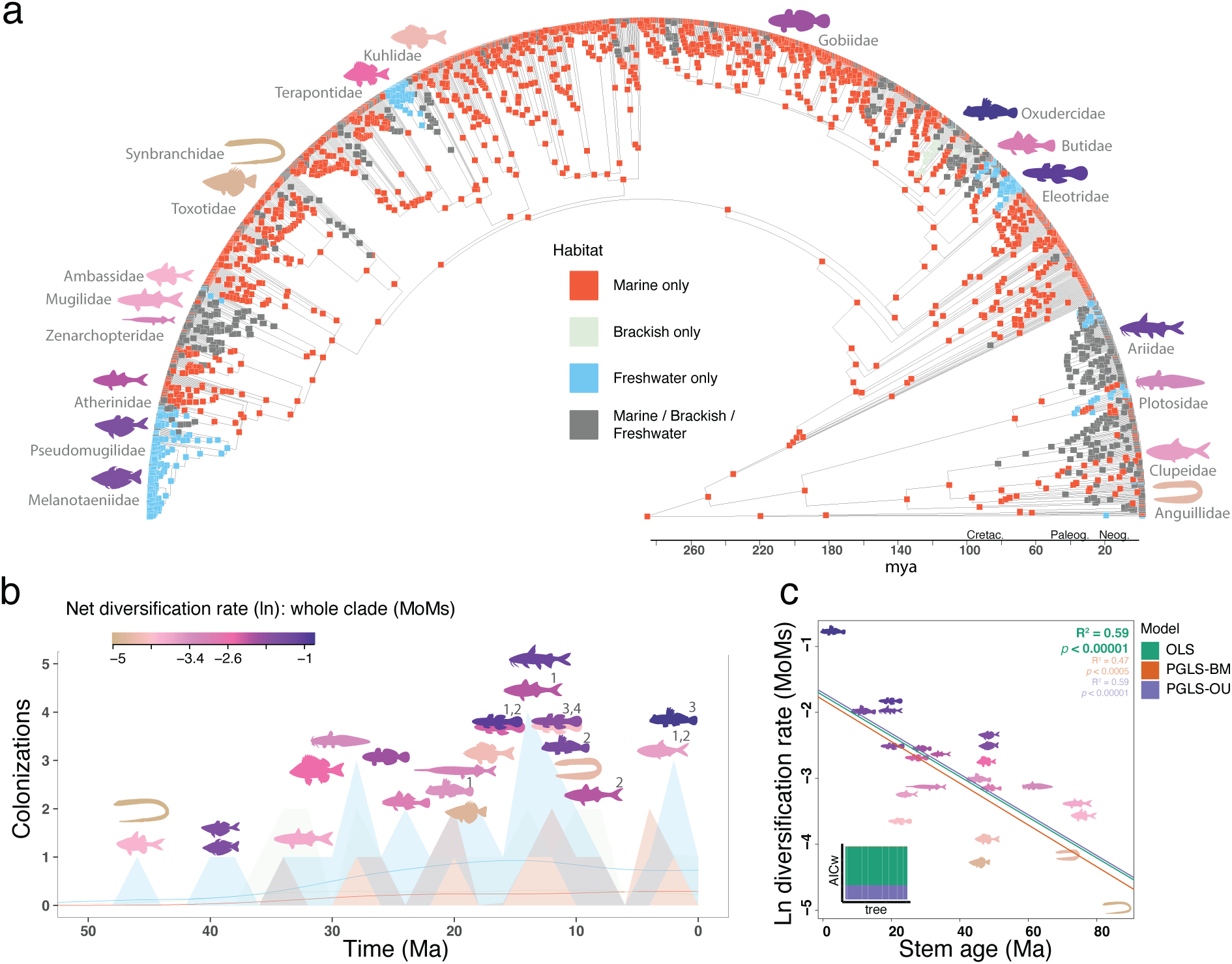
Phylogeny, ancestral habitat reconstructions, colonization dynamics, and diversification rates of Sahul freshwater fishes. (a) Ancestral habitat reconstructions for 1,677 species in the FishBase (FB) reconciled Fresh+ dataset (the full 2,303-species trees were used for diversification and ClaDS analyses; see Fig. 3). Node colors indicate inferred habitat occupancy, and fish silhouettes mark families with marine-to-freshwater transitions in Sahul rivers (see Figs. S6–S11 for alternative datasets). (b) Colonization-through-time plot for the Cenozoic (56–2 Ma) based on 100 stochastic maps from the ASTRAL master tree and the FB reconciled Fresh+ dataset. Fish silhouettes are positioned by the inferred age of freshwater transition and colored by whole-clade net diversification rate (Method-of-Moments; MoMs). Numbers above silhouettes indicate the number of independent marine-to-freshwater transitions inferred for each family or subclade, highlighting repeated transitions within lineages. (c) Ordinary least squares (OLS) and phylogenetic generalized least-squares (PGLS) regressions under Brownian motion (PGLS-BM) and Ornstein-Uhlenbeck (PGLS-OU) models testing the relationship between subclade stem age and net diversification rate from MoMs (see Fig. S14d for crown age). Akaike weights (AICw) for alternative models are summarized across trees with bar plots. Averaged R² and p values are shown in the upper right, with the best-fitting model in bold. Averaged stem and crown ages, diversification rates, and functional metrics for each subclade are provided in Table S26.

Throughout most of the Cenozoic (56–2 Ma), the maximum number of independent freshwater colonization events (including singletons, represented by a single freshwater species; i.e., a colonization event with no subsequent diversification in freshwaters) ranged between 21–38 (Table S24), with most estimates falling within the 27–34 range and averaging ∼0.5 events per 2-Ma time bin (Table S16). To focus on *in-situ* diversification (singletons excluded), we retained clades with at least two descendant Sahul marine-derived freshwater species. Such clades were conservatively defined as the first node with a high likelihood of freshwater occupancy (Figs. 2a, S8). The number of retained colonization events differed between datasets, with 23 events in the FB Reconciled Fresh+ dataset versus 14 in the CoF Reconciled Fresh+ dataset. This discrepancy was driven primarily by alternative estimates of the first freshwater origin in Gobiaria (Figs. S8–S9). The CoF dataset supports a single early transition (>50 Ma) giving rise to butids, eleotrids, oxudercids, and gobies, whereas the FB dataset infers multiple independent transitions in the last 20 Ma (Fig. 2a). We favored the latter interpretation because it is more consistent with recent evidence for relatively recent diversification of sleeper gobies (29), multiple freshwater transitions in East Asian oxudercids (30), and prior proposals of multiple freshwater origins in gobies (31). The biogeographic composition of our Gobiaria sampling is methodologically more sensitive to our study region, with ∼50% of species from Sahul and ∼30% of those in freshwater, which would also likely bias inference toward the more lumped CoF-like scenario of a single ancient transition. We therefore retained subclades concordant between datasets and, for Gobiaria, followed the FB Reconciled Fresh+ dataset, yielding a final set of 23 freshwater subclades for downstream analyses (Table S26). Independent of subclade definitions, colonization-through-time analyses revealed a pronounced Middle Miocene peak (16–11 Ma), during which marine-to-freshwater transitions were especially frequent in Eleotridae and Atherinidae *sensu lato*, and to a lesser extent in Ariidae, Oxudercidae, and Anguillidae (Figs. 2b, S12–S14). A second, weaker but consistent peak occurred during the Oligocene (30–25 Ma), associated primarily with Terapontidae, Plotosidae, and Gobiidae. A third peak in the youngest bin (5–0 Ma) is also recovered, but its magnitude varies across alternative datasets and phylogenies (Figs. S12–S13).

### Younger freshwater clades diversify faster than older ones, challenging the early arriver expectations under ecological opportunity

To test predictions derived from ecological opportunity regarding early versus late colonizers, we analyzed 17–23 independent Sahul freshwater subclades comprising at least two marine-derived freshwater species (Table S26), including both family-level clades with multiple colonizations (e.g., clupeids, gobies) as well as single-origin, multi-family radiations (e.g., Melanotaeniidae + Pseudomugilidae; Fig. 2b). We extracted stem and crown ages across 30 calibrated phylogenies (Table S14), and calculated net diversification rates using method-of-moments (MoMs; (32)) under alternative extinction settings (ε = 0, 0.5, 0.9), which produced consistent rankings across clades (Tables S27–S29). A moderate setting (ε = 0.5) was retained for downstream analyses. To account for estimator uncertainty, we also assessed diversification with MiSSE (Table S30) (33), incorporating sampling fractions across all trees, yielding strongly correlated results with MoMs (R² = 0.82; Fig. S5). Clades with higher diversification rates included Pseudomugilidae, Melanotaeniidae, Ariidae, Atherinidae, Terapontidae, and subclades within Gobiaria (Fig. 2b), a finding consistent with previously reported evolutionary radiations (7, 10, 14, 29). Overall, higher net diversification rates characterized younger freshwater clades (e.g., ariids, eleotrids, oxudercids; Fig. 2b; crown age 13–22 Ma), rather than older ones (e.g. ambassids, mugilids; Fig. 2c; crown age 34–45 Ma), contradicting the early arrivers/late comers hypothesis (priority-effect hypothesis *sensu* (19) for Sahul freshwater fishes). This pattern was further supported by regressing timing of colonization against diversification rate, revealing a strong negative correlation (Fig. 2c; R^2^ = 0.59, *p* = 0.00003; Table S33). Notably, older freshwater transitions occurred during earlier phases of Sahul’s environmental history, including intervals preceding or overlapping the onset of continental aridification in Australia (∼33 Ma), whereas many younger radiations postdate these changes (15, 16).

Closer inspection of individual subclades reveals contrasting outcomes: some colonization events (e.g., eleotrids, oxudercids, ariids) show signatures consistent with evolutionary radiations, in some cases in line with previous work documenting extensive species proliferation in Sahul rivers (11, 34). By contrast, other lineages, such as clupeids, gobiids, and other eleotrid subclades, remain species-poor despite comparable, or longer, residence times in freshwater environments (see Table S26). These patterns indicate that colonization timing alone does not predict diversification magnitude (35). Rather than being determined solely by competition between early and late colonizers (18, 36), diversification trajectories appear contingent on lineage-specific properties and the broader ecological and historical context in which colonization occurred (37, 38). Younger clades may have colonized during periods of heightened ecological opportunity associated with environmental change (39), whereas older lineages may have experienced ecological saturation or extinction-diversification dynamics that obscure initial diversification pulses (40).

### Some Sahul riverine colonizations coincide with bursts of diversification, consistent with adaptive radiation theory

To test whether marine-to-freshwater colonization was associated with elevated diversification rates, we analyzed lineage-specific diversification dynamics using ClaDS, which infers frequent, small-magnitude shifts in speciation rates across branches (41). Analyses were conducted on the full 2,303-species phylogenies using both master trees. ClaDS analyses targeted major clades encompassing focal Sahul freshwater lineages and appropriate marine outgroups, irrespective of geographic setting, including families with multiple freshwater colonization events such as Clupeidae, Eleotridae, Oxudercidae, and Atherinidae (Fig. 2). From these analyses, we focused our interpretation on the 23 freshwater subclades (Fig. 3) previously identified as showing *in situ* diversification (≥2 Sahul freshwater species). Because our taxonomic sampling is biased towards marine-derived freshwater fishes, we adjusted the sampling fraction by our focal families, keeping an overall value for the remaining tips in the phylogeny (Table S31). Complete ClaDS results for all analyzed clades are shown in Figs. S15–S25.

**Figure 3.**
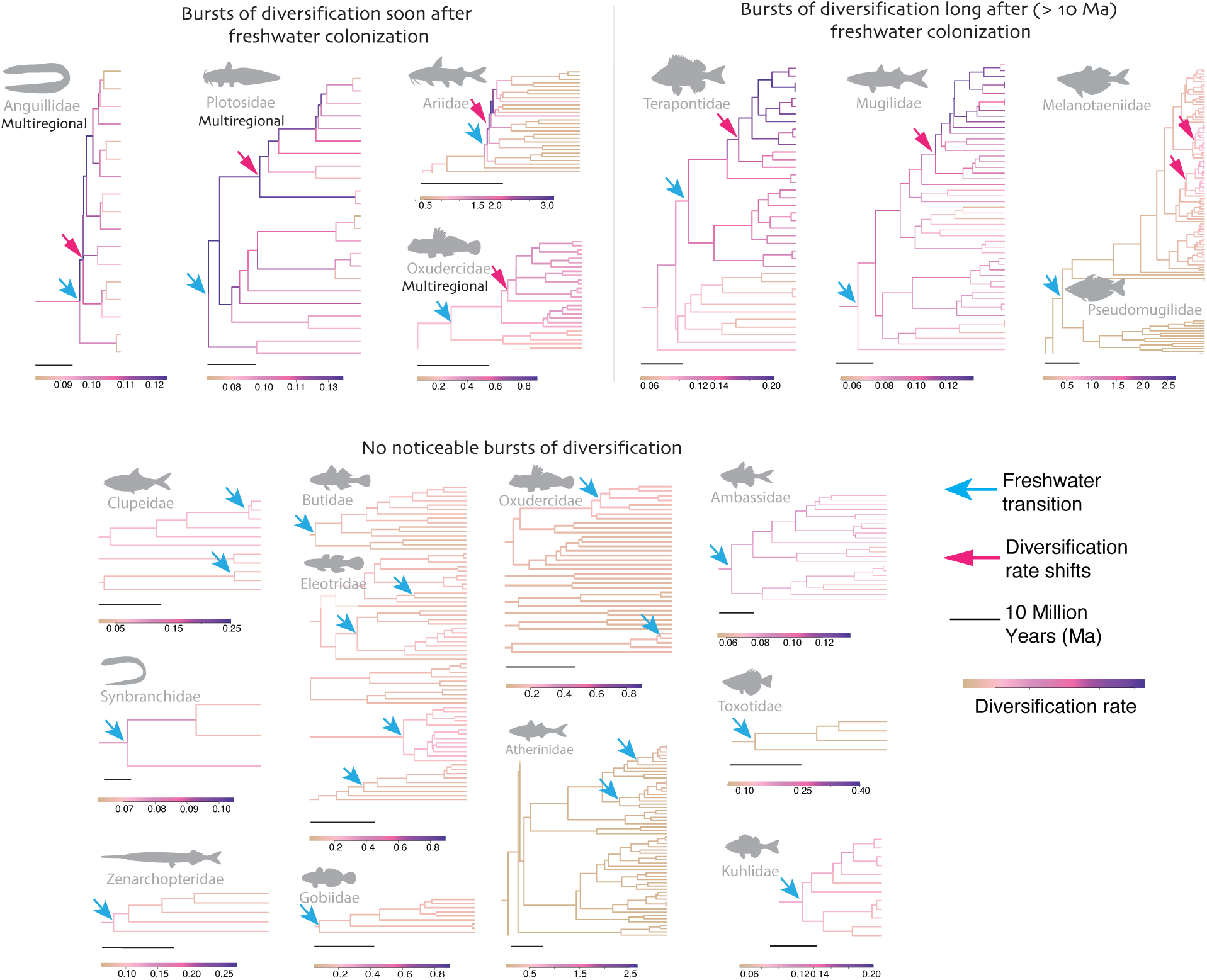
Cladogenic diversification-rate shifts (ClaDS) in Sahul fish lineages that colonized freshwater. Shown are subclades’ snippets depicting marine-to-freshwater transitions (blue arrow) and diversification rate shifts based on ClaDS (magenta arrow). The scale bar indicates 10 million years (Ma). Branch colors indicate speciation rates based on ClaDS. Complete ClaDS’ figures are available in Supplementary Material (Figs. S15–S25). Some habitat transitions are multiregional, indicating they are not unique to Sahul. Telmatherinidae, phylogenetically positioned between Melanotaeniidae and Pseudomugilidae, is not shown here due to the absence of representative Sahul taxa (see Fig. S18).

Across clades, ClaDS identified heterogeneous diversification responses following freshwater colonization (Fig. 3; Table S32). In several lineages, including Ariidae, Anguillidae, Plotosidae, and some oxudercid subclades, speciation rates increased sharply near the inferred timing of freshwater entry. In contrast, other clades exhibited little or no rate acceleration following freshwater colonization, or showed delayed increases in diversification more than 10 Ma after freshwater entry, including Terapontidae, Mugilidae, and Melanotaeniidae. Among clades with multiple independent freshwater transitions, only a subset showed elevated diversification associated with freshwater entry, indicating that repeated invasion by closely related lineages can constrain diversification due to ecological similarity and competition among colonists. We note, however, that our analyses operate at the Sahul-regional level and do not resolve individual drainages, though historical connectivity among Sahul drainages argues against repeated invasions of fully independent systems.

### Functional trait space is largely decoupled from colonization timing and diversification, but expanded space characterizes proposed Sahul adaptive radiations

To test whether functional trait space relates to colonization timing (stem and crown ages) or diversification dynamics, we compiled body size, trophic guild, and salinity breadth trait data for 270 of 343 species spanning 23 freshwater subclades (Table S15). Functional analyses were restricted to subclades with at least three species (n = 18 clades), because hypervolume-based metrics cannot be estimated for smaller clades. Trait variation was summarized using mixed-data ordination and phylogenetically informed axes, and functional space was quantified using complementary hypervolume summaries (Tables S17–S18), with overlap and centroid distances used to assess functional similarity among clades (Fig. S26). Associations between colonization timing and diversification rate were evaluated using both non-phylogenetic and phylogenetically informed regressions across the full tree set.

Among subclades, older freshwater lineages were not more functionally diverse than younger ones (Figs. 4, S27). Contrary to the early arrivers/late comers hypothesis, lineage diversification rates showed no consistent relationship with functional space (Fig. S28), indicating that ecological differentiation is largely decoupled from both colonization timing and diversification dynamics. Despite this decoupling, five lineages (Terapontidae, Ambassidae, Mugilidae, Ariidae, and Kuhliidae) exhibit markedly expanded functional space (Fig. S26d), spanning freshwater entries from ∼50 Ma (ambassids) to ∼15 Ma (ariids, kuhliids) and further confirming that trait space is not tightly linked to colonization timing. Of these, Ariidae and Terapontidae have previously been characterized as Sahul freshwater adaptive radiations based on both lineage diversification and trophic and morphological diversity (11, 34, 42). The remaining three families are strong candidates for unrecognized adaptive radiation that warrant future investigation. All five share functional attributes of habitat generalists (Fig. 4), with broad salinity tolerance and trophic flexibility (43) favor colonization and persistence in novel environments (39, 44). Complementary similarity and centroid distance indices (Fig. S26b,c) also show high functional differentiation among them, suggesting weak competition and a capacity to exploit underutilized resources. Nonetheless, the expanded functional space of these five lineages does not alter the broader pattern that functional trait space and lineage diversification are only weakly associated among Sahul freshwater fishes.

**Figure 4.**
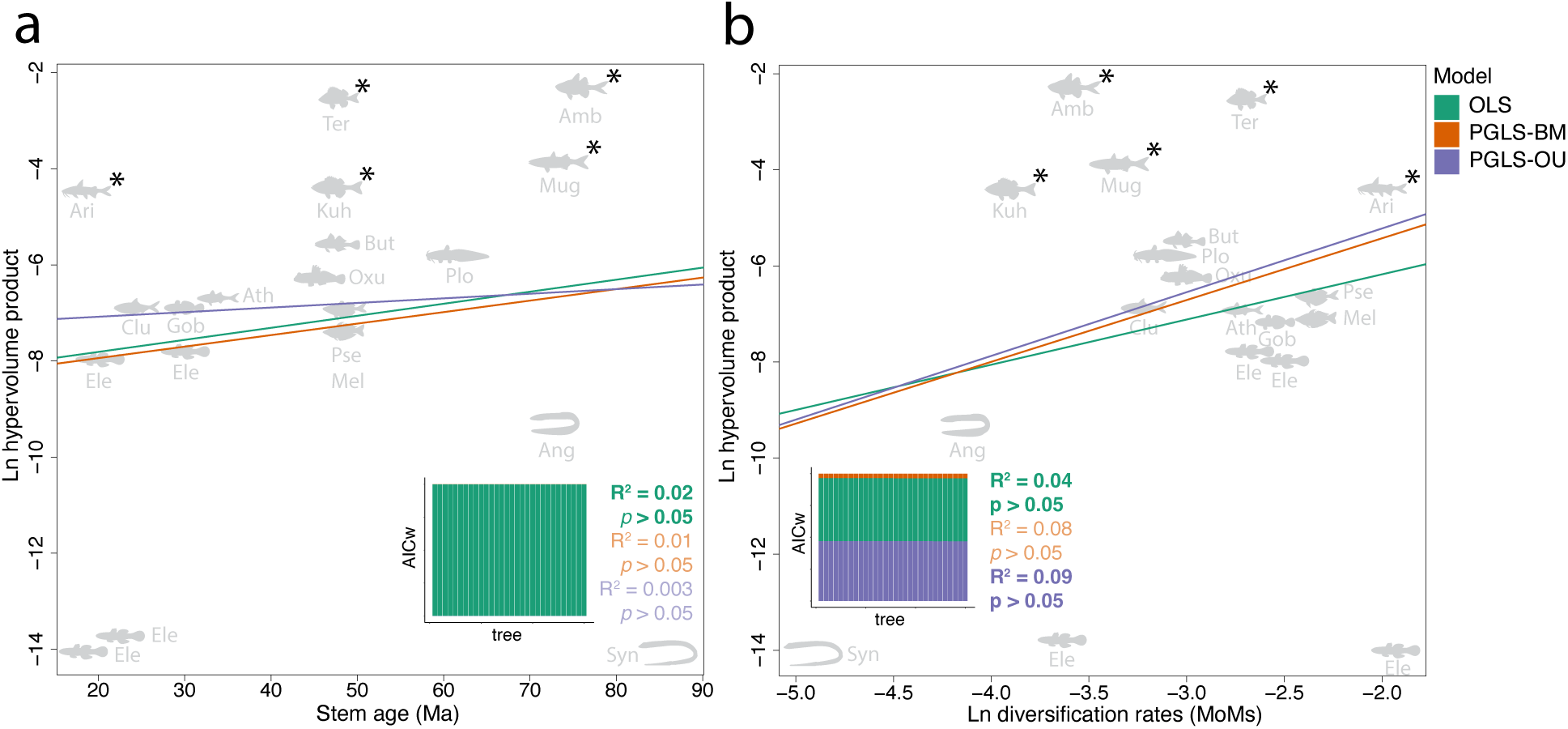
Relationship between subclade age, functional space and diversification rates, assessed using ordinary least squares (OLS) and phylogenetic generalized least squares (PGLS) regressions under Brownian motion (PGLS-BM) and Ornstein–Uhlenbeck (PGLS-OU) models. Regression between (a) stem age and functional space, and (b) diversification rates and functional space measured as hypervolume product. Fish silhouettes represent the 18 subclades for which functional space was estimated. Akaike weights (AICw) for alternative models are summarized over trees using bar plots. Averaged coefficients of determination (R²) and significance levels (*p*) are shown in the lower right (a) and left (b) corners of each panel, with the best-fit model highlighted in bold. Asterisks denote the five families (Ari: Ariidae; Ter: Terapontidae; Kuh: Kuhliidae; Amb: Ambassidae; and Mugilidae) with expanded functional space. The remaining fish shapes are represented by Anguillidae (Ang), Atherinidae (Ath), Butidae (But), Clupeidae (Clu), Eleotridae (Ele), Gobiidae (Gob), Melanotaeniidae (Mel), Oxudercidae (Oxu), Plotosidae (Plo), Pseudomugilidae (Pse), and Synbranchidae (Syn).

### Colonization events align with periods of paleo-oxygen lows and sea level fluctuations throughout the Cenozoic

Freshwater invasions have been documented in multiple taxa (45–47) and regions (48–50). Proposed drivers include marine incursions (e.g., inundation of continental land by oceanic waters; (51)) and climatic transitions during the Middle Miocene (52), which may have created ecological opportunities for marine lineages to establish and diversify in freshwater environments. To test whether Sahul freshwater invasions align with environmental shifts, we compiled paleoenvironmental time series spanning the last 56 Ma and evaluated associations using generalized additive models (GAMs; (53)). We derived freshwater colonization-through-time curves (2-Ma bins) from 2,000 stochastic mapping realizations drawn from 10 phylogenies (including ASTRAL and RAxML master trees) and two reconciled habitat datasets (FB and CoF Fresh+; Table S16). We matched these curves to eight paleoclimatic and geochemical proxies, including global and tropical temperature (proxy for thermal tolerance and habitat suitability for euryhaline lineages), sea level (shelf exposure and marine-freshwater connectivity), Australian continental inundation (availability of shallow transitional habitats), precipitation (regional aridity and drainage-scale hydrologic connectivity), salinity (osmoregulatory gradients at the marine–freshwater interface), ⁸⁷Sr/⁸⁶Sr (continental weathering intensity, reflecting riverine input to coastal habitats), and δ³⁴S (marine redox state and shallow-water oxygenation). We evaluated associations using both univariate and multivariate GAMs, incorporating immediate effects and temporal lags of 2, 4, and 6 Ma (Table S19; Fig. S29). To control for multiple testing, we adjusted *p* values using a false discovery rate (FDR) procedure and summarized corrected significance values as boxplots for each environmental variable (Fig. 5).

**Figure 5.**
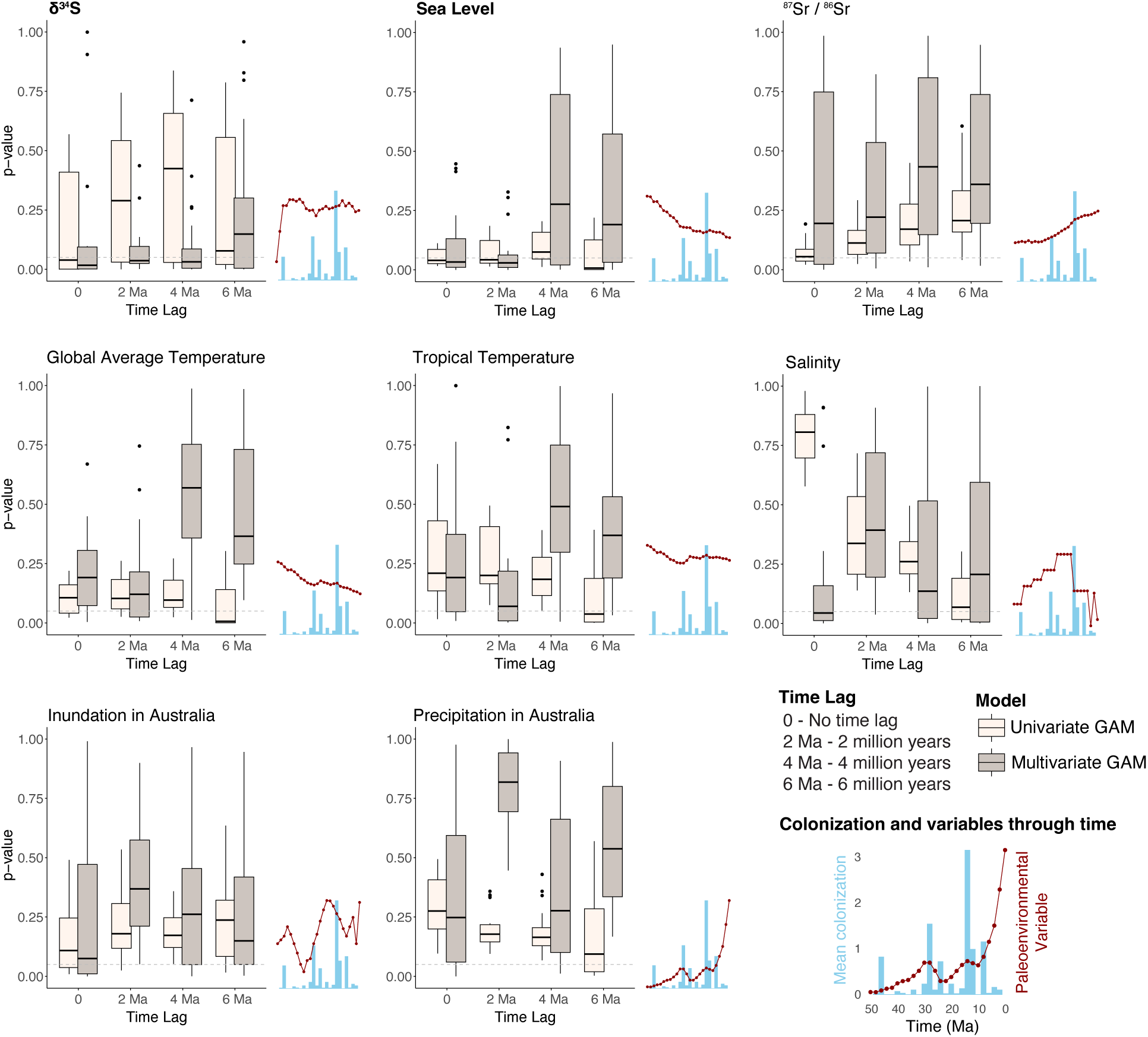
Paleoenvironmental correlates of marine-to-freshwater colonization in Sahul. Boxplots summarize results from generalized additive models testing through-time correlations between marine-to-freshwater colonization frequency and eight paleoenvironmental variables, evaluated using both univariate and multivariate models, with no time lag and with lags of 2, 4, and 6 million years (Table S19). Each boxplot represents the significance (*p*) of 20 analyses based on 10 trees and 2 subsets. Median is represented by the bold line, with the box limits representing the first (25%) and third (75%) quartiles, and whiskers indicating the upper and lower quartiles. Outliers are represented by the points beyond whiskers. Horizontal dashed lines indicate the false discovery rate (FDR)-corrected significance threshold *(p* = 0.05). Colonization events and paleoenvironmental variables through time are plotted on the right side of each boxplot. Variables whose effects repeatedly approach or exceed this threshold over trees, datasets, and lag structures are interpreted as showing consistent associations with colonization dynamics, whereas variables with isolated or inconsistent signals are not emphasized. Among these, marine redox proxies (δ³⁴S) and sea level, in bold, exhibit the most recurrent associations, suggesting that freshwater invasions tended to occur during intervals of enhanced marine-freshwater connectivity and reduced oxygen availability. All parameter’ estimates are reported in Table S19. Fig. S29 shows the colonization events with paleoenvironmental ranges through time.

Among the paleoenvironmental variables assessed, sea level and delta sulfur (δ³⁴S) were significantly correlated with the mean number of freshwater colonizations in Sahul fishes (Fig. 5). The δ³⁴S reflects the marine redox state, with higher values typically indicating more reducing (i.e. anoxic) conditions and lower values indicating more oxygenated seawater (54). Reduced oxygen in seawater may have facilitated freshwater colonization by lineages already adapted to low-oxygen conditions (e.g., elevated hemoglobin concentrations, enlarged gill surface area; (46, 55)). Sea level changes have likewise been associated with increased freshwater colonization in Sahul (3, 6). During the Miocene, sea level dropped and reshaped the Sahul Shelf, expanding shallow transitional habitats along the coast (56, 57). These dynamic estuarine-like conditions imposed a physico-chemical filter (variable salinity, shallow depths, fluctuating flow) that favored lineages capable of persisting under such regimes, facilitating freshwater transitions. Importantly, sea level dropped from the high stand of the Oligocene-Miocene Climatic Optimum (58) rather than to extreme lowstands, and the GAM detected this co-variation (Fig. S29). These results suggest that environmental context helped shape when and how marine lineages invaded freshwater systems: periods of low marine oxygen may have eased initial entry for tolerant species, whereas sea-level fluctuations first facilitated access to freshwater and later promoted isolation and diversification following basin reorganization.

## Discussion

Tropical rivers in Australia and New Guinea provide an unusually powerful test of ecological opportunity because their freshwater ichthyofaunas are dominated by marine-derived lineages rather than primary freshwater groups. The total number of freshwater transitions, including singletons, ranged from 21 to 38, with most estimates falling between 27 and 34. Of these, between 14 and 23 represent colonizations followed by *in-situ* diversification, and these events cluster prominently in the Middle Miocene (16–11 Ma; Fig. 2a). Several classic expectations from adaptive radiation and ecological opportunity theory are not met: younger freshwater clades tend to diversify faster than older ones, and functional trait space is largely decoupled from both colonization timing and diversification rate, with pronounced functional expansion restricted to a minority of families. Instead, colonization success and subsequent diversification align more closely with temporally structured paleoenvironmental windows that modulated access to rivers, including sea-level minima and low-oxygen intervals, and with lineage traits that facilitate transition and persistence, such as habitat generalist, trophic flexibility, and broad salinity tolerance. Thus, although ecological opportunity clearly enabled repeated freshwater transitions into Sahul, it was not sufficient on its own to predict radiation outcomes; diversification was governed by the interaction of time-varying Earth-history processes and lineage-specific traits, producing strong asymmetries in freshwater radiation capacity among clades.

### Why do younger Sahul freshwater fish clades diversify faster?

Among alternative time-calibrated trees and diversification-rate estimators (MoM, MiSSE), we recover a consistent inverse age-rate pattern in Sahul, with younger freshwater clades diversifying faster than older ones. Although methodological artifacts can bias such slopes, including lineage-through-time distortions (the “pull of the present”) and apparent slowdowns from protracted speciation (59), sensitivity to incomplete sampling and dating error (60), and non-identifiability of diversification histories from extant time trees (61), the pattern persists across alternative trees and rate estimators, suggesting a biological component. Notably, MoM and MiSSE rely on different features of the data, with MoM based on clade age and richness and MiSSE on full branching-time likelihoods under a birth-death process, so their concordance argues against a shared estimator artifact driving the slope. These artifacts are also not expected to produce the same directional bias: lineage-through-time effects tend to favor younger-faster signals, protracted-speciation slowdowns bias in the opposite direction, and incomplete sampling and dating error may distort rates either way depending on their distribution among clades.

Our analyses concern diversification rates rather than absolute richness. The time-for-diversification hypothesis holds that older clades accumulate more species simply by having had more time (62, 63), consistent with species-rich older groups like Melanotaeniidae. Yet several equally old Sahul transitions remain species-poor (e.g., Belonidae, Hemiramphidae, Sciaenidae, Synbranchidae), showing that time alone is insufficient. These contrasts suggest that the timing of freshwater colonization relative to transient paleoenvironmental windows, rather than absolute clade age, has been a key determinant of diversification success. Two complementary mechanisms, acting at different spatiotemporal scales, can account for this timing effect. First, long-term Australian aridification since the Eocene-Oligocene transition (∼33 Ma) progressively reduced the extent, permanence, and connectivity of inland freshwater habitats (15, 16). Early freshwater colonizers may therefore have lost substantial habitat over time, reducing diversification mainly through elevated extinction as freshwater systems became smaller and less connected at the continental scale (3, 20). At sub-drainage scale, reduced habitat heterogeneity may also have limited opportunities for niche divergence and ecological specialization among persisting lineages. Second, younger lineages diversified during later intervals when Miocene tectonic and sea-level dynamics renewed coastal connectivity and reorganized drainages (see below). In particular, these clades appear to have capitalized on Middle Miocene sea-level minima and marine low-oxygen episodes at 16–11 Ma, which enhanced marine-freshwater connectivity, promoted founder isolation during subsequent transgressions, and generated secondary ecological opportunities before incumbency constrained diversification (6, 21, 64, 65).

These results are not consistent with open-ended radiation under release from competition, as early arrival does not predict diversification outcomes. Fossil and phylogenetic evidence from the African Great Lakes show cichlids diversifying rapidly alongside diverse non-cichlid assemblages, challenging the view that absence of incumbents is a prerequisite for adaptive radiation (66). In Sahul, functional-space volume is decoupled from clade age (Fig. 4a), and late-arriving lineages fall inside the functional envelope established by earlier invaders rather than extending beyond it (Figs. 4, S26b,c). This pattern is consistent with a niche-preemption priority-effect scenario (19), in which earlier arrivers may limit the trait space available to late comers rather than promoting continued ecological expansion (67, 68). Some older lineages, such as terapontids, show lagged bursts more consistent with delayed environmental triggers (59, 69) than with an early-arrivers advantage. Habitat generalism, broad salinity tolerance, and osmoregulatory flexibility likely amplified these bursts by favoring entry and persistence in variable river and estuarine systems (43, 70, 71). An alternative, complementary explanation is that more recent colonizers were better able to adapt to freshwater conditions, facilitating establishment and diversification. Our data cannot distinguish this from time-dependent rate decline or coincidence with Middle Miocene paleoenvironmental windows, and these mechanisms may have acted together. The same marine-derived lineages that invaded Sahul rivers (ariids, eleotrids, mugilids) have also colonized other freshwater regions with depauperate primary freshwater faunas, such as Mesoamerica, suggesting that release from competition may facilitate successful colonization even when, as in Sahul, it does not lead to open-ended radiation.

Collectively, our results indicate that Sahul’s freshwater radiations follow the episodic, space-limited dynamics typical of island archipelagos and coastal river basins, where opportunity-driven pulses are governed by the timing of geographic and environmental change (72, 73). Sahul inverts the early-arrivers advantage at the level of diversification rate: younger clades diversified faster, even though their trait space fell within the envelope set by earlier invaders. Ecological opportunity was therefore contingent on lineage traits and modulated by intermittent Earth-history events, including tectonic uplift and drainage reorganization that reconfigured seaways and watershed divides, which reopened subsets of niche space to younger colonizers as older lineages settled into narrower, specialized roles (6, 59, 69). This recurrent pattern across replicated aquatic systems shows that diversification trajectories in systems like Sahul reflect the interplay of historical contingency and intrinsic clade properties.

### Radiation following freshwater colonization is episodic, lineage-specific, and often delayed

Diversification patterns among Sahul freshwater fishes indicate that radiation following marine-to-freshwater colonization is neither universal nor tightly constrained to the moment of initial invasion. Some lineages, especially ariids, show rapid accelerations in diversification immediately following freshwater entry, consistent with classic expectations of adaptive radiation in species-poor environments (11). However, many other colonizations yield few species despite long residence times, and others exhibit pronounced lags before diversification accelerates. These outcomes challenge the expectation that invasion of depauperate freshwater systems should consistently result in immediate and sustained lineage proliferation (17, 18). Instead, freshwater entry alone appears insufficient to trigger adaptive radiation, highlighting strong contingency in diversification outcomes and suggesting that ecological opportunity is realized only under particular historical and ecological conditions (35, 36).

Several well-studied clades illustrate this decoupling. Terapontids, despite being frequently highlighted for dramatic trophic diversification spanning carnivory to herbivory and accompanied by shifts in gut morphology and dentition (74), show only moderate overall diversification rates (Fig. 2b), with a pronounced burst confined to a single lineage that originated long after freshwater colonization (Fig. 3). Rainbowfishes (Melanotaeniidae) and blue-eyes (Pseudomugilidae) provide a contrasting case: despite sharing a common freshwater ancestor, their diversification dynamics differ markedly, with consistently lower rate shifts in pseudomugilids than in rainbowfishes (Fig. 2b; 28). Beyond this, in Melanotaeniidae, the strongest diversification pulses are concentrated in younger subclades associated with the Middle Miocene colonization peak (Fig. 3), consistent with independent evidence for deep regional divergence and rapid mid-Miocene diversification in New Guinea (64). These patterns suggest that radiation in Sahul freshwater fishes was not confined to initial freshwater entry but could be delayed or reactivated by secondary ecological opportunities, whether arising from environmental restructuring or interacting with lineage-specific traits.

### Why are functional breadth and lineage diversification rates decoupled?

Among Sahul freshwater fishes, functional breadth and diversification rate are weakly associated. Most clades either diversify extensively in relatively constrained trait space or occupy broad functional space without achieving high species richness, with only a minority combining both high diversification and marked functional expansion. Of these, sea catfishes (Ariidae) and grunters (Terapontidae) stand out as exceptions, having been proposed as regional adaptive radiations that couple elevated diversification with trophic shifts and morphological change at the marine-freshwater interface (11, 34, 42). Many other lineages radiate taxonomically in narrow functional bounds (e.g., eleotrids, rainbowfishes) or span broad trait space without corresponding increases in species richness (e.g., kuhliids).

The decoupling of ecological trait space and lineage diversification rates is not unexpected because these metrics capture distinct evolutionary processes: the extent of occupied ecological trait space versus the net balance of speciation and extinction. Indeed, recent syntheses emphasize that ecological opportunity is often transient and context dependent, and that diversification can decouple from expansion along broad ecological or functional trait axes (72). Even so, adaptive radiation theory often predicts coordinated ecological and taxonomic expansion under strong ecological opportunity (73, 74), raising the question of why these dimensions diverge in this region.

In Sahul, where the absence of dominant primary freshwater incumbents clearly created broad ecological opportunity for marine invaders, several non-exclusive mechanisms plausibly decouple ecological and taxonomic diversification. First, rapid lineage accumulation can proceed via geographic isolation among river drainages with limited divergence along coarse functional axes such as body size, trophic identity, or salinity breadth, yielding high diversification in compact trait space (75), as observed in Melanotaeniidae and Pseudomugilidae (7, 64). Second, habitat generalism and euryhalinity can expand functional space, thereby maintaining gene flow and limiting population fragmentation, producing species-poor but functionally expansive clades such as kuhliids and some mugilids (43, 71). Third, ecological limits and incumbency effects may favor niche packing rather than niche expansion following colonization, inflating species richness around similar optima without increasing functional volume, consistent with theoretical expectations (59, 69). Finally, substantial functional change may occur along physiological, behavioral, or fine-scale morphological axes not captured by our composite trait ordination, consistent with broad comparative evidence that phenotypic or ecological diversification often shows weak correspondence with species diversification rates (76–79). Overall, this heterogeneity reinforces the view that ecological and taxonomic diversification are shaped by partially independent processes, making decoupling the norm rather than the exception, even in systems such as Sahul where ecological opportunity is unusually pronounced.

### Paleoenvironmental modulation of colonization opportunity in Sahul

Paleoenvironmental fluctuations on macroevolutionary timescales profoundly shaped habitat connectivity and dispersal opportunity, and in the Sahul region, this dynamism has been especially pronounced. Repeated tectonic uplift, sea-level fluctuations, and drainage reorganizations throughout the Cenozoic repeatedly reconfigured coastal landscapes, creating episodic windows during which marine lineages could access, persist in, and diversify in freshwater systems (6, 21, 80, 81). Our analyses associate the prominent Middle Miocene peak in marine-to-freshwater colonization (16–11 Ma) with intervals of sea-level regression following the Miocene Climatic Optimum, coincident with expanded marine oxygen minimum zones. This pattern is consistent with independent evidence that sea-level regression enhanced marine-freshwater connectivity throughout the Sahul Shelf (3, 6, 82) and with broader biogeographic evidence from terrestrial and freshwater invertebrates, where late Miocene uplift and landscape emergence in New Guinea promoted diversification (83).

Post-optimum sea-level regressions expanded coastal and estuarine habitats of the Sahul Shelf, forging transient corridors between marine and inland systems (55, 80, 81). These conditions likely promoted founder-event colonization by euryhaline taxa, followed by isolation during subsequent transgressions, a mechanism consistent with repeated cross-shelf exchange documented among Sahul biota (84). At the same time, Middle Miocene intensification of marine hypoxia extended onto shelf and coastal settings (85), physiologically filtering for lineages tolerant of low oxygen. Because tropical rivers are themselves episodically hypoxic under high temperature and stratification (86, 87), this same tolerance would have lowered the physiological barrier to freshwater colonization, biasing entry toward hypoxia-tolerant marine lineages.

Sea-level fluctuations have similarly been invoked to explain freshwater colonization in tropical South America, where Miocene sea-level change and tectonic loading facilitated freshwater entry by marine-derived drums, pufferfishes, and other lineages (48). There, episodic marine incursions into low-lying Amazonian (and more contentiously, Paraná) drainages (49, 51) created transitional environments from the landward side via foreland subsidence, whereas on the Sahul passive margin, Miocene regression from the elevated Oligocene-Miocene Climatic Optimum baseline exposed outer shelf areas and produced shallow environments from the seaward side. In both settings, sea-level dynamics generated transitional habitats that marine-derived lineages exploited, with Sahul lowstands reopening invasion routes into otherwise depauperate rivers during the early-phase New Guinea orogenesis and drainage integration (4, 16, 60). Older colonizing lineages may have specialized into narrower roles early in their freshwater history, whereas younger clades exploited renewed windows of access created by Middle Miocene low sea level. Rather than a general rule that younger lineages diversify faster, the inverse age-rate signal we recover reflects Sahul’s specific tectonic and oceanographic history, which structured the timing and selectivity of freshwater invasions and shaped which marine lineages established and diversified in rivers.

### Conclusions

By leveraging repeated marine-to-freshwater transitions in the tropical rivers of Sahul, comprising Australia and New Guinea, we show that ecological opportunity alone does not predict diversification outcomes. Instead, freshwater radiations emerged selectively during paleoenvironmental windows that governed access, isolation, and persistence in rivers, whereas lineage-specific traits modulated which colonizers succeeded. The resulting decoupling of colonization timing, functional expansion, and diversification rate reveals that adaptive radiation is neither inevitable nor uniform following ecological release. Rather, diversification in this natural experiment reflects the interplay of Earth-history dynamics and biological traits, emphasizing that the tempo and mode of evolutionary radiations are shaped as much by when opportunity arises as by who arrives.

## Material and Methods

Extended Material and Methods are provided in the online *Supplementary Information, Extended Material and Methods*.

### Phylogenomic backbones, expanded legacy-marker trees, time calibration and subclade grafting

We assembled a genomic dataset of 825 FishLife exon markers for teleosts, combining newly sequenced species (144), exon sequences mined from publicly available genomes in NCBI (83), and exons from previously published FishLife exon-capture studies (356). We also incorporated exon data mined from UCE-based raw sequences of 22 species (see Table S1). We inferred gene trees for all loci using codon-partitioned RAxML-NG v0.9.0 analyses under the GTR+G model and generated species trees with ASTRAL-III v5.7.1. We also performed concatenation-based maximum-likelihood analyses in RAxML-NG, partitioning the concatenated alignment by codon position (3 partitions). We constructed six backbone trees using either all loci or two 416-locus subsets anchored by seven shared genes: ASTRAL Master Tree (MT), ASTRAL Subset-1 (S1), ASTRAL Subset-2 (S2), RAxML MT, RAxML S1, and RAxML S2. This design provided a distribution of alternative topologies to account for phylogenetic uncertainty in downstream comparative analyses (see also (88, 89)). After quality control (QC), the backbones comprised 578 taxa.

To broaden taxonomic coverage, we added 765 species using 12 legacy markers retrieved, mostly from NCBI and BOLD (up to 16,639 sites; Table S1), constraining placements to the phylogenomic framework via backbone-constrained analyses. An eight-gene “anchor” panel was used to bridge backbone and legacy sequence data. Constrained ML searches in RAxML-NG (via CIPRES) were run using by-codon partitions. After removal of misplacements, the expanded matrices contained 1,373 species that passed final QC (Fig. S1). The expanded trees were time-calibrated with 318 secondary calibrations (Tables S4–S6) using penalized likelihood in TreePL (90).

We then grafted densely sampled subclade phylogenies onto the expanded trees, with each subclade temporarily rescaled to match backbone divergence times. Subclade phylogenies were drawn from multiple sources, including Ariidae (34), Gobiiformes (12, 29, 91, 92), Atheriniformes (93), Clupeiformes (94), Terapontidae (42), and Syngnathiformes (95) (Table S4). Randomized grafting over replicates yielded 30 final trees comprising 2,303 taxa, representing the most comprehensive phylogenomic teleost framework assembled to date, with particular emphasis on Sahul freshwater colonizations and marine close relatives. These trees provide the foundation for all downstream analyses.

### Habitat occupancy datasets

We assembled habitat records for 2,303 species spanning marine, brackish, and freshwater environments by combining FishBase (FB) (27) and the Catalogue of Fishes (CoF) (8), supplemented with expert input for focal families (Table S7). Habitat assignments differed slightly between databases (CoF: 968 marine only, 14 brackish only, 432 freshwater only; FB: 970, 18, 453, respectively; Tables S7–S13), and a total of 389 conflicts were reconciled via our expert review. To account for euryhalinity and uncertain assignments, we combined three dataset types (raw, Reconciled Fresh+, and Reconciled Fresh Only) with two backbone sources (FB and CoF), yielding six datasets comprising 1,561–1,694 species. All marine and brackish taxa were retained, whereas freshwater species outside Sahul were pruned, consistent with Wallace’s Line as a near-impassable marine barrier for obligate-freshwater lineages. Some subclades, however, are multiregional (e.g., Anguillidae, Plotosidae, Oxudercidae; Fig. 3), spanning Wallace mostly via diadromous, euryhaline, or marine descendants (see Extended Materials and Methods).

### Ancestral habitat inferences and colonizations though time

We reconstructed ancestral habitats using BioGeoBEARS (26) based on alternative habitat datasets derived from FB and CoF. We fitted three biogeographic models, DEC (dispersal-extinction-cladogenesis), DIVALIKE (dispersal-vicariance), and BAYAREALIKE (widespread sympatry), each with and without the founder-event speciation (+J) parameter (92). Model fitting was performed on the ASTRAL and RAxML master trees (96). Across all datasets and trees, BAYAREALIKE+J was consistently favored (Akaike weight = 1 in all cases; Tables S20–S23), supporting marine-to-freshwater transitions via either anagenetic range expansion into freshwater or cladogenetic founder-event colonization. Using the best-fitting BAYAREALIKE+J model, we summarized ancestral habitat probabilities over the full set of 30 phylogenies by projecting node-wise estimates onto a master-tree topology following the approach of (28). Results shown in the main text are based on the ASTRAL master tree with the FB Reconciled Fresh+ dataset (Fig. 2a), with alternative datasets presented in Figs. S4–S9. We further conducted stochastic mapping analyses (100 maps per run) across the six alternative datasets using both ASTRAL and RAxML master trees. For the FB and CoF Reconciled Fresh+ datasets, stochastic mapping was also performed on eight supplementary trees, yielding a total of 2,000 maps (Tables S16, S24). From these maps, we quantified freshwater colonization events through time by calculating minimum, maximum, and mean numbers of transitions in 2 Ma bins, generating colonization-through-time curves (Table S16; Fig. 2b). Comparable results across datasets and trees are shown in Fig. S10. To account for phylogenetic uncertainty, estimates from all 30 phylogenies were projected onto a common master topology, and the full pipeline was repeated for raw and Fresh Only coding schemes. Because results were broadly consistent across treatments (Figs. S4–S9), we retained the Fresh+ datasets for downstream analyses. Possible biases affecting colonization-through-time estimates due to the inclusion of the ancient primary freshwater lineage Osteoglossidae in the trees are addressed in the *SI Extended Methods*.

Applying the criterion of ≥2 Sahul freshwater species, we identified and retained 17–23 independent marine-to-freshwater subclades for further analyses. Clade composition was largely consistent across datasets. Minor differences were observed in Ambassidae and Terapontidae, and several groups showed evidence of multiple independent freshwater colonizations (e.g., clupeids, gobies). For *Craterocephalus*, all species were conservatively grouped as one subclade due to incomplete phylogenetic resolution. The endemic genus *Glossamia* (Apogonidae) might represent another freshwater colonization with *in-situ* speciation, once it has been documented at least four species of this genus in Sahul estuaries and fresh waters (97). Because we only have one species (*Glossamia aprion*) in our trees, we did not retain this group for further analyses. Likewise, other families (e.g., Blenniidae, Polynemidae; see Fig. 1) are underrepresented in our trees and are not discussed further here.

### Diversification rate analyses

We retrieved stem and crown ages for each of the 23 subclades from the 30 time-calibrated trees and averaged the results over trees (Table S14; *ape*; (98)). Net diversification (speciation rate minus extinction rate) was first computed via method-of-moments (MoMs) using CoF richness (8), under different extinction schemes (ε = 0, 0.5, 0.9) using *geiger*. Although absolute values varied with ε, rank order was stable (Tables S27–S29), and ε = 0.5 was retained for downstream analyses. We also applied MiSSE, incorporating sampling fractions and fitting a single-rate class model (one turnover and one extinction fraction) to each of the 23 subclades across all 30 trees, then averaged the resulting diversification rates. MoMs and MiSSE rates were strongly correlated (R² = 0.82; Fig. S5), thus we report MoMs in the main text and MiSSE in Supplementary Information (Table S30). To localize lineage-specific diversification rate shifts, we applied ClaDS in *Julia* (99) on the RAxML-NG and ASTRAL master trees on 11 family- or order level groups (i.e. Ariidae, Plotosidae, Gobiiformes; Tables S30–S31), in which the 23 Sahul freshwater colonization subclades are nested. These analyses were based on the full set of trees (n = 2,303 species).

### Trait data and functional space

We compiled three traits: habitat breadth (salinity range: high = marine+brackish+freshwater; medium = any two; low = single habitat), body size (standard length–SL via *rfishbase*; (100)), and trophic identity using five diet categories (following (101)). We focused on fish traits linked to main ecological roles in aquatic ecosystems (102), also accounting for a broader functional definition (103). Coverage was 270/343 species across the 23 subclades (Table S15). We ran PCAmix with categorical (habitat, trophic) and continuous (log10 SL) variables using PCAmixdata (104), followed by a phylogenetic PCA (105) on the ASTRAL master tree using *phytools* to obtain phylogenetically corrected scores (Table S17). Functional occupancy per subclade was quantified as multidimensional hypervolumes using dynamic range boxes (DRB; (106)) over seven phyloPCA axes. We summarized hypervolume product (total breadth), arithmetic mean (per-dimension range), and geometric mean (central tendency). Because hypervolumes require ≥3 species, five subclades lacked estimates, leaving 18 subclades for hypervolume analyses (see Fig. 4; Tables S17–S18). We quantified functional space overlap with the Sørensen–Dice coefficient and distances as Euclidean distances between hypervolume centroids (Fig. S26b,c).

### Correlations among functional space, clade ages, and diversification rates

We tested relationships among functional space, clade age, and net diversification using OLS and PGLS, comparing OLS, PGLS-BM, and PGLS-OU (“OUrandomRoot”) and retaining the best-fitting model in each case (using *phylolm*; (107)). For PGLS, each of the 30 trees were pruned to one Sahul freshwater representative per subclade with hypervolume data (n = 18; Tables S17–S18); for OLS, the ASTRAL master tree was converted to a star phylogeny using *phytools*. To evaluate the early arrivers versus late comers hypothesis, we regressed net diversification rate (MoM) against stem and crown ages. MoM estimates were available for 22 of the 23 subclades, excluding one clupeid subclade with zero diversification. To test the second prediction, that early arrivers should occupy greater functional space, we regressed hypervolume product (Fig. 4) and geometric mean (Fig. S26) against both stem and crown ages. Finally, to test whether ecological diversification is linked to lineage proliferation, we regressed net diversification rate against hypervolume product and geometric mean (Fig. S27), predicting positive associations under adaptive radiation theory (Table S33).

### Colonizations though time and paleoclimate

From the 2,000 stochastic maps (see section above) based on CoF and FB Reconciled Fresh+ datasets, we derived colonization-through-time curves (2-Ma bins) as the response in generalized additive models (GAMs) using *mgcv::gam* (53). Predictors were eight paleoclimate variables: global average and tropical temperatures (108), sea level (58), Australian inundation (relative to the present-day 200 m isobath; (109)), Australian precipitation (110), salinity (111), and geochemical proxies ⁸⁷Sr/⁸⁶Sr (continental weathering) and δ³⁴S (marine redox state; (54)). Because evolutionary responses can lag environmental change (75), we fit univariate and multivariate GAMs with no lag and with 2, 4, 6 Ma lags (predictors shifted backward relative to colonization rate), and controlled for multiple testing using Benjamini-Hochberg false-discovery-rate procedure in *p.adjust* (argument ‘method=fdr’; (112)(Fig. 5; Table S19).

## Acknowledgments

Bioinformatic analyses were performed at the University of Oklahoma Supercomputing Center for Education & Research (OSCER).

