## Supplementary Information for "REPEATED MARINE-TO-FRESHWATER FISH TRANSITIONS REVEAL PALEOENVIRONMENTAL MODULATION OF ADAPTIVE RADIATION"

[1]Scripps Institution of Oceanography, University of California San Diego, 8622 Kennel Way, La Jolla, CA 92037, USA. [2]Department of Biology, The University of Oklahoma, Norman, OK 73019, USA. [3]Centre for Tropical Water and Aquatic Ecosystem Research, School of Marine and Tropical Biology, James Cook University, Townsville QLD 4811, Australia. [4]Department of Biological Sciences, The George Washington University, Washington, DC 20052, USA. [5]Vertebrate Zoology, Santa Barbara Museum of Natural History, 2559 Puesta del Sol, Santa Barbara, CA 93105, USA. [6]Department of Ichthyology, Natural History Museum of Los Angeles County, 900 Exposition Blvd, Los Angeles, CA 90007, USA. [7]Great Lakes Fishery Commission, 2200 Commonwealth Blvd, Suite 100, Ann Arbor, MI 48105, USA. [8]Bell Museum of Natural History, University of Minnesota, 100 Ecology, 1987 Buford Circle, Saint Paul, MN 55108, USA. [9]Department of Biological Sciences, Western Michigan University, Kalamazoo, MI 49008. [10]The University of Papua New Guinea University, 134 National Capital District Port Moresby, Papua New Guinea. [11]Department of Marine, Earth, and Atmospheric Sciences, North Carolina State University, 2800 Faucette Boulevard, Raleigh, NC 27607, USA. [12]North Carolina Museum of Natural Sciences, 11 W Jones Street, Raleigh, NC 27601, USA. [13]School of the Environment, Geography, and Sustainability, Western Michigan University, 1903 W Michigan Ave, Kalamazoo, MI 49008.

[*]These authors contributed equally to this work

Classification: Biological Sciences, Evolution

### EXTENDED MATERIALS AND METHODS

#### Phylogenomic backbones, expanded legacy-marker trees, time calibration and subclade grafting

*Newly and previously sequenced exon markers; genome mining*

Genomic DNA was extracted from ~20 mg of muscle or fin tissue using the DNeasy Blood & Tissue Kit (QIAGEN). DNA concentration was measured with a Qubit fluorometer (Thermo Fisher Scientific). For exon-capture sequencing, libraries were prepared by Daicel Arbor Biosciences (Ann Arbor, MI) following a TruSeq-style protocol. We used the following 'FishLife' probe sets focusing on Teleost: Elopomorpha (7 species); early-branching teleosts from Osteoglossomorpha to Myctophiformes (30 species; "Backbone 1"); Acanthomorphata to Anabantaria (89; "Backbone 2"); Carangaria (14); Ovalentaria (76); Eupercaria (73); Syngnatharia–Pelagiaria (13); and Otophysa (31) (1), synthesized using myBaits custom probes. Sequencing was performed on an Illumina HiSeq 2500, yielding 150 bp paired-end reads. High-quality DNA was also submitted to GENEWIZ (Azenta Life Sciences, South Plainfield, NJ) for whole-genome short-read sequencing. Libraries were prepared with the Illumina TruSeq Library Construction Kit (target insert ~350 bp) and sequenced on an Illumina NovaSeq S4 to ~30× coverage, generating 150 bp paired-end reads.

Illumina raw reads were quality assessed using FastQC v0.11.5 (2) and trimmed with Trimmomatic v0.38 (3) to remove adapters and low-quality bases. Filtered reads were assembled with SPAdes v3.13.1 (4). Putative contaminants were identified with NCBI's Foreign Contamination Screen (FCS) and removed prior to downstream analyses (available at https://github.com/ncbi/fcs).

Raw reads were processed with the FishLifeExonCapture pipeline (https://github.com/lilychughes/FishLifeExonCapture). Reads were trimmed with Trimmomatic to remove adapters and low-quality bases, then mapped to a representative bait set for the target clades using BWA v0.7.17 (5). PCR duplicates were removed with SAMtools v1.9 (6). Initial contig assemblies were generated with Velvet v1.2.07 (7); for each locus, the longest contig was retained as a seed/reference for iterative refinement with aTRAM v2.29 (8) using up to five iterations. Redundant contigs were collapsed with CD-HIT v4.8.1 (9) at 98% similarity.

We assembled exon data for 605 taxa by combining newly generated sequences with previously published datasets. Newly sequenced exon-capture data were generated for 144 species, and exon sequences for an additional 83 species were mined from publicly available whole-genome assemblies in NCBI. The remaining taxa were obtained from previously published FishLife exon-capture studies, including 303 species from (10), 49 species from (11), 2 species from (12), 2 species from (13). To increase representation of terapontoid lineages, we additionally incorporated data from UCE-based raw sequences of 22 species (see Table S1), from which 4–118 exon markers per species (mean = 23 exons) were mined from raw data using the same pipeline to ensure compatibility with the backbone dataset. Sequencing for those samples followed protocols outlined in (14). All exon sequences were harvested using the FishLife exon-mining pipeline (https://github.com/lilychughes/FishLifeExonHarvesting). Accession numbers and data sources for all exon and legacy markers are provided in Table S1.

*Alignment*
Reading frames were identified by aligning contigs to a curated reference using Exonerate's coding2genome model (v2.4.0; (15)), and exon alignments were produced with the frame-aware aligner MACSE (v2.03; (16)). Profile HMMs were built for each target locus and used to query each genome with 'nhmmer' (HMMER v3.2.1; (17)) under default parameters. Significant hits for each marker were retrieved across species, and the corresponding genomic intervals were extracted with custom Python scripts (https://github.com/lilychughes/FishLifeExonHarvesting). After standard quality-control and assembly steps (see above), coding sequences were aligned with MACSE (v2.03). All alignments were visually inspected in Geneious Prime (v2022.1; (18)). Individual exon alignments were then concatenated into a supermatrix of 825 loci across 605 taxa (333,381 bp), with 47.9% of missing data, which we used to infer the backbone phylogeny.

*Phylogenomic backbones*
To address phylogenetic and divergence time uncertainty, we used a supermatrix of 825 FishLife exons across 605 taxa (333,381 bp) to infer our six phylogenomic backbones. Our comparative analyses used two phylogenetic frameworks: maximum-likelihood (ML) and coalescent-based methods. For the ML tree, we used RAxML-NG v0.9.0 (19) with a codon-partitioned scheme (Codon 1 = 1–333381\3; Codon 2 = 2–333381\3; Codon 3 = 3–333381\3) under a GTR+G model. Bootstrap support was estimated with autoMRE, and the search was conducted under a backbone constraint corresponding to the bony-fish phylogeny of (20). Gene trees for all 825 loci were inferred with RAxML-NG from codon-partitioned alignments. Branches with bootstrap support <70% were contracted to polytomies. The resulting gene trees were then used for coalescent species-tree inference with ASTRAL-III (v5.7.1; (21)). To account for phylogenetic and divergence-time uncertainty, we also constructed two partially overlapping subsets of the expanded matrix (416 loci each; seven loci overlapped: E0448, E0936, E1032, E1090, E1102, E1460, E1491; Supplementary Table S3). After multiple rounds of quality control, we retained six phylogenomic backbone trees comprising 578 of the original 605 taxa (Fig. S1): ASTRAL Master Tree (MT), ASTRAL Subset-1 (S1), ASTRAL Subset-2 (S2), RAxML MT, RAxML S1, and RAxML S2.

Next, we expanded our taxonomic sampling by incorporating 765 additional species with 12 legacy, mitochondrial and nuclear markers retrieved from previous studies (e.g. (20)), and public repositories (e.g. NCBI and BOLD Systems V3; (22): 12S (n= 276 sequences mined), 16S (n= 502), 18S (n= 115), ATP6 (n= 72), COI (n= 1290), CYTB (n= 405), MYH6 (n= 117), ND1 (n= 175), ND2 (n= 129), RAG1 (n= 213), RAG2 (n= 150), and RH1 (n= 214). Additionally, we incorporated data from an additional species in the UCE-based phylogenomic (23), from which 7 exon markers were mined from raw data using the same pipeline described above to ensure compatibility with the dataset. We provide accession numbers and sources of markers in Table S1.

*Phylogenetic inference, tree uncertainty, and subclade grafting*
After aligning our exons and legacy markers, we ran constrained maximum-likelihood (ML) searches in RAxML-NG on the CIPRES Science Gateway using codon-partitioned analyses (available in supplementary files). To identify runtime-efficient strategies, we compared two supermatrices analyzed under the master-tree constraint and evaluated concordance in recovered topologies and branch-length estimates. The first dataset comprised the full panel of 825 exon markers plus legacy loci (347,071 sites). The second used an eight-gene "anchor" panel (E0448, E0936, E1032, E1090, E1102, E1460, E1473, E1491) together with the legacy loci (16,639 bp). This reduced panel was designed to closely approximate the full-matrix results while substantially reducing computational cost, and still preserving relationships imposed by the phylogenomic backbone.

We compared and aligned the two trees using a tanglegram-style cophylogenetic plot in *phytools* (24). This visualization highlighted congruent and conflicting relationships; we then pruned taxa showing topological conflicts. Next, we regressed matched branch lengths between the RAxML-NG master tree inferred from Legacy + Anchor genes and the RAxML-NG master tree inferred from Legacy + 825 exon markers. To reduce computational cost, we proceeded with the phylogenomically constrained Legacy + Anchor panel, which closely approximated results from the full 825-exon matrix. Finally, we conducted five additional constrained ML searches corresponding to the remaining backbone trees in RAxML-NG on the CIPRES Science Gateway using codon-partitioned data.

We carried out several rounds of quality control to identify and remove misplaced or misidentified species. This process involved examining the resulting trees for unexpected placements and comparing them to established phylogenetic relationships. After removing misplaced taxa, the final matrix included 1,373 species, comprising 834 fish species from the Australia-New Guinea adaptive radiation and the broader Indo-Pacific region, along with species from the teleost backbone tree (Table S1).

We time-calibrated the maximum-likelihood and coalescent trees with treePL v1.0 (25) under a penalized-likelihood framework. A total of 318 secondary calibrations (Tables S4–S5) were retrieved from multiple time-calibrated teleost trees, including large-scale phylogenies (10, 20, 26–28) and subclade-focused (11, 12, 29) (Table S5), via congruification using 'congruify' in the R package geiger (30). Resulting node ages were evaluated against published estimates to identify and remove outlier estimates (Table S6).

We then grafted densely sampled subclades from published phylogenies onto our time-calibrated backbone using a custom R workflow (packages *ape*; (31); and *phytools*; (24); script Code_Rescale&Graft.R). Focal subclades included Ariidae (12), Atheriniformes (32), Clupeiformes (11), Terapontidae (33), Syngnathiformes (29), and Gobiiformes (34–37), with nested sets for *Hypseleotris*, *Gobionellus*, Butidae, and *Mugilogobius* (Table S2). Subclades were pruned to their designated roots and attached at the corresponding nodes in the backbone trees. Because branch-length scales differed between backbone and source chronologies, we rescaled subclades via congruification, adjusting crown or stem depths to match backbone ages. We applied this rescaling where temporal mismatches were evident (e.g., Ariidae, Atheriniformes, Gobionellidae) to ensure comparable timescales across grafts.

For Gobiiformes, we used the best-scoring maximum-likelihood tree with bootstrap support from (34) as the backbone. We calibrated this backbone with secondary calibrations from (37) using treePL. The same calibrations were applied to 10 trees from *Hypseleotris* (36) and *Gobiomorphus* (35). We then grafted the Butidae and *Mogurnda* clades (37), along with *Hypseleotris* (36) and *Gobiomorphus* (35), onto this backbone. For Clupeiformes, we recalibrated the root to match the age reported by (20), because the root age in the tree of (11) was older than expected and appeared to be an outlier. After grafting all subclades using randomizations across replicates, we obtained a final set of 30 phylogenetic trees comprising 2,303 tips. We then examined these trees for topological consistency and potential conflicts between grafted subclades and the overall structure.

**Habitat occupancy datasets**

For habitat occupancy, we assigned 2,303 species to geographic units defined by ocean basins and continents (Table S7). We assembled a presence-absence matrix across three habitat categories (marine, brackish, and freshwater) by aggregating records from FishBase (FB; (38)) and the Catalogue of Fishes (CoF; (39)), supplemented with our expert knowledge of Ariidae, Gobiidae, Eleotridae, Oxudercidae, Butidae, Clupeiformes, Atheriniformes, and Terapontidae. Based on CoF database, 968 species were assigned as marine only, 14 as brackish only, 348 as marine and brackish, 358 as marine, brackish and freshwater, 182 as brackish and freshwater, and as 432 freshwater only species (Table S8). In FB, the corresponding values were 970 marine only species; 18 brackish only; 453 freshwater only; 351 marine and brackish; 188 brackish and freshwater; 323 marine, brackish and freshwater (Table S9). Overall, 389 species showed habitat-assignment discrepancies between FB and CoF. Where the databases disagreed, expert input was used to reconcile assignments, resolving records for 133-140 species (Tables S10–S11). To maximize detection of marine or brackish sister lineages to Sahul freshwater transitions, we retained all marine and brackish species for downstream analyses but pruned freshwater species occurring outside Australia and New Guinea, retaining only Sahul freshwater taxa.

Many species occupy broad salinity gradients, either during particular life stages (ontogenetic shifts; (40, 41)) or throughout the life cycle, which often warrants classification as euryhaline for taxa recorded in more than one habitat (e.g., marine, brackish, and freshwater; (42)). For example, anguillids, one of the clades inhabiting Sahul freshwaters, are typically catadromous, migrating downstream to spawn at sea, and occur across marine, brackish, and freshwater environments (43). To accommodate habitat variation, we conducted ancestral-range reconstructions under three coding schemes: (i) raw FB and CoF datasets; (ii) “Fresh+”, retaining freshwater-designated species that also have marine and/or brackish records; and (iii) “Fresh Only”, retaining freshwater-designated species without marine records (i.e., recorded only in brackish and freshwater). Accordingly, we assembled six habitat datasets: (i) FB raw (n = 1,665; Table S9) and CoF raw (n = 1,675; Table S8); (ii) FB Fresh+ (n = 1,677; Table S11) and CoF Fresh+ (n = 1,694; Table S10); and (iii) FB Fresh Only (n = 1,561; Table S13) and CoF Fresh Only (n = 1,578; Table S12). Note that species counts differ between FB and CoF because each database designates some freshwater species outside our focal area (Sahul) differently.

The geographic restriction of our analyses to Sahul, east of Wallace's Line, reflects both the empirical scope of the dataset and the biogeographic improbability of shared freshwater transitions crossing this boundary. Wallace's Line marks one of the sharpest faunal discontinuities on Earth, having limited marsupial-placental and other faunal exchange between Asia and Australia for tens of millions of years (44, 45); for obligate freshwater fishes, the intervening deep-water marine barrier is, if anything, even more impassable. We verified this expectation by manual inspection of the six densely sampled subclade trees used here (Ariidae, Atheriniformes, Clupeiformes, Gobiaria, Terapontidae, and Plotosidae). Every freshwater clade with a marine sister lineage is restricted to one side of Wallace's Line, with a single exception: *Leiopotherapon plumbeus* (Terapontidae), an obligate freshwater endemic from the Philippines (Luzon: Laguna de Bay and Taal Lake; (46)), nested as sister to Australian *Amniataba* in the Sahul terapontid radiation (47). This topology implies a single over-water colonization by a terapontid ancestor, rather than two independent marine-to-freshwater transitions, and therefore does not inflate our Sahul marine-to-freshwater counts. Secondary freshwater-to-marine reversals in Sahul ariids (12) likewise remain confined to the Australian-New Guinean shelf.

Several diadromous or euryhaline lineages have broader Indo-Pacific ranges that span Wallace's Line (e.g., sicydiine and oxudercine gobies (48); anguillid eels (49, 50); *Megalops cyprinoides* (51); *Kuhlia* spp. (52); *Mesopristes* spp. (47); catadromous and euryhaline mugilids including *Mugil cephalus* s.l., *Aldrichetta forsteri*, and *Trachystoma petardi* (53, 54). These taxa are coded as euryhaline rather than obligate-freshwater and therefore contribute to eury-state transitions only; they do not generate shared cross-Wallace marine-to-freshwater events in our counts. The absence of cross-Wallace obligate-freshwater clades in the subclade trees, combined with the Sahul-restricted pruning of the master tree, supports our interpretation that the transition counts reported here reflect transitions endemic to Sahul, consistent with Wallace's Line (sensu Huxley) acting as a near-impassable marine barrier for obligate-freshwater lineages.

**Biogeographic model selection and ancestral habitat inferences**

We performed model selection for Sahul freshwater fishes using the six habitat datasets described above and both ASTRAL and RAxML Master Trees (MTs), which correspond to the trees assembled with the full exon matrix. In addition to the MTs, we also performed this analysis for eight more of the 30 trees derived from Fresh+ datasets (Tables S20–S23), for a total of ten trees. Model selection was performed in a geographic framework implemented in BioGeoBEARS (55), which allows species to occupy two habitats simultaneously rather than forcing instantaneous habitat transitions, as is typically the case with standard Markov models (56, 57). We fit three BioGeoBEARS models: DEC (56, 57), the likelihood version of DIVA (DIVALIKE; (58)), and BAYAREALIKE (59); each with and without the founder-event speciation ("+J," jump-dispersal) parameter (55). Although there is ongoing debate about whether +J inflates likelihoods and biases model choice (60), the best-fitting model across all biogeographical analyses was BAYAREALIKE+J (Tables S20–S23). This result suggests that a marine lineage can give rise to a freshwater lineage via either anagenetic range expansion into freshwater or cladogenetic founder-event speciation (55). Under the anagenetic interpretation, a lineage broadens its salinity tolerance to include freshwater ("freshwater colonization") while retaining its marine range; under the cladogenetic '+J' interpretation, a founder population shifts into freshwater and speciates without retaining the marine range.

To assess the effects of phylogenetic variation on biogeographic inference, we used a script produced by (61) to summarize ancestral-range estimates across trees by adopting the "master-tree" topology as a reference and overlaying the probability estimates from all 30 trees onto it. We applied the same pipeline to the six dataset schemes (FB and CoF raw data; FB and CoF Fresh+; FB and CoF Fresh Only; Figs. 1a, S4–S9). Additionally, based on the best-fitting model, we generated 100 stochastic maps per dataset to assess the number of freshwater colonizations during the last 56 Ma (Tables S16, S24; Figs. S10–S11), yielding a total of 2,000 stochastic maps across the reconciled Fresh+ datasets (10 distinct trees, including RAxML and ASTRAL master trees, multiplied by two coding schemes). For each stochastic map, we generated a table with states at each node and tip following Xing and Ree (2017)'s approach. We set 2 Ma bin and used a function to count colonization events (minimum, maximum and mean rate value) at every 2 Ma of lineages arriving to freshwater. The reconciled Fresh+ and Fresh Only schemes yielded similar results (Figs. S10–S11), with freshwater colonization peaks centered in the Middle Miocene (16–11 Ma) and a weaker but consistent peak during the Oligocene (30–25 Ma). Across both schemes, the same major subclades consistently showed freshwater colonization events, including Ariidae, Plotosidae, Clupeidae, Gobiiformes, Terapontidae, Synbranchidae, Toxotidae, Ambassidae, Atherinidae, and Pseudomugilidae + Melanotaeniidae (Figs. S8–S11). Based on this concordance, we retained the FB and CoF reconciled Fresh+ datasets for all downstream analyses. Phylogenies with ancestral range estimates from the FB and CoF reconciled Fresh+ datasets were then visually inspected to identify marine-to-freshwater transitions and to delineate subclades associated with in situ diversification, defined as those containing two or more Sahul freshwater species. Each subclade was delimited at the first node exhibiting a high likelihood of freshwater occupancy.

A caveat of our ancestral range reconstructions and colonization through time analyses concerns bonytongues (Osteoglossidae), which are represented in Sahul by two extant freshwater species (39, 63) and therefore meet our initial inclusion criteria. However, osteoglossids are widely regarded as an ancient freshwater lineage whose transition from marine environments occurred deep in the Mesozoic, well before the Cenozoic interval that is the focus of our analyses. Although marine and marginal-marine fossils have been documented for early osteoglossomorphs, all extant osteoglossids are obligate freshwater fishes with a long-standing freshwater evolutionary history (64, 65). This deep antiquity, combined with incomplete sampling of non-Sahul freshwater osteoglossids in our phylogenomic framework, likely leads to substantial underestimation of the timing of their marine-to-freshwater transition in our reconstructions. Because our study targets repeated, Cenozoic marine-derived freshwater invasions followed by *in situ* diversification, we therefore exclude Osteoglossidae from downstream analyses of diversification rates and functional trait space, treating them as relict primary freshwater fishes rather than as part of the replicated marine-to-freshwater transitions central to this study.

**Identification of subclades associated with marine-to-freshwater transitions**
Based on the criteria defined above (i.e., subclades with ≥2 Sahul freshwater species), we identified 14 subclades associated with freshwater colonization and subsequent diversification in CoF Fresh+ dataset, and 23 subclades in FB Fresh+ dataset. This difference was mostly due to node placement of first freshwater colonization in Gobiiformes (Figs. S8–S9). CoF Fresh+ habitat reconstruction shows a freshwater colonization around 55–60 Ma for Gobiaria, which includes Eleotridae (Fig. S8). However, there is a reversal to marine habitat around 30 Ma, marking the marine ancestry for Butidae, Oxudercidae and Gobiidae (Fig. S8). Butidae and Gobiidae each show a single freshwater colonization for the family, whereas Oxudercidae exhibit multiple independent freshwater colonizations. FB Fresh+ habitat reconstruction depicts a marine ancestry for Gobiaria, with multiple independent freshwater colonization events in the last 20 Ma within the families (Fig. S9), a result concordant with a recent study on diversification of sleepers (37). For consistency, and following the literature, we retained Gobiaria subclades based on FB Fresh+ freshwater colonization events.

The remaining clades were consistent across both datasets, with only Ambassidae and Terapontidae showing differences: in Ambassidae, the marine-derived freshwater subclade comprised 18 species in CoF+ but 16 in FB+ because *Ambassis ambassis* and *A. kopsii* were coded as marine+brackish in CoF versus marine+brackish+freshwater in FB (Tables S10–S11); in Terapontidae, *Rhynchopelates oxyrhynchus* was coded as marine-only in CoF but as freshwater in FB (Tables S10–S11). Clupeidae, Eleotridae, and Oxudercidae (Table S25) each exhibited multiple independent freshwater colonization events, and we treated each event as a separate subclade. By contrast, Pseudomugilidae and Melanotaeniidae represent a single colonization (see Unmack and Dowling 2010; Campanella et al. 2015) and were therefore grouped into one subclade. Within Atherinidae, the genus *Craterocephalus* shows two independent freshwater colonizations, consistent with previous studies (66, 67) (see Fig. 2a). For atherinids, we nevertheless treated all *Craterocephalus* species as a single subclade for downstream analyses (e.g., net diversification, hypervolume) because, to our knowledge, no comprehensive phylogeny presently resolves the placement of species absent from our dataset and therefore we cannot confidently assign those missing species to one colonization event versus the other.

**Clade age, diversification dynamics, and functional space**

The 23 retained subclades comprised 343 species. To test relationships among clade age, diversification rate, and functional space we extracted stem and crown ages for each subclade, estimated net diversification rates using two complementary approaches, and quantified functional space using trait-based hypervolumes (68).

*Stem and crown ages*

We first pruned each of the 30 time-calibrated trees to the focal species in a given subclade plus a single close relative (outgroup). For each pruned tree, we extracted node heights with and without the outgroup to obtain stem and crown ages, respectively, using functions in the R package *ape* (31). Stem and crown ages were computed for all subclades using the CoF Fresh+ dataset (Table S14). Because only two subclades, Ambassidae and Terapontidae, differed between CoF and FB (Table S25), we also calculated ages for those under the FB Fresh+ coding (Table S14); the resulting clade age estimates were consistent with those from CoF Fresh+. Stem and crown ages were then averaged across the 30 trees to obtain a single estimate per subclade for downstream analyses.

*Net diversification rates*

We first estimated net diversification rates for the 23 subclades identified as marine to freshwater transitions using the method-of-moments (MoM) estimator (69). This estimator requires only clade age and species richness, which we compiled for all subclades; richness counts followed the Catalogue of Fishes (39). Previous studies indicate that MoM performs well across a range of diversification scenarios (70–72), and rates can be compared under alternative relative-extinction fractions. Accordingly, we estimated rates under three extinction schemes ($\varepsilon = 0$, $\varepsilon = 0.5$, $\varepsilon = 0.9$) following best practices (72–74). MoM rates were computed using 'bd.ms' function in the R package *geiger* (30). Although absolute values varied across ε, the relative ranking of subclades was consistent across all three schemes (Tables S26–S28), and we used the moderate-extinction setting ($\varepsilon = 0.5$; Table S27) for downstream analyses.

We also estimated net diversification rates with MiSSE (Missing-State Speciation and Extinction), a likelihood framework implemented in the R package *hisse* (75) that models diversification while accommodating hidden (unobserved) states (76). We applied sampling fractions and fit a single rate-class MiSSE model, with one turnover and one extinction fraction, to each of the 23 subclades across all 30 trees. We then averaged diversification rates across the 30 trees. Net diversification estimates from MoM and MiSSE were strongly correlated ($R^2 = 0.82$; Fig. S5); therefore, we report MoM estimates in the main text and provide MiSSE results here (Table S29). Both approaches yielded concordant estimates of overall net diversification rates.

*Diversification rate shifts*

We also examined diversification-rate shifts within subclades to test whether elevated rates occurred immediately after freshwater colonization or arose later in their history. We traced lineage-specific dynamics using the Cladogenetic Diversification Rate Shift

(ClaDS) model, which infers frequent, small-magnitude rate changes along lineages (77). ClaDS models were fit in *Julia* (78). In ClaDS, each lineage has its own speciation rate; at each speciation event, daughter lineages receive new rates drawn from a distribution parameterized by the parent rate (77). We ran ClaDS on the two MTs (RAxML-NG and ASTRAL) accounting for all species (n = 2,303 species) to include outgroups outside our focal area. We selected monophyletic groups encompassing our 23 focal clades to run this analysis. Because our taxonomic sampling is biased towards marine derived freshwater fishes, we adjusted the sampling fractions on a clade-by-clade basis (Tables S30–S31; Figs. S15–S25). ClaDS was ran with the default settings: 1,000 iterations, 0.25 of burnin, and 10 samples from the posterior distribution of complete phylogenies outputed. ClaDS results were imported to R and plotted using packages ape (31), coda (79), fields (80), and geiger (30).

*Ecomorphological data and functional space*
To quantify functional space for each subclade, we compiled three functional traits: habitat breadth, body size, and trophic identity. We focused on fish functional traits linked to their main ecological roles in aquatic ecosystems (81, 82) and, under a broader functional definition reflect organismal performance and evolutionary differentiation (see (83)). Body size and trophic identity are well-established axes in functional ecology and fish macroecology (47, 84–86), and are tightly linked to ecosystem processes such as food-web structure and nutrient cycling.

Habitat breadth was coded based on salinity range across marine, brackish, and freshwater environments: species occurring in all three were classified as high, those in any two as medium, and those restricted to a single environment as low. This trait reflects physiological tolerance and dispersal potential across salinity gradients. Body size was measured as standard length (SL) obtained from FishBase using the rfishbase package (87). Trophic identity followed the framework of (88), with species assigned to five dietary categories based on primary literature sources (Table S15): herbivores/detritivores (HD), mobile invertivores (MI), generalized carnivores (GC), omnivores (OM), and planktivores (PK).

Altogether, we retrieved ecomorphological data for 270 of the 343 species encompassed by the 23 subclades (Table S15). To quantify functional space, we first ran a principal component analysis for mixed data (PCAmix) using habitat breadth and trophic identity as categorical variables and $\log_{10}$-transformed body size as a continuous variable (R package PCAmixdata; (89). PCAmix scores (Table S17), together with the ASTRAL master tree, were then used to compute a phylogenetic PCA under a Brownian motion model with ‘phyl.pca’ in *phytools* (24), yielding phylogenetically corrected scores for our ecomorphological dataset. To quantify functional occupancy by subclades, we estimated multidimensional hypervolumes with the dynamic range boxes (DRB) estimator (68), using all seven PC-axis scores from the phyloPCA. The hypervolume metrics summarized were the hypervolume product (total multidimensional breadth), the arithmetic mean (average per-dimension range), and the geometric mean (central tendency of per-dimension ranges). Because hypervolume metrics require at least three data points, we could not compute hypervolumes for five subclades (“Clu_1”, “Oxu_2”,

"Oxu_3", "Tox", "Zen"), leaving 18 subclades for analysis (Table S18). Pairwise hypervolume overlap in the phyloPCA space was quantified with the Sørensen–Dice coefficient, which corresponds to the ratio size of the intersection to the sum of individual hypervolumes for two distinct subclades (Fig. S26b). Distances between hypervolumes were measured as the Euclidean distance between hypervolume centroids (Fig. S26c).

*Correlations between clade ages, diversification rates, and functional space*
We tested relationships among diversification rate, functional space, and clade age using ordinary least squares (OLS; no phylogenetic correction) and phylogenetic generalized least squares (PGLS). We compared three models—OLS, PGLS under Brownian motion (PGLS-BM), and PGLS under an Ornstein–Uhlenbeck process (PGLS-OU; model = "OUrandomRoot")—and retained the best-fitting model in each case. PGLS analyses were conducted with *phylolm* (90) after pruning each tree to a single Sahul freshwater representative per subclade. For OLS, we analyzed the same data without phylogenetic structure by converting the ASTRAL MT to a star phylogeny using *phytools::collapse.to.star* (24).

To test the first prediction of the early-arrivals/late-arrivals hypothesis (i.e., that older clades exhibit higher diversification) we regressed net diversification rates (MoM; $\varepsilon = 0.5$) against stem (Fig. 2c) and crown ages (Fig. S14d). We analyzed 22 of 23 subclades, excluding one Clupeidae subclade ("Clu_1"; Tables S27, S32) whose MoMs estimate was zero. Guided by adaptive radiation and ecological opportunity theory (91–93), we also predicted that higher diversification would be associated with larger functional hypervolumes. Accordingly, we (i) regressed functional space (hypervolume product and geometric mean) on clade age (Figs. 4, S27) and (ii) regressed functional space on diversification rate (MoM; $\varepsilon = 0.5$; Fig. S28). These tests used the 18 subclades with functional-space data (Table S18).

**Generalized additive models linking freshwater colonizations to environmental change over the last 56 million years**
We tested whether the mean number of Sahul freshwater colonization events over the last 56 million years (Ma) covaried with paleoclimatic variables using generalized additive models (GAMs), fit with *mgcv::gam* (94) in R. We analyzed a subset of 10 of the 30 trees, five ASTRAL (including the MT) and five RAxML-NG (including the MT) trees, and performed habitat-range reconstructions (see Biogeographic models) using the CoF and FB reconciled Fresh+ datasets. This yielded 20 independent stochastic-mappings: 10 runs using the CoF Fresh+ database and 10 using the FB Fresh+ database. For each run, we generated 100 stochastic maps and summarized the mean number of freshwater colonizations in 2-Ma time bins, producing colonization-through-time curves that were then used as the response variable in the GAM analyses (Table S16).

In the next step, we retrieved paleoclimatic data from eight environmental variables: Global Average Temperature (GAT) and Tropical Temperature from (95), Sea Level fluctuations (96), Inundation in Australia, which was determined by the flooded continental area relative to the present-day 200m isobath (97), Precipitation in Australia (98), Salinity

(99), and Strontium Isotope rate ($^{87}Sr/^{86}Sr$) and Delta-34 Sulfur ($\delta^{34}S$) (100). Strontium Isotope rate can be used as a proxy for continental weathering (the ratio of $^{87}Sr/^{86}Sr$ in seawater reflects the balance between continental weathering and mid-ocean ridge basalt alteration) indicating changes in continental weathering intensity over geological time scales. Delta-34 Sulfur by contrast, reflects the marine redox state: high values typically indicate increased pyrite ($FeS_2$) burial under reducing (anoxic) conditions, whereas lower values suggest more oxygenated seawater (100). The role of aridification is often accessed by taking into account other proxies, such as precipitation rates (e.g. precipitation/year (98)), a combination of precipitation, evapotranspiration and potential evapotranspiration (101), or even the role of hydrological and coastal topographic changes (102). Here, we used precipitation as a proxy for aridification and hydrologic connectivity and, by analyzing our paleoenvironmental variables into different ways (i.e. univariate and multivariate models; see below), we leveraged the role of each variable, alone and combined, into the freshwater colonization dynamics.

Evolutionary responses to environmental changes may occur immediately or after a delay (103). Accordingly, we fit GAMs under four designs: (i) univariate models (single predictor) with no time lag in response (see Fig. S29); (ii) multivariate models (all predictors jointly) with no lag; and (iii) univariate and (iv) multivariate models permitting lags of 2 Ma, 4 Ma, and 6 Ma. Lags were implemented by shifting predictor time series backward relative to the response (colonization rate). To control for multiple testing, we adjusted p values using the Benjamini–Hochberg false-discovery-rate procedure (*p.adjust* function, argument ‘method=fdr’; (104). Adjusted p-values (Table S19) were then used to produce boxplots representing the significance of each environmental variable under different GAMs (Fig. 5).

### EXTENDED TABLES

Tables S1–S19 correspond to raw data, datasets, and results with more than 1,000 rows, and therefore are available in the Supplementary Excel file. Here, we show supplementary tables S20–S32.

**Table S20.** Model-fitting results from BioGeoBEARS based on ASTRAL Master Tree using six dataset schemes: Catalogue of Fishes (CoF) and Fishbase (FB) raw data, CoF and FB reconciled data accounting for freshwater species with broad habitat range ('Fresh+' datasets), and CoF and FB reconciled data accounting for freshwater species with narrow habitat range ('Fresh Only' datasets). Biogeographic models include dispersal-extinction-cladogenesis (DEC), a likelihood version of dispersal-vicariance (DIVALIKE), and a likelihood of the Bayarea (BAYAREALIKE) model. All three models were inferred with and without the jump-dispersal (J) parameter. LnL: Log-likelihood; n: number of parameters estimated for each model; d: dispersal rate; e: extinction rate; w: dispersal matrix power exponential (set 1 as default); AICc: Akaike Information Criterion; AICc_wt: Akaike weight.

| TREE - ASTRAL MASTER TREE | | | | | | | | |
|---|---|---|---|---|---|---|---|---|
| COF RAW DATA | | | | | | | | |
| Model | LnL | n | d | e | j | w | AICc | AICc_wt |
| DEC | -2078 | 2 | 0.01 | 1.00E-12 | 0 | 1 | 4159 | 4.70E-123 |
| DEC+J | -2078 | 3 | 0.01 | 1.00E-12 | 1.00E-05 | 1 | 4161 | 1.70E-123 |
| DIVALIKE | -2251 | 2 | 0.011 | 1.00E-12 | 0 | 1 | 4505 | 3.70E-198 |
| DIVALIKE+J | -2251 | 3 | 0.011 | 1.00E-12 | 1.00E-05 | 1 | 4507 | 1.30E-198 |
| BAYAREALIKE | -1815 | 2 | 0.006 | 0.0044 | 0 | 1 | 3633 | 8.60E-09 |
| BAYAREALIKE+J | -1795 | 3 | 0.006 | 0.0029 | 0.0041 | 1 | 3596 | 1 |
| FB RAW DATA | | | | | | | | |
| Model | LnL | n | d | e | j | w | AICc | AICc_wt |
| DEC | -2058 | 2 | 0.011 | 1.00E-12 | 0 | 1 | 4120 | 2.90E-110 |
| DEC+J | -2058 | 3 | 0.011 | 1.00E-12 | 1.00E-05 | 1 | 4122 | 1.00E-110 |
| DIVALIKE | -2222 | 2 | 0.011 | 1.00E-12 | 0 | 1 | 4448 | 1.80E-181 |
| DIVALIKE+J | -2222 | 3 | 0.011 | 1.00E-12 | 1.00E-05 | 1 | 4450 | 6.60E-182 |
| BAYAREALIKE | -1843 | 2 | 0.006 | 0.0043 | 0 | 1 | 3689 | 9.90E-17 |
| BAYAREALIKE+J | -1805 | 3 | 0.006 | 0.0022 | 0.0059 | 1 | 3616 | 1 |
| COF RECONCILED FRESH+ | | | | | | | | |
| Model | LnL | n | d | e | j | w | AICc | AICc_wt |

| | | | | | | | | |
|---|---|---|---|---|---|---|---|---|
| DEC | -2284 | 2 | 0.012 | 1.00E-12 | 0 | 1 | 4571 | 3.90E-175 |
| DEC+J | -2284 | 3 | 0.012 | 1.00E-12 | 1.00E-05 | 1 | 4574 | 1.40E-175 |
| DIVALIKE | -2473 | 2 | 0.013 | 1.00E-12 | 0 | 1 | 4951 | 1.70E-257 |
| DIVALIKE+J | -2473 | 3 | 0.013 | 1.00E-12 | 1.00E-05 | 1 | 4953 | 5.90E-258 |
| BAYAREALIKE | -1897 | 2 | 0.006 | 0.0049 | 0 | 1 | 3798 | 4.00E-07 |
| BAYAREALIKE+J | -1881 | 3 | 0.006 | 0.0041 | 0.0027 | 1 | 3768 | 1 |

FB RECONCILED FRESH+

| Model | LnL | n | d | e | j | w | AICc | AICc_wt |
|---|---|---|---|---|---|---|---|---|
| DEC | -2213 | 2 | 0.012 | 1.00E-12 | 0 | 1 | 4429 | 9.10E-140 |
| DEC+J | -2213 | 3 | 0.012 | 1.00E-12 | 1.00E-05 | 1 | 4431 | 3.20E-140 |
| DIVALIKE | -2388 | 2 | 0.012 | 1.00E-12 | 0 | 1 | 4781 | 3.90E-216 |
| DIVALIKE+J | -2389 | 3 | 0.012 | 1.00E-12 | 1.00E-05 | 1 | 4783 | 1.40E-216 |
| BAYAREALIKE | -1909 | 2 | 0.006 | 0.0051 | 0 | 1 | 3823 | 4.50E-08 |
| BAYAREALIKE+J | -1891 | 3 | 0.006 | 0.0034 | 0.004 | 1 | 3789 | 1 |

COF RECONCILED FRESH ONLY

| Model | LnL | n | d | e | j | w | AICc | AICc_wt |
|---|---|---|---|---|---|---|---|---|
| DEC | -1834 | 2 | 0.008 | 2.80E-05 | 0 | 1 | 3672 | 8.10E-101 |
| DEC+J | -1834 | 3 | 0.008 | 2.70E-05 | 1.00E-05 | 1 | 3674 | 2.90E-101 |
| DIVALIKE | -1969 | 2 | 0.009 | 1.00E-12 | 0 | 1 | 3941 | 2.70E-159 |
| DIVALIKE+J | -1969 | 3 | 0.009 | 1.00E-12 | 1.00E-05 | 1 | 3943 | 9.70E-160 |
| BAYAREALIKE | -1634 | 2 | 0.005 | 0.0043 | 0 | 1 | 3273 | 4.70E-14 |
| BAYAREALIKE+J | -1603 | 3 | 0.004 | 0.0026 | 0.0052 | 1 | 3211 | 1 |

FB RECONCILED FRESH ONLY

| Model | LnL | n | d | e | j | w | AICc | AICc_wt |
|---|---|---|---|---|---|---|---|---|
| DEC | -1795 | 2 | 0.008 | 3.50E-05 | 0 | 1 | 3594 | 1.60E-98 |
| DEC+J | -1795 | 3 | 0.008 | 3.50E-05 | 1.00E-05 | 1 | 3596 | 5.60E-99 |

| | | | | | | | | |
|---|---|---|---|---|---|---|---|---|
| DIVALIKE | -1915 | 2 | 0.009 | 1.00E-12 | 0 | 1 | 3833 | 1.70E-150 |
| DIVALIKE+J | -1915 | 3 | 0.009 | 1.00E-12 | 1.00E-05 | 1 | 3835 | 6.20E-151 |
| BAYAREALIKE | -1615 | 2 | 0.005 | 0.0043 | 0 | 1 | 3235 | 1.50E-20 |
| BAYAREALIKE+J | -1569 | 3 | 0.005 | 0.0021 | 0.0062 | 1 | 3144 | 1 |

**Table S21.** Model-fitting results from BioGeoBEARS based on RAxML Master Tree using six dataset schemes: Catalogue of Fishes (CoF) and Fishbase (FB) raw data, CoF and FB reconciled data accounting for freshwater species with broad habitat range (‘Fresh+’ datasets), and CoF and FB reconciled data accounting for freshwater species with narrow habitat range (‘Fresh Only’ datasets). Biogeographic models include dispersal-extinction-cladogenesis (DEC), a likelihood version of dispersal-vicariance (DIVALIKE), and a likelihood of the Bayarea (BAYAREALIKE) model. All three models were inferred with and without the jump-dispersal (J) parameter. LnL: Log-likelihood; n: number of parameters estimated for each model; d: dispersal rate; e: extinction rate; w: dispersal matrix power exponential (set 1 as default); AICc: Akaike Information Criterion; AICc_wt: Akaike weight.

| TREE - RAxML MASTER TREE | | | | | | | | |
|---|---|---|---|---|---|---|---|---|
| COF RAW DATA | | | | | | | | |
| Model | LnL | n | d | e | j | w | AICc | AICc_wt |
| DEC | -2076 | 2 | 0.0104 | 1.00E-12 | 0.00E+00 | 1 | 4156 | 3.59E-127 |
| DEC+J | -2076 | 3 | 0.0104 | 1.00E-12 | 1.00E-05 | 1 | 4158 | 1.28E-127 |
| DIVALIKE | -2254 | 2 | 0.0111 | 1.00E-12 | 0.00E+00 | 1 | 4513 | 1.27E-204 |
| DIVALIKE+J | -2254 | 3 | 0.0111 | 1.00E-12 | 1.00E-05 | 1 | 4515 | 4.65E-205 |
| BAYAREALIKE | -1806 | 2 | 0.0057 | 4.28E-03 | 0.00E+00 | 1 | 3616 | 5.99E-10 |
| BAYAREALIKE+J | -1784 | 3 | 0.0057 | 2.73E-03 | 4.44E-03 | 1 | 3574 | 1.00E+00 |
| FB RAW DATA | | | | | | | | |
| Model | LnL | n | d | e | j | w | AICc | AICc_wt |
| DEC | -2054 | 2 | 0.0105 | 1.00E-12 | 0 | 1 | 4113 | 8.13E-115 |
| DEC+J | -2054 | 3 | 0.0105 | 1.00E-12 | 1.00E-05 | 1 | 4115 | 2.91E-115 |
| DIVALIKE | -2225 | 2 | 0.0113 | 1.00E-12 | 0 | 1 | 4454 | 5.40E-189 |
| DIVALIKE+J | -2225 | 3 | 0.0113 | 1.00E-12 | 1.00E-05 | 1 | 4456 | 1.94E-189 |

| | | | | | | | | |
|---|---|---|---|---|---|---|---|---|
| BAYAREALIKE | -1834 | 2 | 0.0061 | 4.21E-03 | 0 | 1 | 3673 | 2.43E-19 |
| BAYAREALIKE+J | -1791 | 3 | 0.006 | 2.14E-03 | 0.00573 | 1 | 3587 | 1.00E+00 |

COF RECONCILED FRESH+

| Model | LnL | n | d | e | j | w | AICc | AICc_wt |
|---|---|---|---|---|---|---|---|---|
| DEC | -2278 | 2 | 0.0118 | 1.00E-12 | 0 | 1 | 4560 | 5.21E-179 |
| DEC+J | -2278 | 3 | 0.0118 | 1.00E-12 | 1.00E-05 | 1 | 4563 | 1.85E-179 |
| DIVALIKE | -2476 | 2 | 0.0127 | 1.00E-12 | 0 | 1 | 4957 | 4.10E-265 |
| DIVALIKE+J | -2477 | 3 | 0.0127 | 1.00E-12 | 1.00E-05 | 1 | 4959 | 1.48E-265 |
| BAYAREALIKE | -1886 | 2 | 0.0057 | 4.69E-03 | 0 | 1 | 3776 | 1.24E-08 |
| BAYAREALIKE+J | -1867 | 3 | 0.0055 | 3.64E-03 | 0.00343 | 1 | 3739 | 1.00E+00 |

FB RECONCILED FRESH+

| Model | LnL | n | d | e | j | w | AICc | AICc_wt |
|---|---|---|---|---|---|---|---|---|
| DEC | -2208 | 2 | 0.0117 | 1.00E-12 | 0 | 1 | 4420 | 1.51E-144 |
| DEC+J | -2208 | 3 | 0.0117 | 1.00E-12 | 1.00E-05 | 1 | 4422 | 5.39E-145 |
| DIVALIKE | -2391 | 2 | 0.0125 | 1.00E-12 | 0 | 1 | 4785 | 7.88E-224 |
| DIVALIKE+J | -2391 | 3 | 0.0125 | 1.00E-12 | 1.00E-05 | 1 | 4787 | 2.82E-224 |
| BAYAREALIKE | -1900 | 2 | 0.006 | 4.88E-03 | 0 | 1 | 3805 | 6.94E-11 |
| BAYAREALIKE+J | -1876 | 3 | 0.0063 | 2.79E-03 | 0.00477 | 1 | 3758 | 1.00E+00 |

COF RECONCILED FRESH ONLY

| Model | LnL | n | d | e | j | w | AICc | AICc_wt |
|---|---|---|---|---|---|---|---|---|
| DEC | -1828 | 2 | 0.0083 | 3.02E-05 | 0 | 1 | 3660 | 2.77E-108 |
| DEC+J | -1828 | 3 | 0.0083 | 3.06E-05 | 1.00E-05 | 1 | 3662 | 9.91E-109 |
| DIVALIKE | -1975 | 2 | 0.0088 | 5.88E-06 | 0 | 1 | 3954 | 5.30E-172 |
| DIVALIKE+J | -1975 | 3 | 0.0088 | 1.59E-08 | 1.82E-04 | 1 | 3956 | 2.11E-172 |
| BAYAREALIKE | -1618 | 2 | 0.0046 | 4.05E-03 | 0 | 1 | 3240 | 5.88E-17 |
| BAYAREALIKE+J | -1580 | 3 | 0.0043 | 2.43E-03 | 0.00527 | 1 | 3165 | 1.00E+00 |

FB RECONCILED FRESH ONLY

| Model | LnL | n | d | e | j | w | AICc | AICc_wt |
|---|---|---|---|---|---|---|---|---|
| DEC | -1788 | 2 | 0.0081 | 5.51E-05 | 0 | 1 | 3580 | 3.66E-103 |

| | | | | | | | | |
|---|---|---|---|---|---|---|---|---|
| DEC+J | -1788 | 3 | 0.0081 | 5.49E-05 | 1.00E-05 | 1 | 3582 | 1.32E-103 |
| DIVALIKE | -1918 | 2 | 0.0086 | 7.27E-06 | 0 | 1 | 3841 | 1.11E-159 |
| DIVALIKE+J | -1918 | 3 | 0.0086 | 7.32E-06 | 1.00E-05 | 1 | 3843 | 4.03E-160 |
| BAYAREALIKE | -1605 | 2 | 0.0047 | 4.02E-03 | 0 | 1 | 3214 | 1.47E-23 |
| BAYAREALIKE+J | -1551 | 3 | 0.0044 | 1.96E-03 | 0.00624 | 1 | 3109 | 1.00E+00 |

**Table S22.** Model-fitting results from BioGeoBEARS based on ASTRAL subset of 8 trees using the Reconciled Fresh+ Catalog of Fishes (CoF) and Fishbase (FB) datasets. Model-fitting for Reconciled Fresh+ dataset based on the two Master Trees can be found in Tables S20–S21. Biogeographic models include dispersal-extinction-cladogenesis (DEC), a likelihood version of dispersal-vicariance (DIVALIKE), and a likelihood of the Bayarea (BAYAREALIKE) model. All three models were inferred with and without the jump-dispersal (J) parameter. LnL: Log-likelihood; n: number of parameters estimated for each model; d: dispersal rate; e: extinction rate; w: dispersal matrix power exponential (set 1 as default); AICc: Akaike Information Criterion; AICc_wt: Akaike weight.

| Dataset - CoF Reconciled Fresh+ | | | | | | | | |
|---|---|---|---|---|---|---|---|---|
| TREE 3 | | | | | | | | |
| Model | LnL | n | d | e | j | w | AICc | AICc_wt |
| DEC | -2283 | 2 | 0.012 | 1.00E-12 | 0 | 1 | 4569 | 5.20E-175 |
| DEC+J | -2283 | 3 | 0.012 | 1.00E-12 | 1.00E-05 | 1 | 4571 | 1.80E-175 |
| DIVALIKE | -2476 | 2 | 0.013 | 1.00E-12 | 0 | 1 | 4956 | 6.00E-259 |
| DIVALIKE+J | -2476 | 3 | 0.013 | 1.00E-12 | 1.00E-05 | 1 | 4958 | 2.20E-259 |
| BAYAREALIKE | -1898 | 2 | 0.0058 | 0.0048 | 0 | 1 | 3800 | 6.40E-08 |
| BAYAREALIKE+J | -1880 | 3 | 0.0056 | 0.004 | 0.0028 | 1 | 3767 | 1 |
| TREE 4 | | | | | | | | |
| Model | LnL | n | d | e | j | w | AICc | AICc_wt |
| DEC | -2288 | 2 | 0.012 | 1.00E-12 | 0 | 1 | 4581 | 1.20E-178 |
| DEC+J | -2289 | 3 | 0.012 | 1.00E-12 | 1.00E-05 | 1 | 4583 | 4.40E-179 |
| DIVALIKE | -2480 | 2 | 0.013 | 1.00E-12 | 0 | 1 | 4965 | 5.60E-262 |
| DIVALIKE+J | -2480 | 3 | 0.013 | 1.00E-12 | 1.00E-05 | 1 | 4967 | 2.00E-262 |
| BAYAREALIKE | -1895 | 2 | 0.0057 | 0.0048 | 0 | 1 | 3795 | 7.00E-08 |
| BAYAREALIKE+J | -1878 | 3 | 0.0055 | 0.004 | 0.0027 | 1 | 3762 | 1 |

| TREE 5 | | | | | | | | |
|---|---|---|---|---|---|---|---|---|
| Model | LnL | n | d | e | j | w | AICc | AICc_wt |
| DEC | -2285 | 2 | 0.012 | 8.50E-10 | 0 | 1 | 4575 | 1.30E-177 |
| DEC+J | -2285 | 3 | 0.012 | 1.00E-12 | 1.00E-05 | 1 | 4577 | 4.70E-178 |
| DIVALIKE | -2483 | 2 | 0.013 | 1.00E-12 | 0 | 1 | 4969 | 3.30E-263 |
| DIVALIKE+J | -2483 | 3 | 0.013 | 1.00E-12 | 1.00E-05 | 1 | 4971 | 1.20E-263 |
| BAYAREALIKE | -1893 | 2 | 0.0057 | 0.0048 | 0 | 1 | 3791 | 2.30E-07 |
| BAYAREALIKE+J | -1877 | 3 | 0.0055 | 0.004 | 0.0028 | 1 | 3760 | 1 |
| TREE 6 | | | | | | | | |
| Model | LnL | n | d | e | j | w | AICc | AICc_wt |
| DEC | -2280 | 2 | 0.012 | 5.10E-09 | 0 | 1 | 4564 | 7.80E-178 |
| DEC+J | -2280 | 3 | 0.012 | 1.00E-12 | 1.00E-05 | 1 | 4566 | 2.80E-178 |
| DIVALIKE | -2475 | 2 | 0.013 | 1.00E-12 | 0 | 1 | 4954 | 1.50E-262 |
| DIVALIKE+J | -2475 | 3 | 0.013 | 1.00E-12 | 1.00E-05 | 1 | 4956 | 5.30E-263 |
| BAYAREALIKE | -1888 | 2 | 0.0058 | 0.0047 | 0 | 1 | 3780 | 1.60E-07 |
| BAYAREALIKE+J | -1871 | 3 | 0.0056 | 0.0039 | 0.0029 | 1 | 3748 | 1 |
| TREE 7 | | | | | | | | |
| Model | LnL | n | d | e | j | w | AICc | AICc_wt |
| DEC | -2283 | 2 | 0.012 | 1.00E-12 | 0 | 1 | 4570 | 1.10E-175 |
| DEC+J | -2283 | 3 | 0.012 | 1.00E-12 | 1.00E-05 | 1 | 4572 | 3.80E-176 |
| DIVALIKE | -2478 | 2 | 0.013 | 1.00E-12 | 0 | 1 | 4961 | 1.40E-260 |
| DIVALIKE+J | -2478 | 3 | 0.013 | 1.00E-12 | 1.00E-05 | 1 | 4963 | 4.90E-261 |
| BAYAREALIKE | -1898 | 2 | 0.0058 | 0.0047 | 0 | 1 | 3799 | 2.20E-08 |
| BAYAREALIKE+J | -1879 | 3 | 0.0057 | 0.0036 | 0.0036 | 1 | 3764 | 1 |
| TREE 8 | | | | | | | | |
| Model | LnL | n | d | e | j | w | AICc | AICc_wt |
| DEC | -2280 | 2 | 0.012 | 4.70E-09 | 0 | 1 | 4564 | 3.50E-177 |
| DEC+J | -2280 | 3 | 0.012 | 1.00E-12 | 1.00E-05 | 1 | 4566 | 1.30E-177 |
| DIVALIKE | -2474 | 2 | 0.013 | 1.00E-12 | 0 | 1 | 4952 | 2.30E-261 |
| DIVALIKE+J | -2474 | 3 | 0.013 | 1.00E-12 | 1.00E-05 | 1 | 4954 | 8.10E-262 |
| BAYAREALIKE | -1890 | 2 | 0.0057 | 0.0048 | 0 | 1 | 3785 | 5.70E-08 |
| BAYAREALIKE+J | -1873 | 3 | 0.0056 | 0.0037 | 0.0034 | 1 | 3752 | 1 |
| TREE 9 | | | | | | | | |

| Model | LnL | n | d | e | j | w | AICc | AICc_wt |
|---|---|---|---|---|---|---|---|---|
| DEC | -2270 | 2 | 0.012 | 1.00E-12 | 0 | 1 | 4544 | 2.30E-178 |
| DEC+J | -2270 | 3 | 0.012 | 1.00E-12 | 1.00E-05 | 1 | 4547 | 8.10E-179 |
| DIVALIKE | -2463 | 2 | 0.013 | 1.00E-12 | 0 | 1 | 4930 | 3.80E-262 |
| DIVALIKE+J | -2463 | 3 | 0.013 | 1.00E-12 | 1.00E-05 | 1 | 4932 | 1.30E-262 |
| BAYAREALIKE | -1882 | 2 | 0.0058 | 0.0047 | 0 | 1 | 3768 | 9.10E-10 |
| BAYAREALIKE+J | -1860 | 3 | 0.0056 | 0.0037 | 0.0031 | 1 | 3726 | 1 |
| TREE 10 | | | | | | | | |
| Model | LnL | n | d | e | j | w | AICc | AICc_wt |
| DEC | -2277 | 2 | 0.012 | 1.00E-12 | 0 | 1 | 4559 | 5.80E-177 |
| DEC+J | -2277 | 3 | 0.012 | 1.00E-12 | 1.00E-05 | 1 | 4561 | 2.10E-177 |
| DIVALIKE | -2470 | 2 | 0.013 | 1.00E-12 | 0 | 1 | 4945 | 7.60E-261 |
| DIVALIKE+J | -2470 | 3 | 0.013 | 1.00E-12 | 1.00E-05 | 1 | 4947 | 2.70E-261 |
| BAYAREALIKE | -1891 | 2 | 0.0058 | 0.0048 | 0 | 1 | 3787 | 2.40E-09 |
| BAYAREALIKE+J | -1870 | 3 | 0.0057 | 0.0036 | 0.0032 | 1 | 3747 | 1 |

**Table S23.** Model-fitting results from BioGeoBEARS based on RAxML subset of 8 trees using the Reconciled Fresh+ Catalog of Fishes (CoF) and Fishbase (FB) datasets. Model-fitting for Reconciled Fresh+ dataset based on the two Master Trees can be found in Tables S20–S21. Biogeographic models include dispersal-extinction-cladogenesis (DEC), a likelihood version of dispersal-vicariance (DIVALIKE), and a likelihood of the Bayarea (BAYAREALIKE) model. All three models were inferred with and without the jump-dispersal (J) parameter. LnL: Log-likelihood; n: number of parameters estimated for each model; d: dispersal rate; e: extinction rate; w: dispersal matrix power exponential (set 1 as default); AICc: Akaike Information Criterion; AICc_wt: Akaike weight.

| Dataset - FB Reconciled Fresh+ | | | | | | | | |
|---|---|---|---|---|---|---|---|---|
| TREE 3 | | | | | | | | |
| Model | LnL | n | d | e | j | w | AICc | AICc_wt |
| DEC | -2210 | 2 | 0.012 | 1.00E-12 | 0 | 1 | 4425 | 2.90E-139 |
| DEC+J | -2210 | 3 | 0.012 | 1.00E-12 | 1.00E-05 | 1 | 4427 | 1.00E-139 |
| DIVALIKE | -2389 | 2 | 0.012 | 1.00E-12 | 0 | 1 | 4783 | 5.70E-217 |
| DIVALIKE+J | -2389 | 3 | 0.012 | 1.00E-12 | 1.00E-05 | 1 | 4785 | 2.00E-217 |

| | | | | | | | | |
|---|---|---|---|---|---|---|---|---|
| BAYAREALIKE | -1910 | 2 | 0.006 | 0.005 | 0 | 1 | 3824 | 8.70E-09 |
| BAYAREALIKE+J | -1890 | 3 | 0.0064 | 0.0028 | 0.0048 | 1 | 3787 | 1 |
| TREE 4 | | | | | | | | |
| Model | LnL | n | d | e | j | w | AICc | AICc_wt |
| DEC | -2215 | 2 | 0.012 | 1.00E-12 | 0 | 1 | 4433 | 1.10E-141 |
| DEC+J | -2215 | 3 | 0.012 | 1.00E-12 | 1.00E-05 | 1 | 4435 | 4.10E-142 |
| DIVALIKE | -2393 | 2 | 0.012 | 1.00E-12 | 0 | 1 | 4791 | 2.80E-219 |
| DIVALIKE+J | -2393 | 3 | 0.012 | 1.00E-12 | 1.00E-05 | 1 | 4793 | 1.00E-219 |
| BAYAREALIKE | -1909 | 2 | 0.006 | 0.005 | 0 | 1 | 3822 | 6.00E-09 |
| BAYAREALIKE+J | -1889 | 3 | 0.0062 | 0.0032 | 0.0041 | 1 | 3784 | 1 |
| TREE 5 | | | | | | | | |
| Model | LnL | n | d | e | j | w | AICc | AICc_wt |
| DEC | -2214 | 2 | 0.012 | 1.00E-12 | 0 | 1 | 4433 | 2.50E-142 |
| DEC+J | -2214 | 3 | 0.012 | 1.00E-12 | 1.00E-05 | 1 | 4435 | 9.00E-143 |
| DIVALIKE | -2397 | 2 | 0.012 | 1.00E-12 | 0 | 1 | 4799 | 7.50E-222 |
| DIVALIKE+J | -2397 | 3 | 0.012 | 1.00E-12 | 1.00E-05 | 1 | 4801 | 2.70E-222 |
| BAYAREALIKE | -1906 | 2 | 0.006 | 0.005 | 0 | 1 | 3817 | 1.20E-08 |
| BAYAREALIKE+J | -1887 | 3 | 0.0062 | 0.0032 | 0.0042 | 1 | 3780 | 1 |
| TREE 6 | | | | | | | | |
| Model | LnL | n | d | e | j | w | AICc | AICc_wt |
| DEC | -2209 | 2 | 0.012 | 1.00E-12 | 0 | 1 | 4423 | 4.80E-143 |
| DEC+J | -2209 | 3 | 0.012 | 1.00E-12 | 1.00E-05 | 1 | 4425 | 1.70E-143 |
| DIVALIKE | -2391 | 2 | 0.013 | 1.00E-12 | 0 | 1 | 4785 | 8.20E-222 |
| DIVALIKE+J | -2391 | 3 | 0.013 | 1.00E-12 | 1.00E-05 | 1 | 4787 | 2.90E-222 |
| BAYAREALIKE | -1901 | 2 | 0.0061 | 0.0049 | 0 | 1 | 3805 | 5.60E-09 |
| BAYAREALIKE+J | -1881 | 3 | 0.0064 | 0.0029 | 0.0046 | 1 | 3767 | 1 |
| TREE 7 | | | | | | | | |
| Model | LnL | n | d | e | j | w | AICc | AICc_wt |

| | | | | | | | | |
|---|---|---|---|---|---|---|---|---|
| DEC | -2213 | 2 | 0.012 | 1.00E-12 | 0 | 1 | 4430 | 8.20E-142 |
| DEC+J | -2213 | 3 | 0.012 | 1.00E-12 | 1.00E-05 | 1 | 4432 | 2.90E-142 |
| DIVALIKE | -2394 | 2 | 0.013 | 1.00E-12 | 0 | 1 | 4791 | 2.80E-220 |
| DIVALIKE+J | -2394 | 3 | 0.013 | 1.00E-12 | 1.00E-05 | 1 | 4793 | 1.00E-220 |
| BAYAREALIKE | -1912 | 2 | 0.0061 | 0.0049 | 0 | 1 | 3827 | 5.00E-11 |
| BAYAREALIKE+J | -1887 | 3 | 0.0064 | 0.0027 | 0.005 | 1 | 3780 | 1 |
| TREE 8 | | | | | | | | |
| Model | LnL | n | d | e | j | w | AICc | AICc_wt |
| DEC | -2210 | 2 | 0.012 | 1.00E-12 | 0 | 1 | 4425 | 1.20E-142 |
| DEC+J | -2210 | 3 | 0.012 | 1.00E-12 | 1.00E-05 | 1 | 4427 | 4.50E-143 |
| DIVALIKE | -2391 | 2 | 0.013 | 1.00E-12 | 0 | 1 | 4786 | 3.60E-221 |
| DIVALIKE+J | -2391 | 3 | 0.013 | 1.00E-12 | 1.00E-05 | 1 | 4788 | 1.30E-221 |
| BAYAREALIKE | -1906 | 2 | 0.006 | 0.005 | 0 | 1 | 3817 | 1.30E-10 |
| BAYAREALIKE+J | -1883 | 3 | 0.0063 | 0.0028 | 0.0048 | 1 | 3771 | 1 |
| TREE 9 | | | | | | | | |
| Model | LnL | n | d | e | j | w | AICc | AICc_wt |
| DEC | -2201 | 2 | 0.012 | 1.00E-12 | 0 | 1 | 4406 | 3.60E-144 |
| DEC+J | -2201 | 3 | 0.012 | 1.00E-12 | 1.00E-05 | 1 | 4408 | 1.30E-144 |
| DIVALIKE | -2380 | 2 | 0.013 | 1.00E-12 | 0 | 1 | 4763 | 1.00E-221 |
| DIVALIKE+J | -2380 | 3 | 0.013 | 1.00E-12 | 1.00E-05 | 1 | 4765 | 3.50E-222 |
| BAYAREALIKE | -1895 | 2 | 0.006 | 0.0049 | 0 | 1 | 3795 | 1.70E-11 |
| BAYAREALIKE+J | -1870 | 3 | 0.0063 | 0.0029 | 0.0043 | 1 | 3745 | 1 |
| TREE 10 | | | | | | | | |
| Model | LnL | n | d | e | j | w | AICc | AICc_wt |
| DEC | -2207 | 2 | 0.012 | 1.00E-12 | 0 | 1 | 4418 | 8.90E-143 |
| DEC+J | -2207 | 3 | 0.012 | 1.00E-12 | 1.00E-05 | 1 | 4420 | 3.20E-143 |
| DIVALIKE | -2385 | 2 | 0.013 | 1.00E-12 | 0 | 1 | 4773 | 8.30E-220 |
| DIVALIKE+J | -2385 | 3 | 0.013 | 1.00E-12 | 1.00E-05 | 1 | 4775 | 2.90E-220 |

| | | | | | | | | |
|---|---|---|---|---|---|---|---|---|
| BAYAREALIKE | -1904 | 2 | 0.0061 | 0.0049 | 0 | 1 | 3813 | 2.80E-11 |
| BAYAREALIKE+J | -1879 | 3 | 0.0064 | 0.0028 | 0.0045 | 1 | 3764 | 1 |

**Table S24.** Maximum number of freshwater colonization events in the Sahul fish fauna during the Cenozoic (56-2 Ma). Biogeographic stochastic mapping analyses were based on a set of 100 biostochastic maps generated for each run. Here, we used a set of 10 trees, including ASTRAL and RAxML Master Trees (MT). Mean number of colonization events by 2 Ma bins can be found in the Supplementary Excel file.

| Tree | Database | Max number of colonizations |
|---|---|---|
| Tree1_ASTRAL_MT | CoF_Fresh+ | 22 |
| Tree1_ASTRAL_MT | FB_Fresh+ | 33 |
| Tree16_RAxML_MT | CoF_Fresh+ | 24.2 |
| Tree16_RAxML_MT | FB_Fresh+ | 34 |
| Tree17_RAxML | CoF_Fresh+ | 27.1 |
| Tree17_RAxML | FB_Fresh+ | 35.1 |
| Tree18_RAxML | CoF_Fresh+ | 27 |
| Tree18_RAxML | FB_Fresh+ | 34 |
| Tree19_RAxML | CoF_Fresh+ | 25 |
| Tree19_RAxML | FB_Fresh+ | 34.2 |
| Tree2_ASTRAL | CoF_Fresh+ | 24 |
| Tree2_ASTRAL | FB_Fresh+ | 36.1 |
| Tree20_RAxML | CoF_Fresh+ | 24.1 |
| Tree20_RAxML | FB_Fresh+ | 33 |
| Tree3_ASTRAL | CoF_Fresh+ | 21 |
| Tree3_ASTRAL | FB_Fresh+ | 34.2 |
| Tree4_ASTRAL | CoF_Fresh+ | 23 |
| Tree4_ASTRAL | FB_Fresh+ | 35.2 |
| Tree5_ASTRAL | CoF_Fresh+ | 27 |
| Tree5_ASTRAL | FB_Fresh+ | 38.2 |

**Table S25.** Topology comparison between subclades using CoF Reconciled Fresh+ and FB Reconciled Fresh+ datasets. Note that the only differing subclades are Ambassidae and Terapontidae.

| Clade | Family | Tips_Topology | N_tips_CoF_Fresh+ | N_tips_FB_Fresh+ | CoF_Fresh+_FB_Fresh+_Topology_Comparison |
|---|---|---|---|---|---|
| Amb | Ambassidae | DIFFERENT tips/topology | 18 (including outgroup) | 16 (including outgroup) | Same outgroup; FB_Fresh+ without A. ambassis & A. kopsii |
| Ang | Anguillidae | SAME tips/topology | 8 (including outgroup) | 8 (including outgroup) | Same outgroup |
| Ari | Ariidae | SAME tips/topology | 28 (including outgroup) | 28 (including outgroup) | Same outgroup |
| Ath | Atherinidae | SAME tips/topology | 21 (including outgroup) | 21 (including outgroup) | Same outgroup |
| But | Butidae | SAME tips/topology | 14 (including outgroup) | 14 (including outgroup) | Same outgroup |
| Clu_1 | Clupeidae | SAME tips/topology | 3 (including outgroup) | 3 (including outgroup) | Same outgroup |
| Clu_2 | Clupeidae | SAME tips/topology | 6 (including outgroup) | 6 (including outgroup) | Same outgroup |
| Ele_1 | Eleotridae | SAME tips/topology | 8 (including outgroup) | 8 (including outgroup) | Same outgroup |
| Ele_2 | Eleotridae | SAME tips/topology | 14 (including outgroup) | 14 (including outgroup) | Same outgroup |
| Ele_3 | Eleotridae | SAME tips/topology | 8 (including outgroup) | 8 (including outgroup) | Same outgroup |
| Ele_4 | Eleotridae | SAME tips/topology | 4 (including outgroup) | 4 (including outgroup) | Same outgroup |
| Gob | Gobiidae | SAME tips/topology | 6 (including outgroup) | 6 (including outgroup) | Same outgroup |
| Kuh | Kuhlidae | SAME tips/topology | 7 (including outgroup) | 7 (including outgroup) | Same outgroup |

| | | | | | |
|---|---|---|---|---|---|
| Mug | Mugilidae | SAME tips/topology | 33 (including outgroup) | 33 (including outgroup) | Same outgroup |
| Oxu_1 | Oxudercidae | SAME tips/topology | 6 (including outgroup) | 6 (including outgroup) | Same outgroup |
| Oxu_2 | Oxudercidae | SAME tips/topology | 3 (including outgroup) | 3 (including outgroup) | Same outgroup |
| Oxu_3 | Oxudercidae | SAME tips/topology | 4 (including outgroup) | 4 (including outgroup) | Same outgroup |
| Plo | Plotosidae | SAME tips/topology | 22 (including outgroup) | 22 (including outgroup) | Same outgroup |
| PseMel | Pseudomugilidae & Melanotaeniidae | SAME tips/topology | 99 (including outgroup) | 99 (including outgroup) | Same outgroup |
| Syn | Synbranchidae | SAME tips/topology | 4 (including outgroup) | 4 (including outgroup) | Same outgroup |
| Ter | Terapontidae | DIFFERENT tips/topology | 37 (including outgroup) | 36 (including outgroup) | Same outgroup; FB_Fresh+ without Rhynchopelates oxyrhynchus |
| Tox | Toxotidae | SAME tips/topology | 4 (including outgroup) | 4 (including outgroup) | Same outgroup |
| Zen | Zenarchopteridae | SAME tips/topology | 5 (including outgroup) | 5 (including outgroup) | Same outgroup |

**Table S26. Subclades identified as freshwater transitions in Sahul and retained for downstream analyses.** We identified 23 subclades that each represent one freshwater transition. Subclades with numbers indicate families with more than one freshwater transition in Sahul (e.g., Clupeidae, with two identified freshwater transitions; see Figs. S8–S9). For each subclade, we report the Fresh+ datasets and the averaged stem ages, crown ages, diversification rates, and functional metrics. Full data are available in Table S14 (clade ages), Tables S27–S30 (diversification rates), and Table S18 (functional metrics).

| | | | Clade Ages | | Diversification rates | | Functional Metrics | |
|---|---|---|---|---|---|---|---|---|
| **Subclade** | **Family** | **Dataset Fresh+** | **Stem Age (Ma)** | **Crown Age (Ma)** | **MoM (ε=0.5)** | **MiSSE** | **Hypervolume Product** | **Geometric Mean** |
| **Amb** | Ambassidae | CoF & FB | 75.5 | 45.4 | 0.03 | 0.04 | 0.101 | 0.955 |
| **Ang** | Anguillidae | CoF & FB | 73.3 | 9.9 | 0.02 | -0.05 | 9.08E-05 | 0.559 |
| **Ari** | Ariidae | CoF & FB | 18.0 | 13.0 | 0.14 | 0.12 | 0.012 | 0.865 |
| **Ath** | Atherinidae | CoF & FB | 32.0 | 20.6 | 0.07 | 0.07 | 0.001 | 0.67 |
| **But** | Butidae | FB | 47.5 | 23.5 | 0.05 | 0.06 | 0.004 | 0.772 |
| **Clu_1** | Clupeidae | CoF & FB | 25.3 | 4.3 | – | -0.16 | – | – |
| **Clu_2** | Clupeidae | CoF & FB | 24.4 | 16.9 | 0.04 | -0.003 | 0.001 | 0.67 |
| **Ele_1** | Eleotridae | FB | 18.4 | 16.0 | 0.15 | 0.10 | 8.03E-07 | 0.359 |
| **Ele_2** | Eleotridae | FB | 30.2 | 9.8 | 0.07 | 0.03 | 4.20E-04 | 0.653 |
| **Ele_3** | Eleotridae | FB | 20.4 | 17.0 | 0.08 | 0.06 | 3.47E-04 | 0.652 |
| **Ele_4** | Eleotridae | FB | 22.7 | 15.9 | 0.03 | 0.04 | 1.09E-06 | 0.371 |
| **Gob** | Gobiidae | FB | 29.6 | 25.9 | 0.08 | 0.08 | 0.001 | 0.707 |
| **Kuh** | Kuhliidae | CoF & FB | 47.3 | 17.8 | 0.02 | -0.02 | 0.013 | 0.85 |
| **Mug** | Mugilidae | CoF & FB | 73.6 | 34.5 | 0.03 | 0.05 | 0.021 | 0.89 |
| **Oxu_1** | Oxudercidae | FB | 45.2 | 23.0 | 0.05 | -0.02 | 0.002 | 0.732 |
| **Oxu_2** | Oxudercidae | FB | 2.0 | 1.6 | 0.46 | 0.45 | – | – |
| **Oxu_3** | Oxudercidae | FB | 11.4 | 10.2 | 0.14 | 0.12 | – | – |
| **Plo** | Plotosidae | CoF & FB | 61.5 | 31.8 | 0.04 | 0.05 | 0.003 | 0.772 |

| | | | | | | | | |
|---|---|---|---|---|---|---|---|---|
| **PseMel** | Pseudomugilidae & Melanotaeniidae | CoF & FB | 48.1 | 42.5 | 0.08 | 0.04 | 0.001 | 0.399 |
| **Syn** | Synbranchidae | CoF & FB | 87.4 | 49.4 | 0.01 | 0.01 | 7.80E-07 | 0.358 |
| **Ter** | Terapontidae | CoF & FB | 47.3 | 34.9 | 0.06 | 0.06 | 0.078 | 0.954 |
| **Tox** | Toxotidae | CoF & FB | 45.7 | 19.4 | 0.02 | -0.001 | – | – |
| **Zen** | Zenarchopteridae | CoF & FB | 30.0 | 21.3 | 0.04 | 0.04 | – | – |

**Table S27.** Net diversification rates based on Method-of-Moments (MoM) under a low extinction scheme (ε=0) averaged by the 30 trees and Reconciled Fresh+ Catalog of Fishes (CoF) and Fishbase (FB) datasets.

| Subset | CoF_Reconc_Fresh+ | | | FB_Reconc_Fresh+ | | |
|---|---|---|---|---|---|---|
| MoM_ε_value | ε=0 | | | | | |
| Subclades | Median | Min | Max | Median | Min | Max |
| Amb | 0.031 | 0.030 | 0.034 | 0.031 | 0.030 | 0.034 |
| Ang | 0.018 | 0.013 | 0.019 | 0.018 | 0.013 | 0.019 |
| Ari | 0.154 | 0.137 | 0.177 | 0.154 | 0.137 | 0.177 |
| Ath | 0.079 | 0.077 | 0.081 | 0.079 | 0.077 | 0.081 |
| But | 0.053 | 0.053 | 0.053 | 0.053 | 0.053 | 0.053 |
| Clu_1 | 0.000 | 0.000 | 0.000 | 0.000 | 0.000 | 0.000 |
| Clu_2 | 0.045 | 0.045 | 0.045 | 0.045 | 0.045 | 0.045 |
| Ele_1 | 0.164 | 0.164 | 0.164 | 0.164 | 0.164 | 0.164 |
| Ele_2 | 0.076 | 0.076 | 0.076 | 0.076 | 0.076 | 0.076 |
| Ele_3 | 0.092 | 0.092 | 0.092 | 0.092 | 0.092 | 0.092 |
| Ele_4 | 0.031 | 0.031 | 0.031 | 0.031 | 0.031 | 0.031 |
| Gob | 0.087 | 0.087 | 0.087 | 0.087 | 0.087 | 0.087 |
| Kuh | 0.023 | 0.022 | 0.024 | 0.023 | 0.022 | 0.024 |
| Mug | 0.037 | 0.035 | 0.040 | 0.037 | 0.035 | 0.040 |
| Oxu_1 | 0.055 | 0.055 | 0.055 | 0.055 | 0.055 | 0.055 |
| Oxu_2 | 0.537 | 0.537 | 0.537 | 0.537 | 0.537 | 0.537 |
| Oxu_3 | 0.157 | 0.157 | 0.157 | 0.157 | 0.157 | 0.157 |
| Plo | 0.049 | 0.041 | 0.051 | 0.049 | 0.041 | 0.051 |
| PseMel | 0.088 | 0.084 | 0.092 | 0.088 | 0.084 | 0.092 |
| Syn | 0.008 | 0.008 | 0.008 | 0.008 | 0.008 | 0.008 |
| Ter | 0.071 | 0.068 | 0.073 | 0.071 | 0.068 | 0.073 |
| Tox | 0.019 | 0.011 | 0.020 | 0.019 | 0.011 | 0.020 |
| Zen | 0.050 | 0.050 | 0.050 | 0.050 | 0.050 | 0.050 |

**Table S28.** Net diversification rates based on Method-of-Moments (MoM) under a moderate extinction scheme (ε=0.5) averaged by the 30 trees and Reconciled Fresh+ Catalog of Fishes (CoF) and Fishbase (FB) datasets.

| Subset | CoF_Reconc_Fresh+ | | | FB_Reconc_Fresh+ | | |
|---|---|---|---|---|---|---|
| MoM_ε_value | ε=0.5 | | | | | |
| Subclades | Median | Min | Max | Median | Min | Max |
| Amb | 0.028 | 0.027 | 0.030 | 0.028 | 0.027 | 0.030 |
| Ang | 0.016 | 0.011 | 0.016 | 0.016 | 0.011 | 0.016 |
| Ari | 0.140 | 0.125 | 0.161 | 0.140 | 0.125 | 0.161 |
| Ath | 0.071 | 0.069 | 0.073 | 0.071 | 0.069 | 0.073 |
| But | 0.048 | 0.048 | 0.048 | 0.048 | 0.048 | 0.048 |
| Clu_1 | 0.000 | 0.000 | 0.000 | 0.000 | 0.000 | 0.000 |
| Clu_2 | 0.039 | 0.039 | 0.039 | 0.039 | 0.039 | 0.039 |
| Ele_1 | 0.150 | 0.150 | 0.150 | 0.150 | 0.150 | 0.150 |
| Ele_2 | 0.068 | 0.068 | 0.068 | 0.068 | 0.068 | 0.068 |
| Ele_3 | 0.081 | 0.081 | 0.081 | 0.081 | 0.081 | 0.081 |
| Ele_4 | 0.026 | 0.026 | 0.026 | 0.026 | 0.026 | 0.026 |
| Gob | 0.078 | 0.078 | 0.078 | 0.078 | 0.078 | 0.078 |
| Kuh | 0.020 | 0.019 | 0.021 | 0.020 | 0.019 | 0.021 |
| Mug | 0.034 | 0.031 | 0.037 | 0.034 | 0.031 | 0.037 |
| Oxu_1 | 0.049 | 0.049 | 0.049 | 0.049 | 0.049 | 0.049 |
| Oxu_2 | 0.461 | 0.461 | 0.461 | 0.461 | 0.461 | 0.461 |
| Oxu_3 | 0.138 | 0.138 | 0.138 | 0.138 | 0.138 | 0.138 |
| Plo | 0.044 | 0.037 | 0.046 | 0.044 | 0.037 | 0.046 |
| PseMel | 0.082 | 0.079 | 0.086 | 0.082 | 0.079 | 0.086 |
| Syn | 0.007 | 0.007 | 0.007 | 0.007 | 0.007 | 0.007 |
| Ter | 0.065 | 0.062 | 0.067 | 0.065 | 0.062 | 0.067 |
| Tox | 0.017 | 0.009 | 0.017 | 0.017 | 0.009 | 0.017 |
| Zen | 0.044 | 0.043 | 0.044 | 0.044 | 0.043 | 0.044 |

**Table S29.** Net diversification rates based on Method-of-Moments (MoM) under a moderate extinction scheme (ε=0.9) averaged by the 30 trees and Reconciled Fresh+ Catalog of Fishes (CoF) and Fishbase (FB) datasets.

| Subset | CoF_Reconc_Fresh+ | | | FB_Reconc_Fresh+ | | |
|---|---|---|---|---|---|---|
| MoM_ε_value | ε=0.9 | | | | | |
| Subclades | Median | Min | Max | Median | Min | Max |
| Amb | 0.014 | 0.014 | 0.015 | 0.014 | 0.014 | 0.015 |
| Ang | 0.006 | 0.004 | 0.006 | 0.006 | 0.004 | 0.006 |
| Ari | 0.076 | 0.068 | 0.087 | 0.076 | 0.068 | 0.087 |
| Ath | 0.037 | 0.036 | 0.038 | 0.037 | 0.036 | 0.038 |
| But | 0.025 | 0.025 | 0.025 | 0.025 | 0.025 | 0.025 |
| Clu_1 | 0.000 | 0.000 | 0.000 | 0.000 | 0.000 | 0.000 |
| Clu_2 | 0.014 | 0.014 | 0.014 | 0.014 | 0.014 | 0.014 |
| Ele_1 | 0.085 | 0.085 | 0.085 | 0.085 | 0.085 | 0.085 |
| Ele_2 | 0.033 | 0.033 | 0.033 | 0.033 | 0.033 | 0.033 |
| Ele_3 | 0.036 | 0.036 | 0.036 | 0.036 | 0.036 | 0.036 |
| Ele_4 | 0.009 | 0.009 | 0.009 | 0.009 | 0.009 | 0.009 |
| Gob | 0.041 | 0.041 | 0.041 | 0.041 | 0.041 | 0.041 |
| Kuh | 0.007 | 0.007 | 0.008 | 0.007 | 0.007 | 0.008 |
| Mug | 0.018 | 0.017 | 0.020 | 0.018 | 0.017 | 0.020 |
| Oxu_1 | 0.025 | 0.025 | 0.025 | 0.025 | 0.025 | 0.025 |
| Oxu_2 | 0.171 | 0.171 | 0.171 | 0.171 | 0.171 | 0.171 |
| Oxu_3 | 0.060 | 0.060 | 0.060 | 0.060 | 0.060 | 0.060 |
| Plo | 0.025 | 0.020 | 0.026 | 0.025 | 0.020 | 0.026 |
| PseMel | 0.055 | 0.052 | 0.057 | 0.055 | 0.052 | 0.057 |
| Syn | 0.002 | 0.002 | 0.002 | 0.002 | 0.002 | 0.002 |
| Ter | 0.039 | 0.037 | 0.040 | 0.039 | 0.037 | 0.040 |
| Tox | 0.006 | 0.003 | 0.006 | 0.006 | 0.003 | 0.006 |
| Zen | 0.018 | 0.018 | 0.018 | 0.018 | 0.018 | 0.018 |

**Table S30.** Missing State Speciation and Extinction (MiSSE) net diversification rates for the whole subclades with one MiSSE state using CoF Reconciled Fresh+ and FB Reconciled Fresh+ datasets. Values indicate the median, minimum (Min) and maximum (Max) considering all 30 trees.

| Subset | CoF_Reconc_Fresh+ | | | FB_Reconc_Fresh+ | | |
|---|---|---|---|---|---|---|
| Subclades | Median | Min | Max | Median | Min | Max |
| Amb | 0.037 | 0.037 | 0.038 | 0.035 | 0.035 | 0.037 |
| Ang | -0.047 | -0.066 | -0.034 | -0.047 | -0.066 | -0.034 |
| Ari | 0.116 | 0.11 | 0.131 | 0.116 | 0.11 | 0.131 |
| Ath | 0.066 | 0.064 | 0.069 | 0.066 | 0.064 | 0.069 |
| But | 0.056 | 0.056 | 0.056 | 0.056 | 0.056 | 0.056 |
| Clu_1 | -0.158 | -0.158 | -0.158 | -0.158 | -0.158 | -0.158 |
| Clu_2 | -0.003 | -0.004 | -0.002 | -0.003 | -0.004 | -0.002 |
| Ele_1 | 0.096 | 0.096 | 0.096 | 0.096 | 0.096 | 0.096 |
| Ele_2 | 0.032 | 0.031 | 0.034 | 0.032 | 0.031 | 0.034 |
| Ele_3 | 0.055 | 0.055 | 0.055 | 0.055 | 0.055 | 0.055 |
| Ele_4 | 0.043 | 0.043 | 0.043 | 0.043 | 0.043 | 0.043 |
| Gob | 0.08 | 0.08 | 0.08 | 0.08 | 0.08 | 0.08 |
| Kuh | -0.017 | -0.026 | 0.006 | -0.017 | -0.026 | 0.006 |
| Mug | 0.053 | 0.048 | 0.056 | 0.053 | 0.048 | 0.056 |
| Oxu_1 | -0.018 | -0.018 | -0.018 | -0.018 | -0.018 | -0.018 |
| Oxu_2 | 0.446 | 0.446 | 0.446 | 0.446 | 0.446 | 0.446 |
| Oxu_3 | 0.116 | 0.116 | 0.116 | 0.116 | 0.116 | 0.116 |
| Plo | 0.053 | 0.04 | 0.056 | 0.053 | 0.04 | 0.056 |
| PseMel | 0.037 | 0.036 | 0.038 | 0.037 | 0.036 | 0.038 |
| Syn | 0.01 | 0.01 | 0.01 | 0.01 | 0.01 | 0.01 |
| Ter | 0.056 | 0.044 | 0.058 | 0.052 | 0.041 | 0.054 |
| Tox | -0.001 | -0.036 | 0.003 | -0.001 | -0.036 | 0.003 |
| Zen | 0.041 | 0.04 | 0.041 | 0.041 | 0.04 | 0.041 |

**Table S31.** Sampling fractions used to run Cladogenic Diversification Rate Shift (ClaDS).

| Clade | Taxa - Focal Family | N of species | Sampling Fraction |
|---|---|---|---|
| Ariidae | Ariidae | 119 | 0.76 |
| Plotosidae | Plotosidae | 23 | 0.55 |
| Atheriniformes | Atherinidae | 56 | 0.7 |
| | Melanotaeniidae | 84 | 0.74 |
| | Pseudomugilidae | 13 | 0.83 |
| | Remaining taxa | 114 | 0.68 |
| Gobiiformes | Butidae | 15 | 0.26 |
| | Eleotridae | 59 | 0.39 |
| | Gobiidae | 343 | 0.24 |
| | Oxudercidae | 205 | 0.27 |
| | Remaining taxa | 19 | 0.24 |
| Centrarchiformes | Kuhliidae | 12 | 0.92 |
| | Terapontidae | 38 | 0.61 |
| | Remaining taxa | 5 | 0.18 |
| Mugiliformes | Ambassidae | 23 | 0.4 |
| | Mugilidae | 51 | 0.68 |
| Anguilliformes | Anguillidae | 17 | 0.94 |
| | Remaining taxa | 3 | 0.01 |
| Clupeiformes | Clupeidae | 82 | 0.48 |
| | Remaning taxa | 98 | 0.39 |
| Anabantaria | Synbranchidae | 3 | 0.1 |
| | Remaining taxa | 8 | 0.03 |
| Carangaria | Toxotidae | 4 | 0.44 |
| | Remaining taxa | 116 | 0.11 |
| Beloniformes | Zenarchopteridae | 24 | 0.38 |
| | Remaining taxa | 50 | 0.24 |

**Table S32.** Cladogenic Diversification Rate Shift (ClaDS) parameters by clades. sigma: variance after speciation events; alpha: trend parameter of diversification increase or decrease; epsilon: extinction-to-speciation rate; lambda: average speciation rate at the root of the tree

| ClaDS_Clade | Focal Family | Tree | sigma | alpha | eps | lambda |
|---|---|---|---|---|---|---|
| Ariidae | Ariidae | ASTRAL_MT | 0.958 | 0.762 | 0.782 | 0.137 |
| Plotosidae | Plotosidae | ASTRAL_MT | 0.209 | 0.869 | 0.180 | 0.148 |
| Atheriniformes | Atherinidae | ASTRAL_MT | 0.220 | 1.055 | 0.049 | 0.051 |
| | Melanotaeniidae | ASTRAL_MT | | | | |
| | Pseudomugilidae | ASTRAL_MT | | | | |
| | Remaining taxa | ASTRAL_MT | | | | |
| Gobiiformes | Butidae | ASTRAL_MT | 0.220 | 0.933 | 0.041 | 0.075 |
| | Eleotridae | ASTRAL_MT | | | | |
| | Gobiidae | ASTRAL_MT | | | | |
| | Oxudercidae | ASTRAL_MT | | | | |
| | Remaining taxa | ASTRAL_MT | | | | |
| Centrarchiformes | Kuhliidae | ASTRAL_MT | 0.231 | 0.941 | 0.212 | 0.089 |
| | Terapontidae | ASTRAL_MT | | | | |
| | Remaining taxa | ASTRAL_MT | | | | |
| Mugiliformes | Ambassidae | ASTRAL_MT | 0.186 | 0.924 | 0.048 | 0.110 |
| | Mugilidae | ASTRAL_MT | | | | |
| Anguilliformes | Anguillidae | ASTRAL_MT | 0.177 | 0.895 | 0.057 | 0.121 |
| | Remaining taxa | ASTRAL_MT | | | | |
| Clupeiformes | Clupeidae | ASTRAL_MT | 0.222 | 0.950 | 0.128 | 0.031 |
| | Remaining taxa | ASTRAL_MT | | | | |
| Anabantaria | Synbranchidae | ASTRAL_MT | 0.164 | 0.923 | 0.073 | 0.109 |
| | Remaining taxa | ASTRAL_MT | | | | |
| Carangaria | Toxotidae | ASTRAL_MT | 0.161 | 0.946 | 0.312 | 0.133 |
| | Remaining taxa | ASTRAL_MT | | | | |
| Beloniformes | Zenarchopteridae | ASTRAL_MT | 0.193 | 0.928 | 0.054 | 0.113 |
| | Remaining taxa | ASTRAL_MT | | | | |
| Ariidae | Ariidae | RAxML_MT | 0.790 | 0.766 | 0.716 | 0.188 |
| Plotosidae | Plotosidae | RAxML_MT | 0.227 | 0.894 | 0.102 | 0.150 |
| Atheriniformes | Atherinidae | RAxML_MT | 0.228 | 1.033 | 0.102 | 0.062 |
| | Melanotaeniidae | RAxML_MT | | | | |
| | Pseudomugilidae | RAxML_MT | | | | |
| | Remaining taxa | RAxML_MT | | | | |
| Gobiiformes | Butidae | RAxML_MT | 0.182 | 0.945 | 0.021 | 0.071 |
| | Eleotridae | RAxML_MT | | | | |
| | Gobiidae | RAxML_MT | | | | |
| | Oxudercidae | RAxML_MT | | | | |
| | Remaining taxa | RAxML_MT | | | | |
| Centrarchiformes | Kuhliidae | RAxML_MT | 0.226 | 0.940 | 0.370 | 0.082 |
| | Terapontidae | RAxML_MT | | | | |

| | | | | | | |
|---|---|---|---|---|---|---|
| | Remaining taxa | RAxML_MT | | | | |
| Mugiliformes | Ambassidae | RAxML_MT | 0.184 | 0.926 | 0.039 | 0.104 |
| | Mugilidae | RAxML_MT | | | | |
| Anguilliformes | Anguillidae | RAxML_MT | 0.206 | 0.897 | 0.066 | 0.122 |
| | Remaining taxa | RAxML_MT | | | | |
| Clupeiformes | Clupeidae | RAxML_MT | 0.207 | 0.955 | 0.057 | 0.033 |
| | Remaining taxa | RAxML_MT | | | | |
| Anabantaria | Synbranchidae | RAxML_MT | 0.203 | 0.955 | 0.092 | 0.119 |
| | Remaining taxa | RAxML_MT | | | | |
| Carangaria | Toxotidae | RAxML_MT | 0.236 | 0.922 | 0.694 | 0.192 |
| | Remaining taxa | RAxML_MT | | | | |
| Beloniformes | Zenarchopteridae | RAxML_MT | 0.186 | 0.934 | 0.097 | 0.124 |
| | Remaining taxa | RAxML_MT | | | | |

**Table S33.** Ordinary (OLS) and phylogenetic least-square (PGLS) regressions under Brownian motion (PGLS-BM) and Ornstein–Uhlenbeck process (PGLS-OU). PGLS parameters are averaged by the 30 trees.

| Correlation | Model | N of subclades | Estimate | p.value | R-squared | Adj_R-squared | AIC |
|---|---|---|---|---|---|---|---|
| log(MoM)~Stem_Age | OLS | n=22 | -0.031 | 0.00003 | 0.59 | 0.57 | 44.4 |
| log(MoM)~Stem_Age | PGLS_BM | n=22; 30 trees | -0.032 | 0.0005 | 0.47 | 0.44 | 54.8 |
| log(MoM)~Stem_Age | PGLS_OU | n=22; 30 trees | -0.031 | 0.00003 | 0.59 | 0.57 | 46.4 |
| log(MoM)~Crown_Age | OLS | n=22 | -0.035 | 0.03 | 0.22 | 0.18 | 58.4 |
| log(MoM)~Crown_Age | PGLS_BM | n=22; 30 trees | -0.034 | 0.05 | 0.18 | 0.14 | 64.3 |
| log(MoM)~Crown_Age | PGLS_OU | n=22; 30 trees | -0.035 | 0.03 | 0.22 | 0.18 | 60.4 |
| log(Hv_product)~log(MoM) | OLS | n=18 | 0.941 | 0.42 | 0.04 | -0.02 | 101.4 |
| log(Hv_product)~log(MoM) | PGLS_BM | n=18; 30 trees | 1.281 | 0.27 | 0.07 | 0.02 | 106.8 |
| log(Hv_product)~log(MoM) | PGLS_OU | n=18; 30 trees | 1.321 | 0.24 | 0.09 | 0.03 | 101.6 |
| Hv_gmean~log(MoM) | OLS | n=18 | 0.031 | 0.63 | 0.01 | -0.05 | -2.3 |
| Hv_gmean~log(MoM) | PGLS_BM | n=18; 30 trees | 0.047 | 0.46 | 0.03 | -0.03 | 3.0 |
| Hv_gmean~log(MoM) | PGLS_OU | n=18; 30 trees | 0.049 | 0.44 | 0.04 | -0.02 | -1.4 |
| log(Hv_product)~Stem_Age | OLS | n=18 | 0.025 | 0.54 | 0.02 | -0.04 | 101.8 |
| log(Hv_product)~Stem_Age | PGLS_BM | n=18; 30 trees | 0.024 | 0.66 | 0.01 | -0.05 | 107.9 |
| log(Hv_product)~Stem_Age | PGLS_OU | n=18; 30 trees | 0.010 | 0.82 | 0.003 | -0.06 | 103.1 |
| Hv_gmean~Stem_Age | OLS | n=18 | 0.001 | 0.57 | 0.02 | -0.04 | -2.4 |
| Hv_gmean~Stem_Age | PGLS_BM | n=18; 30 trees | 0.001 | 0.66 | 0.01 | -0.05 | 3.4 |
| Hv_gmean~Stem_Age | PGLS_OU | n=18; 30 trees | 0.001 | 0.79 | 0.005 | -0.06 | -0.8 |
| log(Hv_product)~Crown_Age | OLS | n=18 | 0.052 | 0.49 | 0.03 | -0.03 | 101.6 |
| log(Hv_product)~Crown_Age | PGLS_BM | n=18; 30 trees | -0.003 | 0.97 | 0.0001 | -0.06 | 108.1 |
| log(Hv_product)~Crown_Age | PGLS_OU | n=18; 30 trees | 0.026 | 0.73 | 0.01 | -0.05 | 103.0 |
| Hv_gmean~Crown_Age | OLS | n=18 | 0.001 | 0.79 | 0.004 | -0.06 | -2.1 |
| Hv_gmean~Crown_Age | PGLS_BM | n=18; 30 trees | -0.002 | 0.65 | 0.01 | -0.05 | 3.4 |

| | | | | | | | |
|---|---|---|---|---|---|---|---|
| Hv_gmean~Crown_Age | PGLS_OU | n=18; 30 trees | -0.0002 | 0.96 | 0.0002 | -0.06 | -0.7 |

**EXTENDED FIGURES**

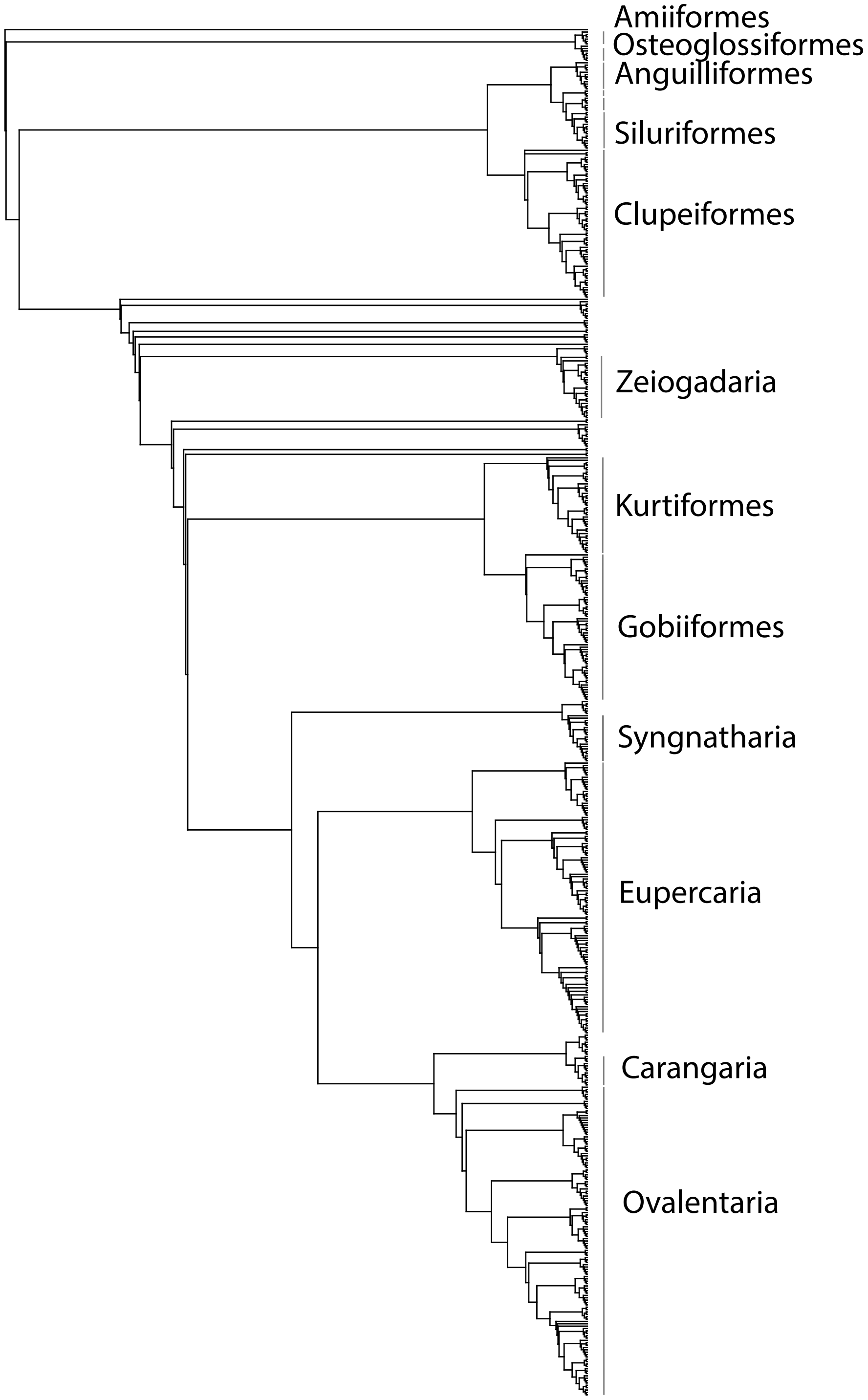

**Figure S1.** Phylogram representing the backbone trees with 578 species. Trees in newick format are available in Supplementary Files.

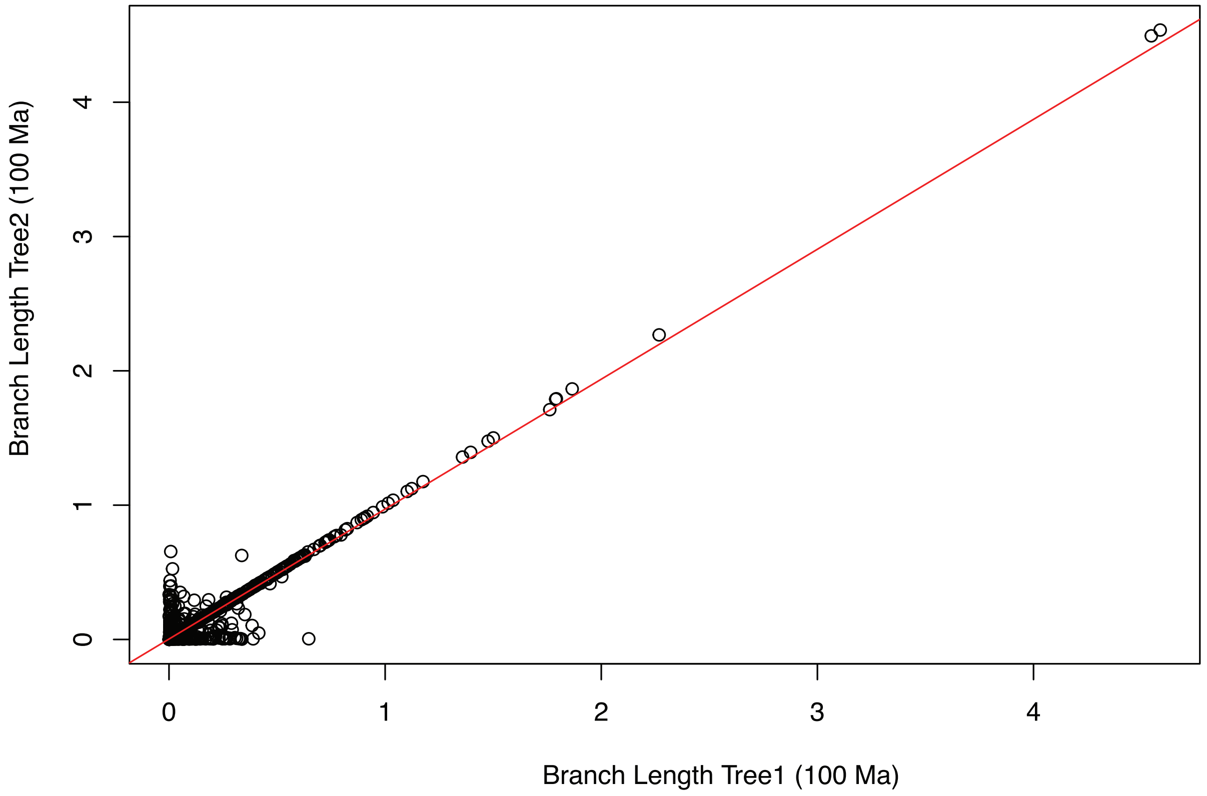

**Figure S2.** Linear regression of branch lengths between two calibrated RAxML trees: one inferred using legacy markers + anchor genes (x-axis) and the other using legacy markers + 825 exon markers (y-axis). The trees show a strong linear relationship ($R^2 = 0.94$), with a slope of 0.97 and an intercept not significantly different from zero, indicating highly consistent branch-length scaling between calibration approaches.

**Figure S3.** Calibrated ASTRAL Master Tree with 1,183 taxa before grafting. Trees in newick format are available in Supplementary Files.

**Figure S4.** Calibrated ASTRAL Master Tree with 2,303 taxa, after grafting. The 30 complete trees in newick format are available in Supplementary Files.

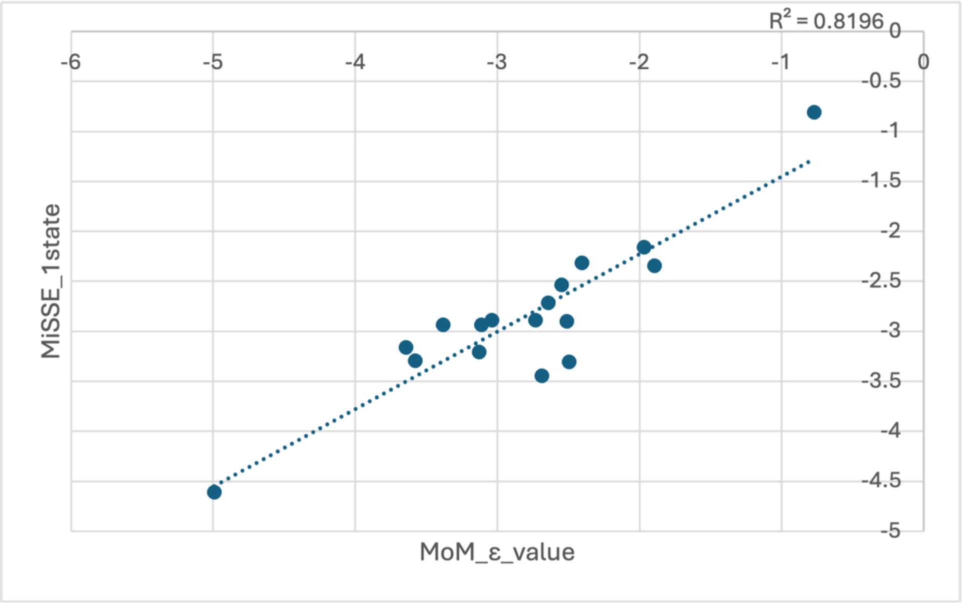

**Figure S5.** Correlation between net diversification estimates from the method-of-moments (MoM; ε = 0.5) and Missing-State Speciation and Extinction (MiSSE) across 17 marine-derived subclades analyzed here. MiSSE was estimated using a single rate-class model (one turnover and one extinction fraction), and net diversification rate was computed as speciation minus extinction. Because MiSSE estimates can be negative, subclades with negative MiSSE values ("Ang," "Clu_1," "Clu_2," "Kuh," "Oxu_1," "Tox") were excluded; the correlation was computed on the remaining 17 subclades.

**Figure S6.** Model-average ancestral habitat range reconstruction based on the 30 trees using Catalogue of Fishes (CoF) raw habitat data for 1,675 species. States 'A', 'B', and 'C' indicate marine only, brackish only and freshwater only habitats, respectively. States colored in grey correspond to multiple habitats (i.e. AB, BC, and ABC). Black dashed lines correspond to 50 million years (my), 30 my, and 15 my. High-resolution figure is available as a supplementary file.

**Figure S7.** Model-average ancestral habitat range reconstruction based on the 30 trees using Fishbase (FB) raw habitat data for 1,665 species. States 'A', 'B', and 'C' indicate marine only, brackish only and freshwater only habitats, respectively. States colored in grey correspond to multiple habitats (i.e. AB, BC, and ABC). Black dashed lines correspond to 50 million years (my), 30 my, and 15 my. High-resolution figure is available as a supplementary file.

**Figure S8.** Model-average ancestral habitat range reconstruction based on the 30 trees using Catalogue of Fishes (CoF) Reconciled Fresh+ habitat data for 1,694 species. States 'A', 'B', and 'C' indicate marine only, brackish only and freshwater only habitats, respectively. States colored in grey correspond to multiple habitats (i.e. AB, BC, and ABC). Black dashed lines correspond to 50 million years (my), 30 my, and 15 my. Green arrow shows an underestimation of timing in bonytongues (Osteoglossidae) marine-to-freshwater transition in Sahul (see supplementary methods). High-resolution figure is available as a supplementary file.

**Figure S9.** Model-average ancestral habitat range reconstruction based on the 30 trees using Fishbase (FB) Reconciled Fresh+ habitat data for 1,667 species. States 'A', 'B', and 'C' indicate marine only, brackish only and freshwater only habitats, respectively. States colored in grey correspond to multiple habitats (i.e. AB, BC, and ABC). Black dashed lines correspond to 50 million years (my), 30 my, and 15 my. Green arrow shows an underestimation of timing in bonytongues (Osteoglossidae) marine-to-freshwater transition in Sahul (see supplementary methods). High-resolution figure is available as a supplementary file.

**Figure S10.** Model-average ancestral habitat range reconstruction based on the 30 trees using Catalogue of Fishes (CoF) Reconciled Fresh Only habitat data for 1,578 species. States 'A', 'B', and 'C' indicate marine only, brackish only and freshwater only habitats, respectively. States colored in grey correspond to multiple habitats (i.e. AB, BC, and ABC). Black dashed lines correspond to 50 million years (my), 30 my, and 15 my. High-resolution figure is available as a supplementary file.

**Figure S11.** Model-average ancestral habitat range reconstruction based on the 30 trees using Fishbase (FB) Reconciled Fresh Only habitat data for 1,561 species. States 'A', 'B', and 'C' indicate marine only, brackish only and freshwater only habitats, respectively. States colored in grey correspond to multiple habitats (i.e. AB, BC, and ABC). Black dashed lines correspond to 50 million years (my), 30 my, and 15 my. High-resolution figure is available as a supplementary file.

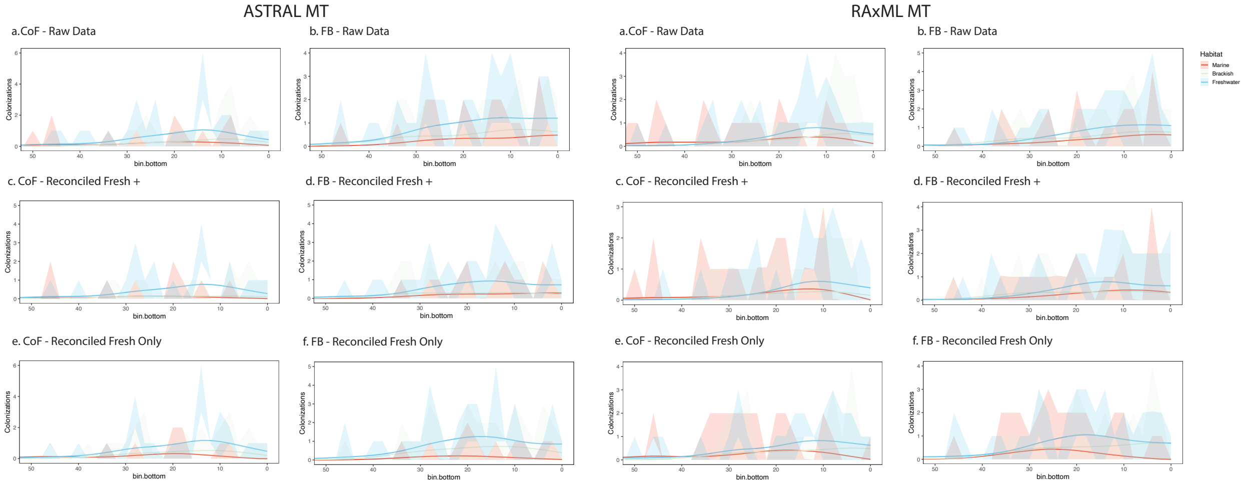

**Figure S12.** Colonization through time plots from BiogeoBEARS ancestral range reconstruction based on 100 biostochastic maps using the ASTRAL and RAxML Master Trees. Six habitat datasets are shown for each tree, including CoF Raw (Catalogue of Fishes raw data) and FB Raw (FishBase raw data), each accounting for freshwater-only, fresh/brackish, and marine/brackish/freshwater species; CoF Reconciled Fresh+ and FB Reconciled Fresh+, based on reconciled Catalogue of Fishes and FishBase datasets and accounting for freshwater-only, fresh/brackish, and marine/brackish/freshwater species; and CoF Reconciled Fresh Only and FB Reconciled Fresh Only, based on reconciled Catalogue of Fishes and FishBase datasets and accounting only for freshwater (including fresh/brackish) species.

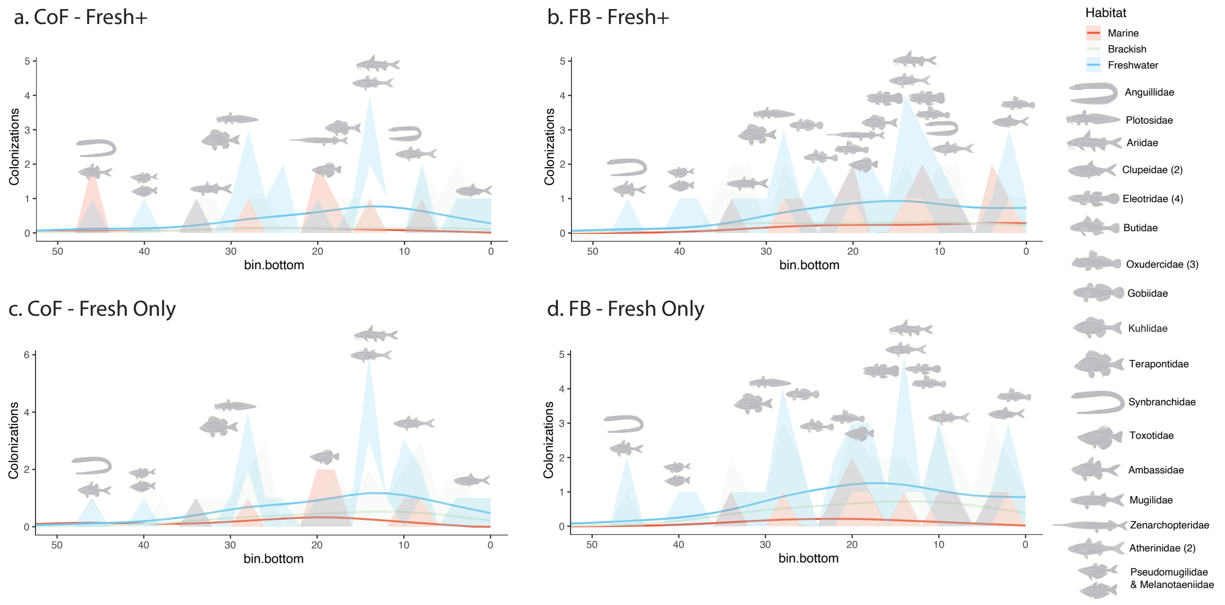

**Figure S13.** Colonization-through-time plots from BioGeoBEARS ancestral range reconstruction (100 stochastic maps) based on the ASTRAL Master Tree. Four habitat datasets are shown: CoF Fresh+ and FB Fresh+ (reconciled Catalogue of Fishes and

FishBase datasets accounting for freshwater-only, fresh/brackish, and marine/brackish/freshwater species), and CoF Reconciled Fresh Only and FB Reconciled Fresh Only (reconciled Catalogue of Fishes and FishBase datasets accounting only for freshwater, including fresh/brackish species). Fish icons indicate families associated with each colonization peak, with families experiencing multiple colonizations depicted according to the number given in parenthesis. Icons shown in light grey denote Gobiaria families in the CoF subsets, where ancestral range reconstruction inferred a freshwater colonization before 50 Ma.

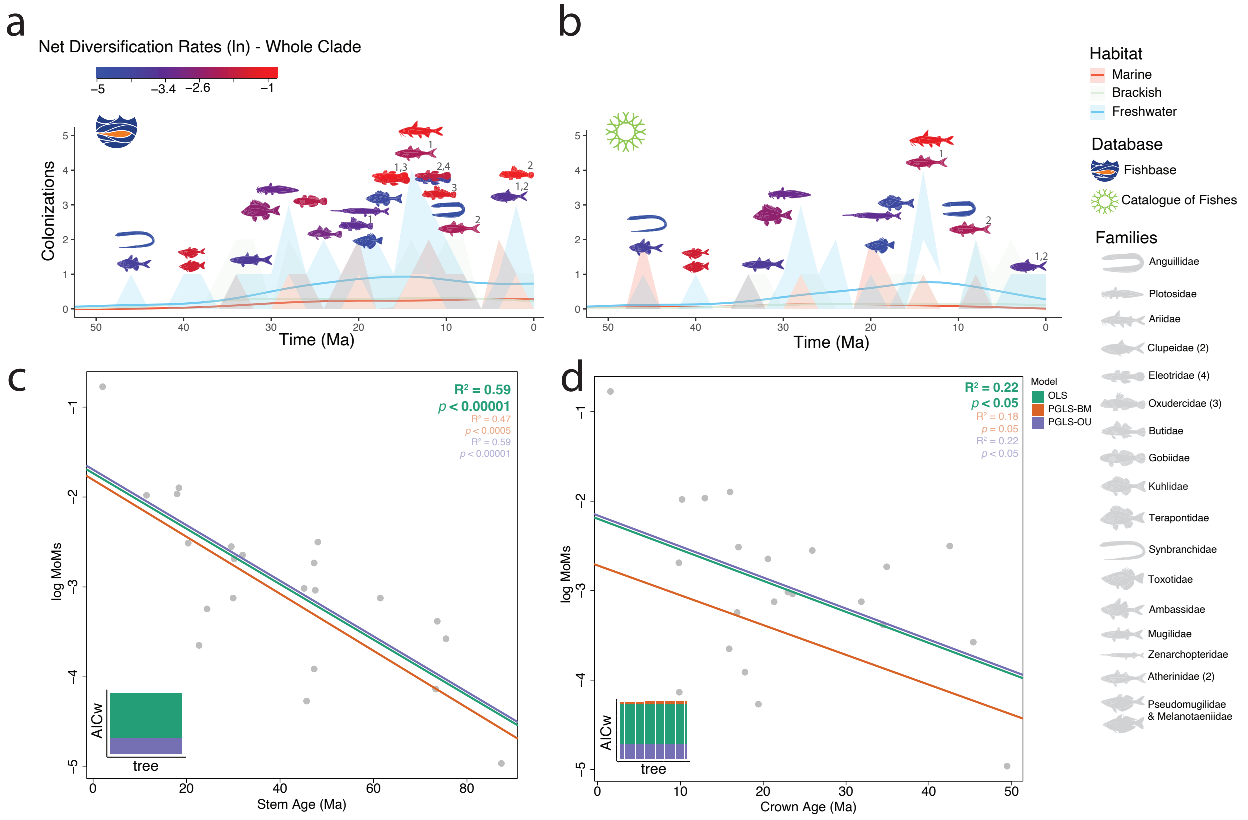

**Figure S14. Net diversification rates and regressions between MoM and clade ages.** Net diversification rates for the whole clade based on MoM for both Fishbase (a) and CoF Fresh+ (b) subsets. Net diversification rates for the whole clade were based on Method-of-Moments MoM ($\varepsilon$= 0.5; see comparison between MoM and MiSSE net diversification rates in Fig. S4). Colonization-through-time plots from BioGeoBEARS ancestral range reconstruction (100 stochastic maps) based on the ASTRAL Master Tree. Fish shapes are colored based on diversification rates' values. Ordinary (OLS) and phylogenetic (PGLS) least-square under Brownian motion (PGLS-BM) and Ornstein–Uhlenbeck process (PGLS-OU) regressions between stem (c) and crown (d) ages (in million years) and averaged net diversification rate (MoM) using ASTRAL MT. Akaike weights (AICw)

are provided by each of the 30 trees. Averaged R-squared and significance (p) are provided in the upper right corner emphasizing (in bold) the best-fit model. Ages used in the regressions are averaged by the 30 trees. PGLS correlations were conducted for all 30 trees (see Table S32).

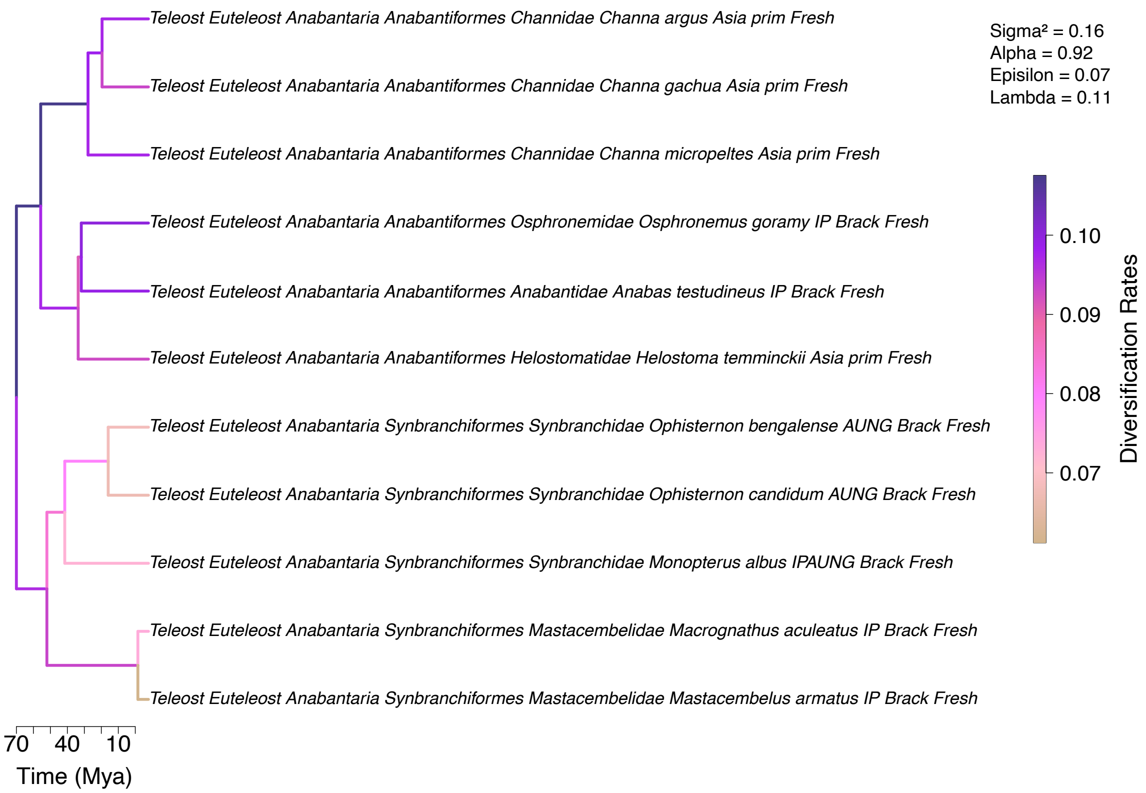

**Figure S15. Anabantaria clade pruned from the complete trees (n = 2,303 species) with speciation rate shifts based on ClaDS.** Focal family: Synbranchidae. ClaDS analyses were performed using ASTRAL and RAxML MTs (see parameters in Table S32). Here, we show ClaDS on ASTRAL MT topology.

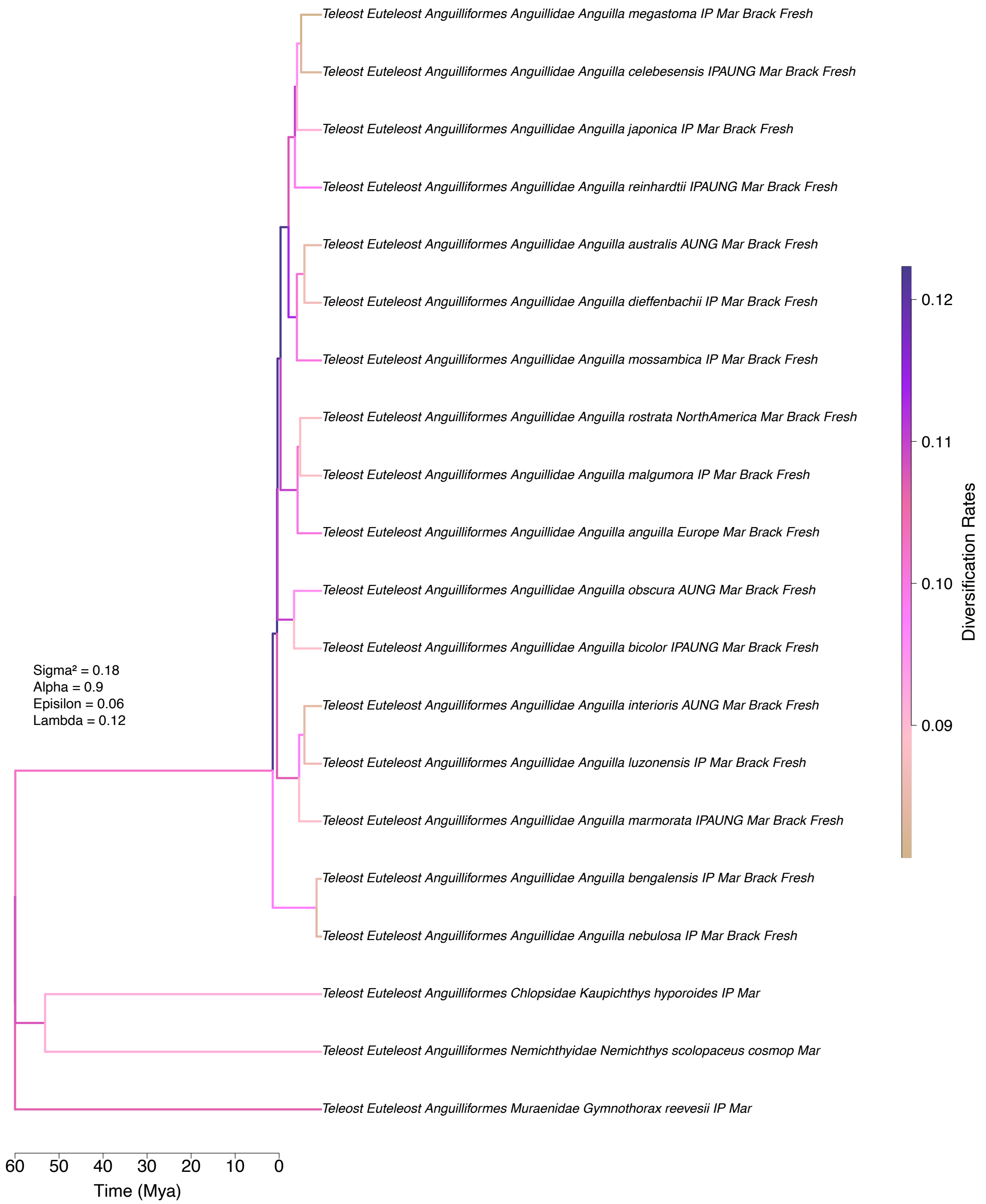

**Figure S16. Anguilliformes clade pruned from the complete trees (n = 2,303 species) with speciation rate shifts based on ClaDS.** Focal family: Anguillidae. ClaDS analyses were performed using ASTRAL and RAxML MTs (see parameters in Table S32). Here, we show ClaDS on ASTRAL MT topology.

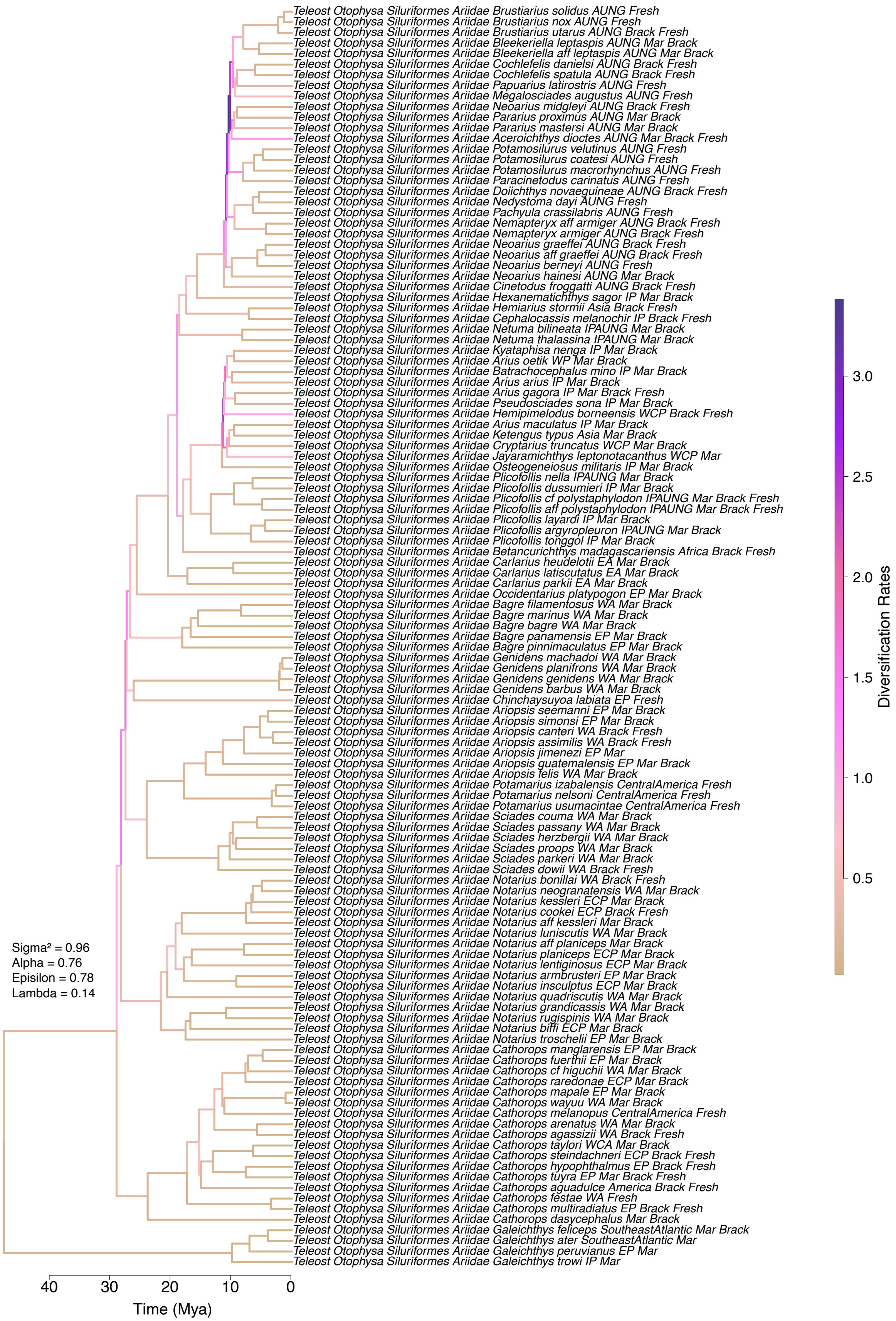

Teleost Otophysa Siluriformes Ariidae Brustiarius solidus AUNG Fresh
Teleost Otophysa Siluriformes Ariidae Brustiarius nox AUNG Fresh
Teleost Otophysa Siluriformes Ariidae Brustiarius utarus AUNG Brack Fresh
Teleost Otophysa Siluriformes Ariidae Bleekeriella leptaspis AUNG Mar Brack
Teleost Otophysa Siluriformes Ariidae Bleekeriella aff leptaspis AUNG Mar Brack
Teleost Otophysa Siluriformes Ariidae Cochlefelis danielsi AUNG Brack Fresh
Teleost Otophysa Siluriformes Ariidae Cochlefelis spatula AUNG Brack Fresh
Teleost Otophysa Siluriformes Ariidae Papuarius latirostris AUNG Fresh
Teleost Otophysa Siluriformes Ariidae Megalosciades augustus AUNG Fresh
Teleost Otophysa Siluriformes Ariidae Neoarius midgleyi AUNG Brack Fresh
Teleost Otophysa Siluriformes Ariidae Pararius proximus AUNG Mar Brack
Teleost Otophysa Siluriformes Ariidae Pararius mastersi AUNG Mar Brack
Teleost Otophysa Siluriformes Ariidae Aceroichthys dioctes AUNG Mar Brack Fresh
Teleost Otophysa Siluriformes Ariidae Potamosilurus velutinus AUNG Fresh
Teleost Otophysa Siluriformes Ariidae Potamosilurus coatesi AUNG Fresh
Teleost Otophysa Siluriformes Ariidae Potamosilurus macrorhynchus AUNG Fresh
Teleost Otophysa Siluriformes Ariidae Paracinetodus carinatus AUNG Fresh
Teleost Otophysa Siluriformes Ariidae Doiichthys novaeguineae AUNG Brack Fresh
Teleost Otophysa Siluriformes Ariidae Nedystoma dayi AUNG Fresh
Teleost Otophysa Siluriformes Ariidae Pachyula crassilabris AUNG Fresh
Teleost Otophysa Siluriformes Ariidae Nemapteryx aff armiger AUNG Brack Fresh
Teleost Otophysa Siluriformes Ariidae Nemapteryx armiger AUNG Brack Fresh
Teleost Otophysa Siluriformes Ariidae Neoarius graeffei AUNG Brack Fresh
Teleost Otophysa Siluriformes Ariidae Neoarius aff graeffei AUNG Brack Fresh
Teleost Otophysa Siluriformes Ariidae Neoarius berneyi AUNG Fresh
Teleost Otophysa Siluriformes Ariidae Neoarius hainesi AUNG Mar Brack
Teleost Otophysa Siluriformes Ariidae Cinetodus froggatti AUNG Brack Fresh
Teleost Otophysa Siluriformes Ariidae Hexanematichthys sagor IP Mar Brack
Teleost Otophysa Siluriformes Ariidae Hemiarius stormii Asia Brack Fresh
Teleost Otophysa Siluriformes Ariidae Cephalocassis melanochir IP Brack Fresh
Teleost Otophysa Siluriformes Ariidae Netuma bilineata IPAUNG Mar Brack
Teleost Otophysa Siluriformes Ariidae Netuma thalassina IPAUNG Mar Brack
Teleost Otophysa Siluriformes Ariidae Kyataphisa nenga IP Mar Brack
Teleost Otophysa Siluriformes Ariidae Arius oetik WP Mar Brack
Teleost Otophysa Siluriformes Ariidae Batrachocephalus mino IP Mar Brack
Teleost Otophysa Siluriformes Ariidae Arius arius IP Mar Brack
Teleost Otophysa Siluriformes Ariidae Arius gagora IP Mar Brack Fresh
Teleost Otophysa Siluriformes Ariidae Pseudosciades sona IP Mar Brack
Teleost Otophysa Siluriformes Ariidae Hemipimelodus borneensis WCP Brack Fresh
Teleost Otophysa Siluriformes Ariidae Arius maculatus IP Mar Brack
Teleost Otophysa Siluriformes Ariidae Ketengus typus Asia Mar Brack
Teleost Otophysa Siluriformes Ariidae Cryptarius truncatus WCP Mar Brack
Teleost Otophysa Siluriformes Ariidae Jayaramichthys leptonotacanthus WCP Mar
Teleost Otophysa Siluriformes Ariidae Osteogeneiosus militaris IP Mar Brack
Teleost Otophysa Siluriformes Ariidae Plicofollis nella IPAUNG Mar Brack
Teleost Otophysa Siluriformes Ariidae Plicofollis dussumieri IP Mar Brack
Teleost Otophysa Siluriformes Ariidae Plicofollis cf polystaphylodon IPAUNG Mar Brack Fresh
Teleost Otophysa Siluriformes Ariidae Plicofollis aff polystaphylodon IPAUNG Mar Brack Fresh
Teleost Otophysa Siluriformes Ariidae Plicofollis layardi IP Mar Brack
Teleost Otophysa Siluriformes Ariidae Plicofollis argyropleuron IPAUNG Mar Brack
Teleost Otophysa Siluriformes Ariidae Plicofollis tonggol IP Mar Brack
Teleost Otophysa Siluriformes Ariidae Betancurichthys madagascariensis Africa Brack Fresh
Teleost Otophysa Siluriformes Ariidae Carlarius heudelotii EA Mar Brack
Teleost Otophysa Siluriformes Ariidae Carlarius latiscutatus EA Mar Brack
Teleost Otophysa Siluriformes Ariidae Carlarius parkii EA Mar Brack
Teleost Otophysa Siluriformes Ariidae Occidentarius platypogon EP Mar Brack
Teleost Otophysa Siluriformes Ariidae Bagre filamentosus WA Mar Brack
Teleost Otophysa Siluriformes Ariidae Bagre marinus WA Mar Brack
Teleost Otophysa Siluriformes Ariidae Bagre bagre WA Mar Brack
Teleost Otophysa Siluriformes Ariidae Bagre panamensis EP Mar Brack
Teleost Otophysa Siluriformes Ariidae Bagre pinnimaculatus EP Mar Brack
Teleost Otophysa Siluriformes Ariidae Genidens machadoi WA Mar Brack
Teleost Otophysa Siluriformes Ariidae Genidens planifrons WA Mar Brack
Teleost Otophysa Siluriformes Ariidae Genidens genidens WA Mar Brack
Teleost Otophysa Siluriformes Ariidae Genidens barbus WA Mar Brack
Teleost Otophysa Siluriformes Ariidae Chinchaysuyoa labiata EP Fresh
Teleost Otophysa Siluriformes Ariidae Ariopsis seemanni EP Mar Brack
Teleost Otophysa Siluriformes Ariidae Ariopsis simonsi EP Mar Brack
Teleost Otophysa Siluriformes Ariidae Ariopsis canteri WA Brack Fresh
Teleost Otophysa Siluriformes Ariidae Ariopsis assimilis WA Brack Fresh
Teleost Otophysa Siluriformes Ariidae Ariopsis jimenezi EP Mar
Teleost Otophysa Siluriformes Ariidae Ariopsis guatemalensis EP Mar Brack
Teleost Otophysa Siluriformes Ariidae Ariopsis felis WA Mar Brack
Teleost Otophysa Siluriformes Ariidae Potamarius izabalensis CentralAmerica Fresh
Teleost Otophysa Siluriformes Ariidae Potamarius nelsoni CentralAmerica Fresh
Teleost Otophysa Siluriformes Ariidae Potamarius usumacintae CentralAmerica Fresh
Teleost Otophysa Siluriformes Ariidae Sciades couma WA Mar Brack
Teleost Otophysa Siluriformes Ariidae Sciades passany WA Mar Brack
Teleost Otophysa Siluriformes Ariidae Sciades herzbergii WA Mar Brack
Teleost Otophysa Siluriformes Ariidae Sciades proops WA Mar Brack
Teleost Otophysa Siluriformes Ariidae Sciades parkeri WA Mar Brack
Teleost Otophysa Siluriformes Ariidae Sciades dowii WA Brack Fresh
Teleost Otophysa Siluriformes Ariidae Notarius bonillai WA Brack Fresh
Teleost Otophysa Siluriformes Ariidae Notarius neogranatensis WA Mar Brack
Teleost Otophysa Siluriformes Ariidae Notarius kessleri ECP Mar Brack
Teleost Otophysa Siluriformes Ariidae Notarius cookei ECP Brack Fresh
Teleost Otophysa Siluriformes Ariidae Notarius aff kessleri Mar Brack
Teleost Otophysa Siluriformes Ariidae Notarius luniscutis WA Mar Brack
Teleost Otophysa Siluriformes Ariidae Notarius aff planiceps Mar Brack
Teleost Otophysa Siluriformes Ariidae Notarius planiceps ECP Mar Brack
Teleost Otophysa Siluriformes Ariidae Notarius lentiginosus ECP Mar Brack
Teleost Otophysa Siluriformes Ariidae Notarius armbrusteri EP Mar Brack
Teleost Otophysa Siluriformes Ariidae Notarius insculptus ECP Mar Brack
Teleost Otophysa Siluriformes Ariidae Notarius quadriscutis WA Mar Brack
Teleost Otophysa Siluriformes Ariidae Notarius grandicassis WA Mar Brack
Teleost Otophysa Siluriformes Ariidae Notarius rugispinis WA Mar Brack
Teleost Otophysa Siluriformes Ariidae Notarius biffi ECP Mar Brack
Teleost Otophysa Siluriformes Ariidae Notarius troschelii EP Mar Brack
Teleost Otophysa Siluriformes Ariidae Cathorops manglarensis EP Mar Brack
Teleost Otophysa Siluriformes Ariidae Cathorops fuerthii EP Mar Brack
Teleost Otophysa Siluriformes Ariidae Cathorops cf higuchii WA Mar Brack
Teleost Otophysa Siluriformes Ariidae Cathorops raredonae ECP Mar Brack
Teleost Otophysa Siluriformes Ariidae Cathorops mapale EP Mar Brack
Teleost Otophysa Siluriformes Ariidae Cathorops wayuu WA Mar Brack
Teleost Otophysa Siluriformes Ariidae Cathorops melanopus CentralAmerica Fresh
Teleost Otophysa Siluriformes Ariidae Cathorops arenatus WA Mar Brack
Teleost Otophysa Siluriformes Ariidae Cathorops agassizii WA Brack Fresh
Teleost Otophysa Siluriformes Ariidae Cathorops taylori WCA Mar Brack
Teleost Otophysa Siluriformes Ariidae Cathorops steindachneri ECP Brack Fresh
Teleost Otophysa Siluriformes Ariidae Cathorops hypophthalmus EP Brack Fresh
Teleost Otophysa Siluriformes Ariidae Cathorops tuyra EP Mar Brack Fresh
Teleost Otophysa Siluriformes Ariidae Cathorops aguadulce America Brack Fresh
Teleost Otophysa Siluriformes Ariidae Cathorops festae WA Fresh
Teleost Otophysa Siluriformes Ariidae Cathorops multiradiatus EP Brack Fresh
Teleost Otophysa Siluriformes Ariidae Cathorops dasycephalus Mar Brack
Teleost Otophysa Siluriformes Ariidae Galeichthys feliceps SoutheastAtlantic Mar Brack
Teleost Otophysa Siluriformes Ariidae Galeichthys ater SoutheastAtlantic Mar
Teleost Otophysa Siluriformes Ariidae Galeichthys peruvianus EP Mar
Teleost Otophysa Siluriformes Ariidae Galeichthys trowi IP Mar
Sigma² = 0.96
Alpha = 0.76
Episilon = 0.78
Lambda = 0.14
40
30
20
10
0
Time (Mya)
Diversification Rates
3.0
2.5
2.0
1.5
1.0
0.5

321

**Figure S17. Ariidae (focal family) clade pruned from the complete trees (n = 2,303 species) with speciation rate shifts based on ClaDS**. ClaDS analyses were performed using ASTRAL and RAxML MTs (see parameters in Table S32). Here, we show ClaDS on ASTRAL MT topology.

**Figure S18. Atheriniformes clade pruned from the complete trees (n = 2,303 species) with speciation rate shifts based on ClaDS.** Focal families: Atherinidae, Melanotaeniidae, and Pseudomugilidae. ClaDS analyses were performed using ASTRAL and RAxML MTs (see parameters in Table S32). Here, we show ClaDS on ASTRAL MT topology. High resolution figure available in supplementary files.

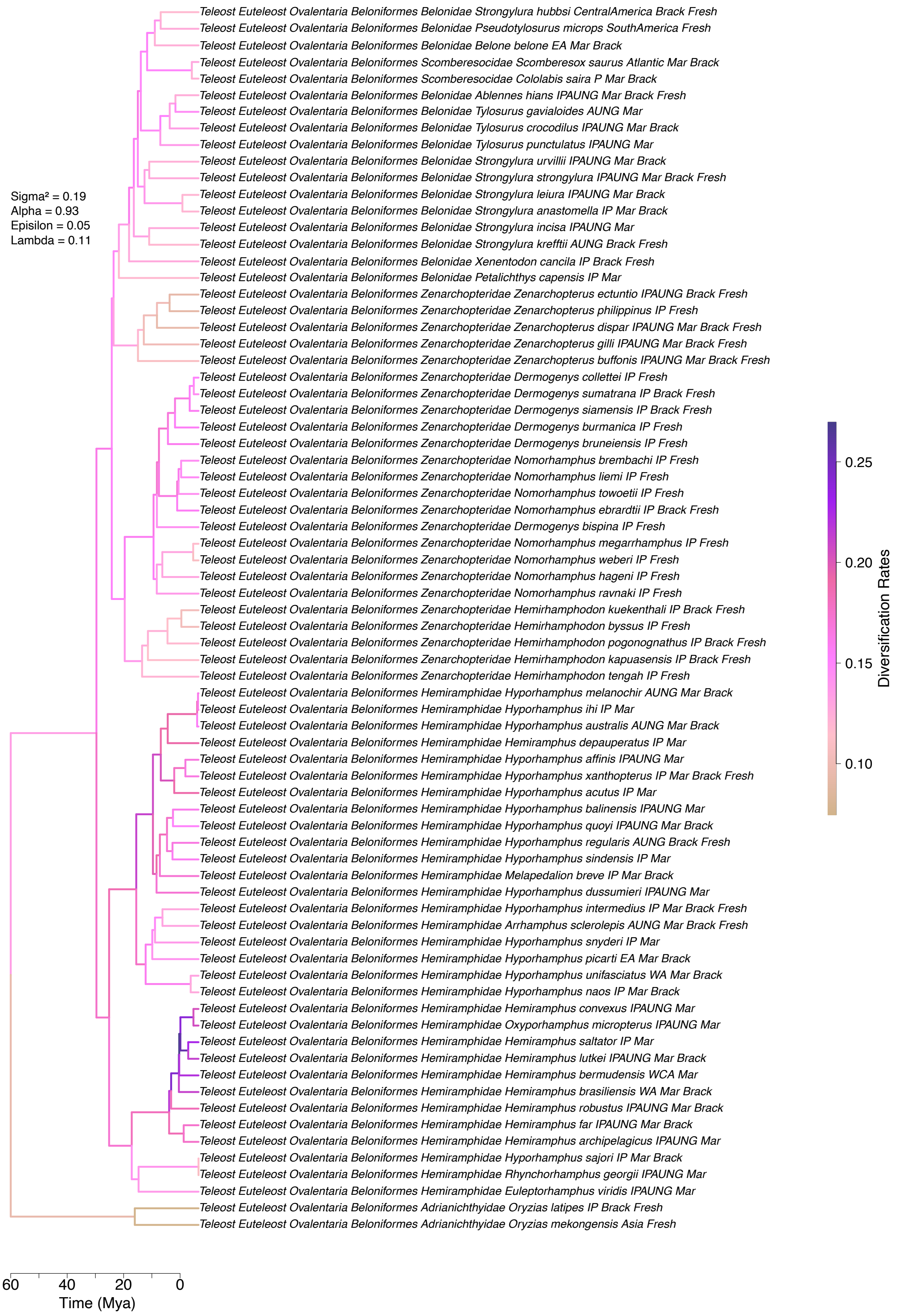
Teleost Euteleost Ovalentaria Beloniformes Belonidae Strongylura hubbsi CentralAmerica Brack Fresh
Teleost Euteleost Ovalentaria Beloniformes Belonidae Pseudotylosurus microps SouthAmerica Fresh
Teleost Euteleost Ovalentaria Beloniformes Belonidae Belone belone EA Mar Brack
Teleost Euteleost Ovalentaria Beloniformes Scomberesocidae Scomberesox saurus Atlantic Mar Brack
Teleost Euteleost Ovalentaria Beloniformes Scomberesocidae Cololabis saira P Mar Brack
Teleost Euteleost Ovalentaria Beloniformes Belonidae Ablennes hians IPAUNG Mar Brack Fresh
Teleost Euteleost Ovalentaria Beloniformes Belonidae Tylosurus gavialoides AUNG Mar
Teleost Euteleost Ovalentaria Beloniformes Belonidae Tylosurus crocodilus IPAUNG Mar Brack
Teleost Euteleost Ovalentaria Beloniformes Belonidae Tylosurus punctulatus IPAUNG Mar
Teleost Euteleost Ovalentaria Beloniformes Belonidae Strongylura urvillii IPAUNG Mar Brack
Teleost Euteleost Ovalentaria Beloniformes Belonidae Strongylura strongylura IPAUNG Mar Brack Fresh
Teleost Euteleost Ovalentaria Beloniformes Belonidae Strongylura leiura IPAUNG Mar Brack
Teleost Euteleost Ovalentaria Beloniformes Belonidae Strongylura anastomella IP Mar Brack
Teleost Euteleost Ovalentaria Beloniformes Belonidae Strongylura incisa IPAUNG Mar
Teleost Euteleost Ovalentaria Beloniformes Belonidae Strongylura krefftii AUNG Brack Fresh
Teleost Euteleost Ovalentaria Beloniformes Belonidae Xenentodon cancila IP Brack Fresh
Teleost Euteleost Ovalentaria Beloniformes Belonidae Petalichthys capensis IP Mar
Teleost Euteleost Ovalentaria Beloniformes Zenarchopteridae Zenarchopterus ectuntio IPAUNG Brack Fresh
Teleost Euteleost Ovalentaria Beloniformes Zenarchopteridae Zenarchopterus philippinus IP Fresh
Teleost Euteleost Ovalentaria Beloniformes Zenarchopteridae Zenarchopterus dispar IPAUNG Mar Brack Fresh
Teleost Euteleost Ovalentaria Beloniformes Zenarchopteridae Zenarchopterus gilli IPAUNG Mar Brack Fresh
Teleost Euteleost Ovalentaria Beloniformes Zenarchopteridae Zenarchopterus buffonis IPAUNG Mar Brack Fresh
Teleost Euteleost Ovalentaria Beloniformes Zenarchopteridae Dermogenys collettei IP Fresh
Teleost Euteleost Ovalentaria Beloniformes Zenarchopteridae Dermogenys sumatrana IP Brack Fresh
Teleost Euteleost Ovalentaria Beloniformes Zenarchopteridae Dermogenys siamensis IP Brack Fresh
Teleost Euteleost Ovalentaria Beloniformes Zenarchopteridae Dermogenys burmanica IP Fresh
Teleost Euteleost Ovalentaria Beloniformes Zenarchopteridae Dermogenys bruneiensis IP Fresh
Teleost Euteleost Ovalentaria Beloniformes Zenarchopteridae Nomorhamphus brembachi IP Fresh
Teleost Euteleost Ovalentaria Beloniformes Zenarchopteridae Nomorhamphus liemi IP Fresh
Teleost Euteleost Ovalentaria Beloniformes Zenarchopteridae Nomorhamphus towoetii IP Fresh
Teleost Euteleost Ovalentaria Beloniformes Zenarchopteridae Nomorhamphus ebrardtii IP Brack Fresh
Teleost Euteleost Ovalentaria Beloniformes Zenarchopteridae Dermogenys bispina IP Fresh
Teleost Euteleost Ovalentaria Beloniformes Zenarchopteridae Nomorhamphus megarrhamphus IP Fresh
Teleost Euteleost Ovalentaria Beloniformes Zenarchopteridae Nomorhamphus weberi IP Fresh
Teleost Euteleost Ovalentaria Beloniformes Zenarchopteridae Nomorhamphus hageni IP Fresh
Teleost Euteleost Ovalentaria Beloniformes Zenarchopteridae Nomorhamphus ravnaki IP Fresh
Teleost Euteleost Ovalentaria Beloniformes Zenarchopteridae Hemirhamphodon kuekenthali IP Brack Fresh
Teleost Euteleost Ovalentaria Beloniformes Zenarchopteridae Hemirhamphodon byssus IP Fresh
Teleost Euteleost Ovalentaria Beloniformes Zenarchopteridae Hemirhamphodon pogonognathus IP Brack Fresh
Teleost Euteleost Ovalentaria Beloniformes Zenarchopteridae Hemirhamphodon kapuasensis IP Brack Fresh
Teleost Euteleost Ovalentaria Beloniformes Zenarchopteridae Hemirhamphodon tengah IP Fresh
Teleost Euteleost Ovalentaria Beloniformes Hemiramphidae Hyporhamphus melanochir AUNG Mar Brack
Teleost Euteleost Ovalentaria Beloniformes Hemiramphidae Hyporhamphus ihi IP Mar
Teleost Euteleost Ovalentaria Beloniformes Hemiramphidae Hyporhamphus australis AUNG Mar Brack
Teleost Euteleost Ovalentaria Beloniformes Hemiramphidae Hemiramphus depauperatus IP Mar
Teleost Euteleost Ovalentaria Beloniformes Hemiramphidae Hyporhamphus affinis IPAUNG Mar
Teleost Euteleost Ovalentaria Beloniformes Hemiramphidae Hyporhamphus xanthopterus IP Mar Brack Fresh
Teleost Euteleost Ovalentaria Beloniformes Hemiramphidae Hyporhamphus acutus IP Mar
Teleost Euteleost Ovalentaria Beloniformes Hemiramphidae Hyporhamphus balinensis IPAUNG Mar
Teleost Euteleost Ovalentaria Beloniformes Hemiramphidae Hyporhamphus quoyi IPAUNG Mar Brack
Teleost Euteleost Ovalentaria Beloniformes Hemiramphidae Hyporhamphus regularis AUNG Brack Fresh
Teleost Euteleost Ovalentaria Beloniformes Hemiramphidae Hyporhamphus sindensis IP Mar
Teleost Euteleost Ovalentaria Beloniformes Hemiramphidae Melapedalion breve IP Mar Brack
Teleost Euteleost Ovalentaria Beloniformes Hemiramphidae Hyporhamphus dussumieri IPAUNG Mar
Teleost Euteleost Ovalentaria Beloniformes Hemiramphidae Hyporhamphus intermedius IP Mar Brack Fresh
Teleost Euteleost Ovalentaria Beloniformes Hemiramphidae Arrhamphus sclerolepis AUNG Mar Brack Fresh
Teleost Euteleost Ovalentaria Beloniformes Hemiramphidae Hyporhamphus snyderi IP Mar
Teleost Euteleost Ovalentaria Beloniformes Hemiramphidae Hyporhamphus picarti EA Mar Brack
Teleost Euteleost Ovalentaria Beloniformes Hemiramphidae Hyporhamphus unifasciatus WA Mar Brack
Teleost Euteleost Ovalentaria Beloniformes Hemiramphidae Hyporhamphus naos IP Mar Brack
Teleost Euteleost Ovalentaria Beloniformes Hemiramphidae Hemiramphus convexus IPAUNG Mar
Teleost Euteleost Ovalentaria Beloniformes Hemiramphidae Oxyporhamphus micropterus IPAUNG Mar
Teleost Euteleost Ovalentaria Beloniformes Hemiramphidae Hemiramphus saltator IP Mar
Teleost Euteleost Ovalentaria Beloniformes Hemiramphidae Hemiramphus lutkei IPAUNG Mar Brack
Teleost Euteleost Ovalentaria Beloniformes Hemiramphidae Hemiramphus bermudensis WCA Mar
Teleost Euteleost Ovalentaria Beloniformes Hemiramphidae Hemiramphus brasiliensis WA Mar Brack
Teleost Euteleost Ovalentaria Beloniformes Hemiramphidae Hemiramphus robustus IPAUNG Mar Brack
Teleost Euteleost Ovalentaria Beloniformes Hemiramphidae Hemiramphus far IPAUNG Mar Brack
Teleost Euteleost Ovalentaria Beloniformes Hemiramphidae Hemiramphus archipelagicus IPAUNG Mar
Teleost Euteleost Ovalentaria Beloniformes Hemiramphidae Hyporhamphus sajori IP Mar Brack
Teleost Euteleost Ovalentaria Beloniformes Hemiramphidae Rhynchorhamphus georgii IPAUNG Mar
Teleost Euteleost Ovalentaria Beloniformes Hemiramphidae Euleptorhamphus viridis IPAUNG Mar
Teleost Euteleost Ovalentaria Beloniformes Adrianichthyidae Oryzias latipes IP Brack Fresh
Teleost Euteleost Ovalentaria Beloniformes Adrianichthyidae Oryzias mekongensis Asia Fresh
Sigma² = 0.19
Alpha = 0.93
Episilon = 0.05
Lambda = 0.11
0.25
0.20
0.15
0.10
Diversification Rates
60
40
20
0
Time (Mya)

**Figure S19. Beloniformes clade pruned from the complete trees (n = 2,303 species) with speciation rate shifts based on ClaDS.** Focal family: Zenarchopteridae. ClaDS analyses were performed using ASTRAL and RAxML MTs (see parameters in Table S32). Here, we show ClaDS on ASTRAL MT topology.

**Figure S20. Carangaria clade pruned from the complete trees (n = 2,303 species) with speciation rate shifts based on ClaDS.** Focal family: Toxotidae. ClaDS analyses were performed using ASTRAL and RAxML MTs (see parameters in Table S32). Here, we show ClaDS on ASTRAL MT topology. High resolution figure available in supplementary files.

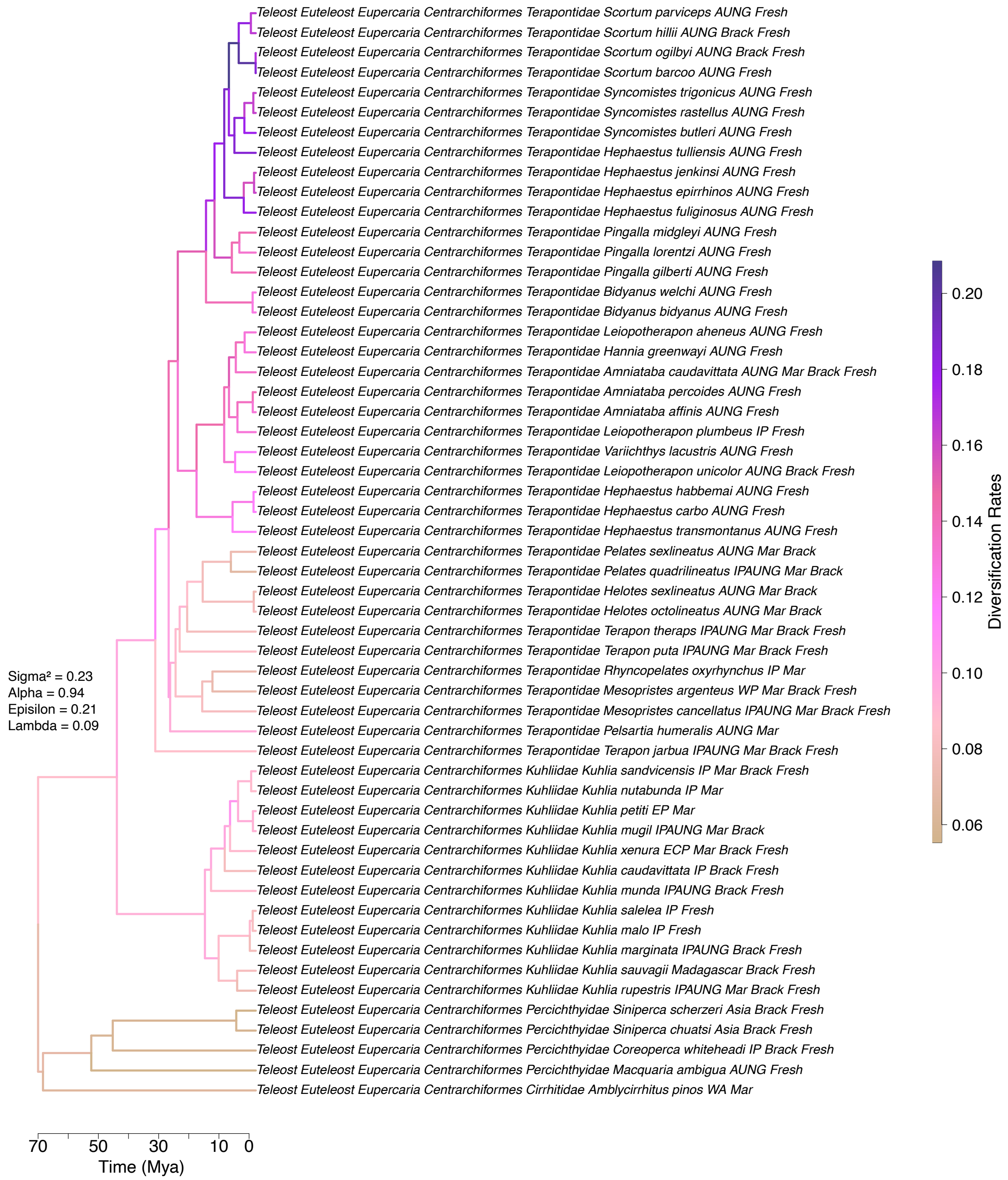

**Figure S21. Centrarchiformes clade pruned from the complete trees (n = 2,303 species) with speciation rate shifts based on ClaDS**. Focal families: Kuhliidae and Terapontidae. ClaDS analyses were performed using ASTRAL and RAxML MTs (see parameters in Table S32). Here, we show ClaDS on ASTRAL MT topology.

**Figure S22. Clupeiformes clade pruned from the complete trees (n = 2,303 species) with speciation rate shifts based on ClaDS.** Focal family: Clupeidae. ClaDS analyses were performed using ASTRAL and RAxML MTs (see parameters in Table S32). Here, we show ClaDS on ASTRAL MT topology. High resolution figure available in supplementary files.

**Figure S23. Gobiiformes clade pruned from the complete trees (n = 2,303 species) with speciation rate shifts based on ClaDS.** Focal families: Butidae, Eleotridae, Gobiidae, and Oxudercidae. ClaDS analyses were performed using ASTRAL and RAxML MTs (see parameters in Table S32). Here, we show ClaDS on ASTRAL MT topology. High resolution figure available in supplementary files.

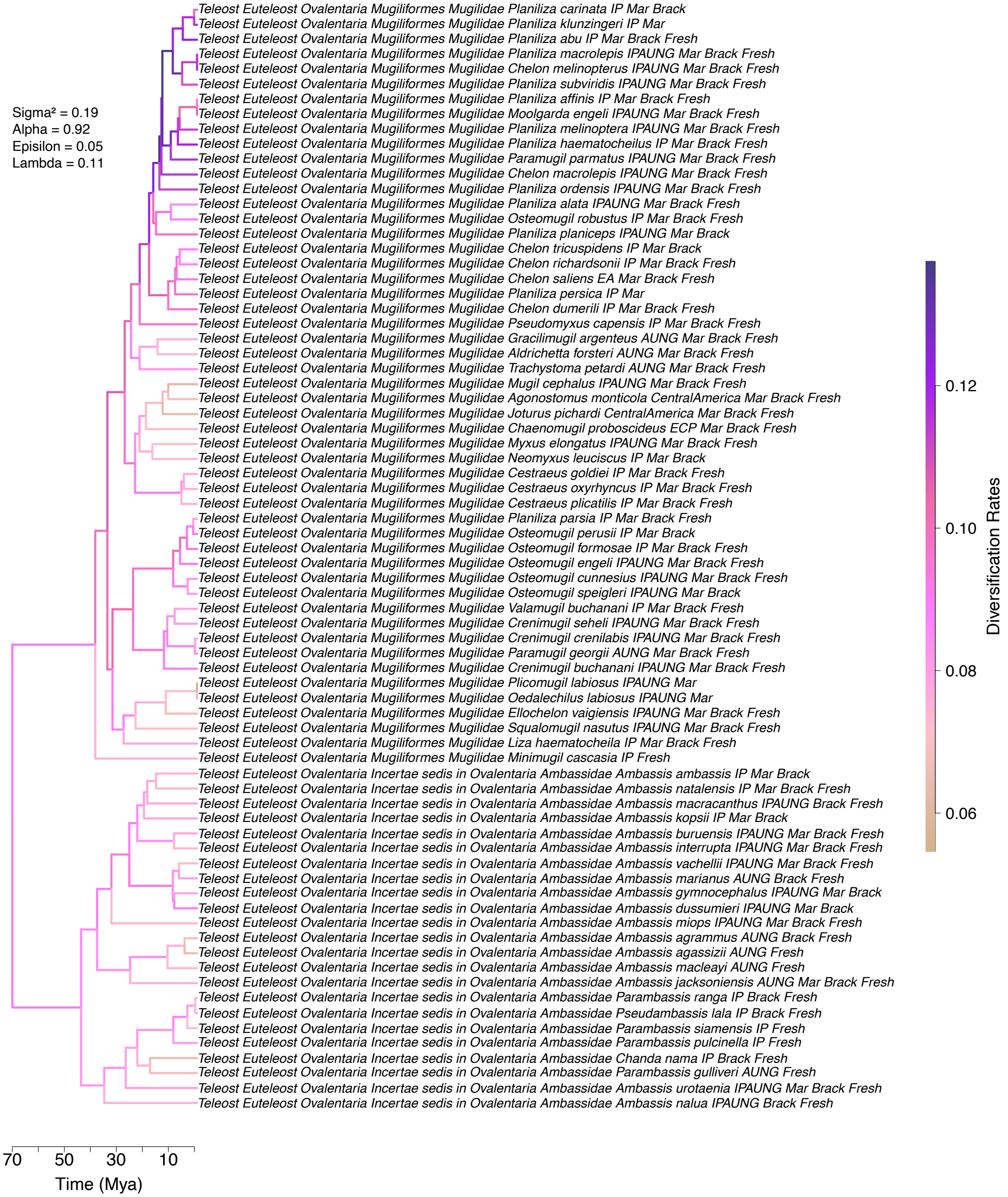

**Figure S24. Mugiliformes clade pruned from the complete trees (n = 2,303 species) with speciation rate shifts based on ClaDS.** Focal families: Ambassidae and Mugilidae. ClaDS analyses were performed using ASTRAL and RAxML MTs (see parameters in Table S32). Here, we show ClaDS on ASTRAL MT topology.

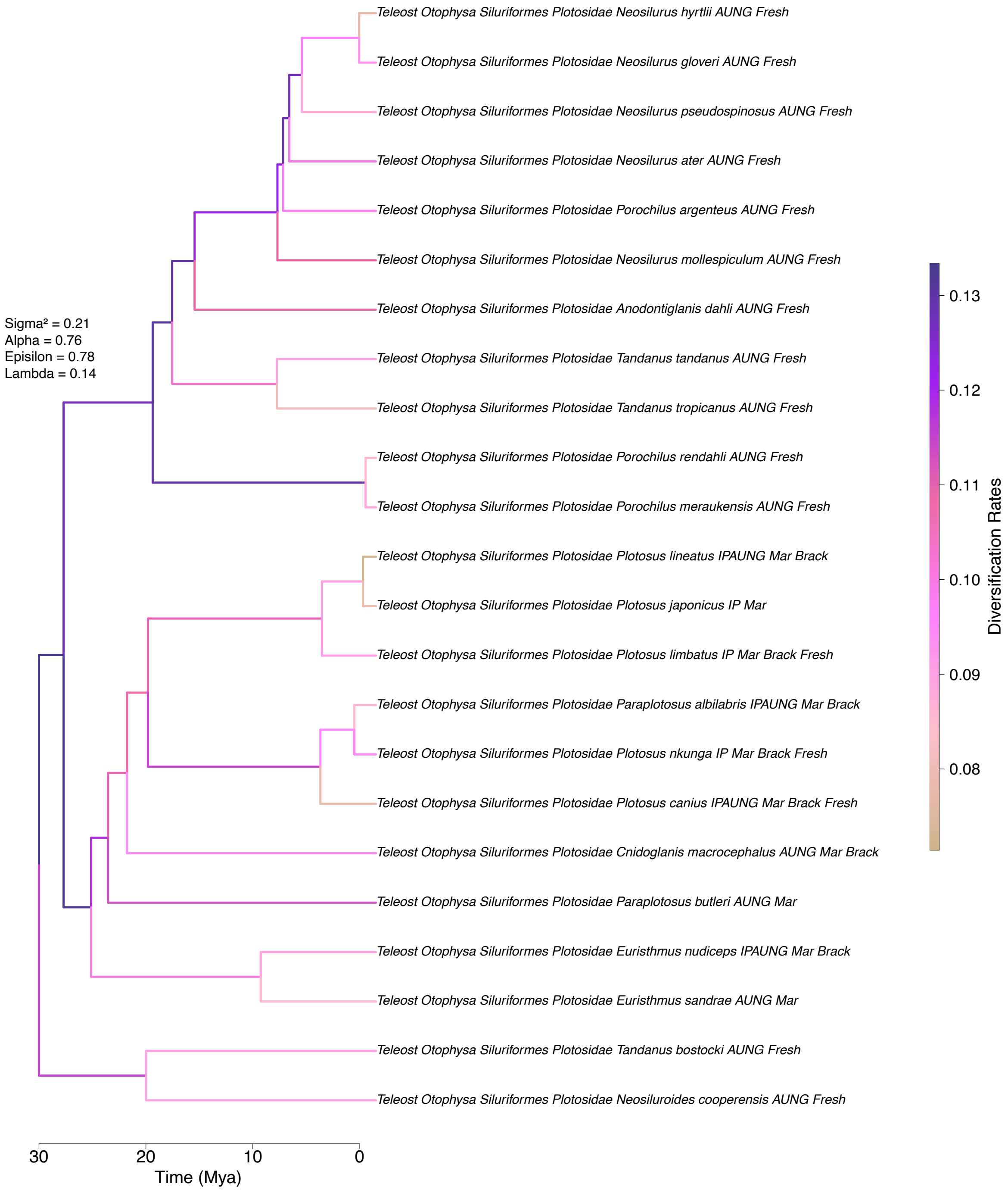

**Figure S25. Plotosidae (focal family) clade pruned from the complete trees (n = 2,303 species) with speciation rate shifts based on ClaDS**. ClaDS analyses were performed using ASTRAL and RAxML MTs (see parameters in Table S32). Here, we show ClaDS on ASTRAL MT topology.

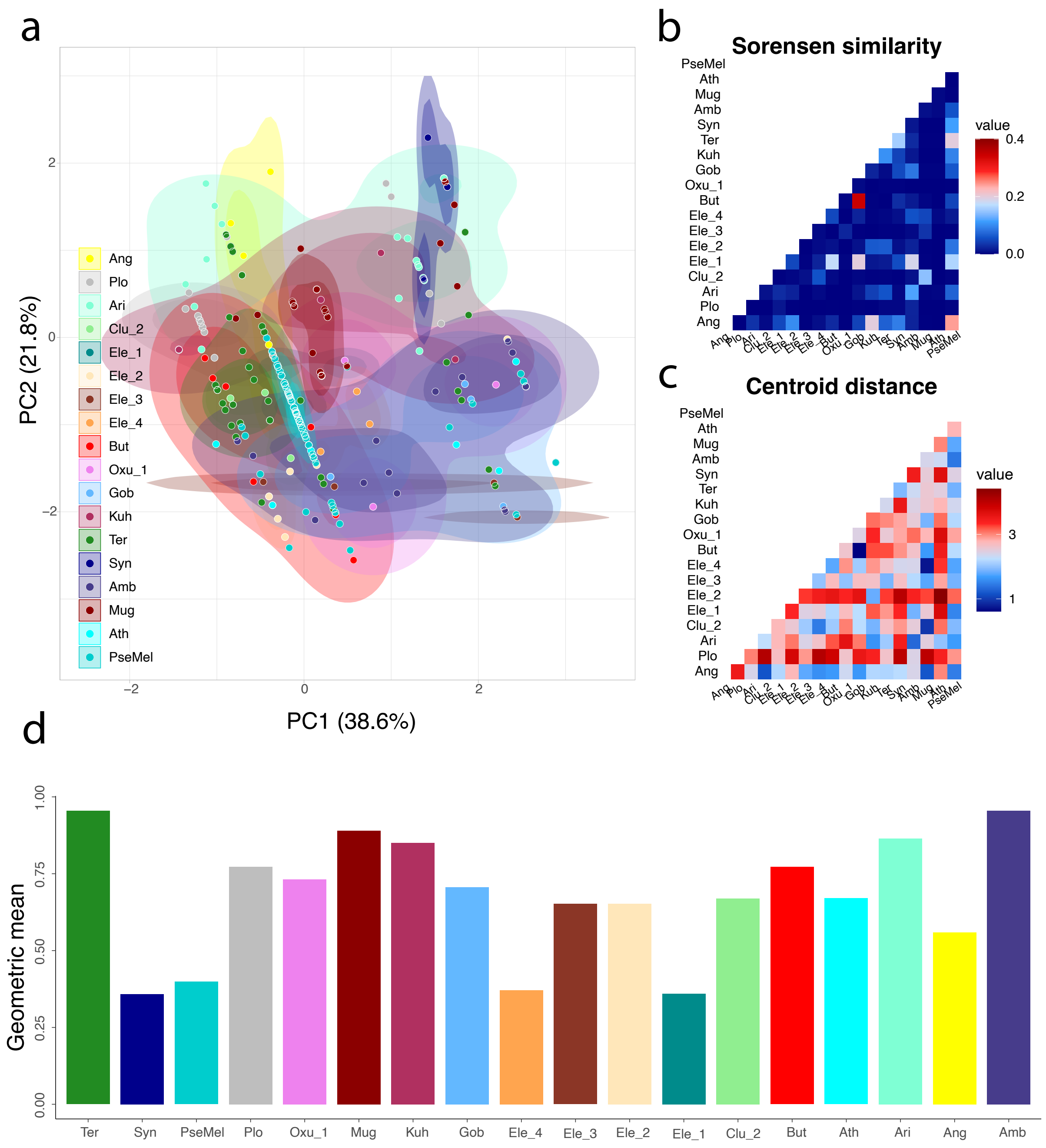

**Figure S26. Hypervolume and principal component analyses for 18 subclades with ≥3 species.** (a) Functional space of PC1 and PC2 with points and slices of hypervolume colored by subclades. Darker shades within each shaded hypervolume represent the region where the local estimated point density is at least two thirds of the maximum point density, whereas the lighter shaded region has a local estimated point density of at least one third of the maximum estimated point density. (b) Heatmap plot of Sorensen similarity between overlap hypervolumes, which corresponds to the ratio size of the intersection to the sum of individual hypervolumes for two distinct subclades. (c) Heatmap plot of centroid distances, measured as the Euclidean distance between hypervolume centroids.

(d) Hypervolume size for each subclade represented by the geometric mean (central tendency of per-dimension ranges). Ang: Anguillidae; Plo: Plotosidae; Ari: Ariidae; Clu_2: Clupeidae subclade ‘2’; Ele_1: Eleotridae subclade ‘1’; Ele_2: Eleotridae subclade ‘2’; Ele_3: Eleotridae subclade ‘3’; Ele_4: Eleotridae subclade ‘4’; But: Butidae; Oxu_1: Oxudercidae subclade ‘1’; Gob: Gobiidae; Kuh: Kuhliidae; Ter: Terapontidae; Syn: Synbranchidae; Amb: Ambassidae; Mug: Mugilidae; Ath: Atherinidae; PseMel: Pseudomugilidae & Melanotaeniidae.

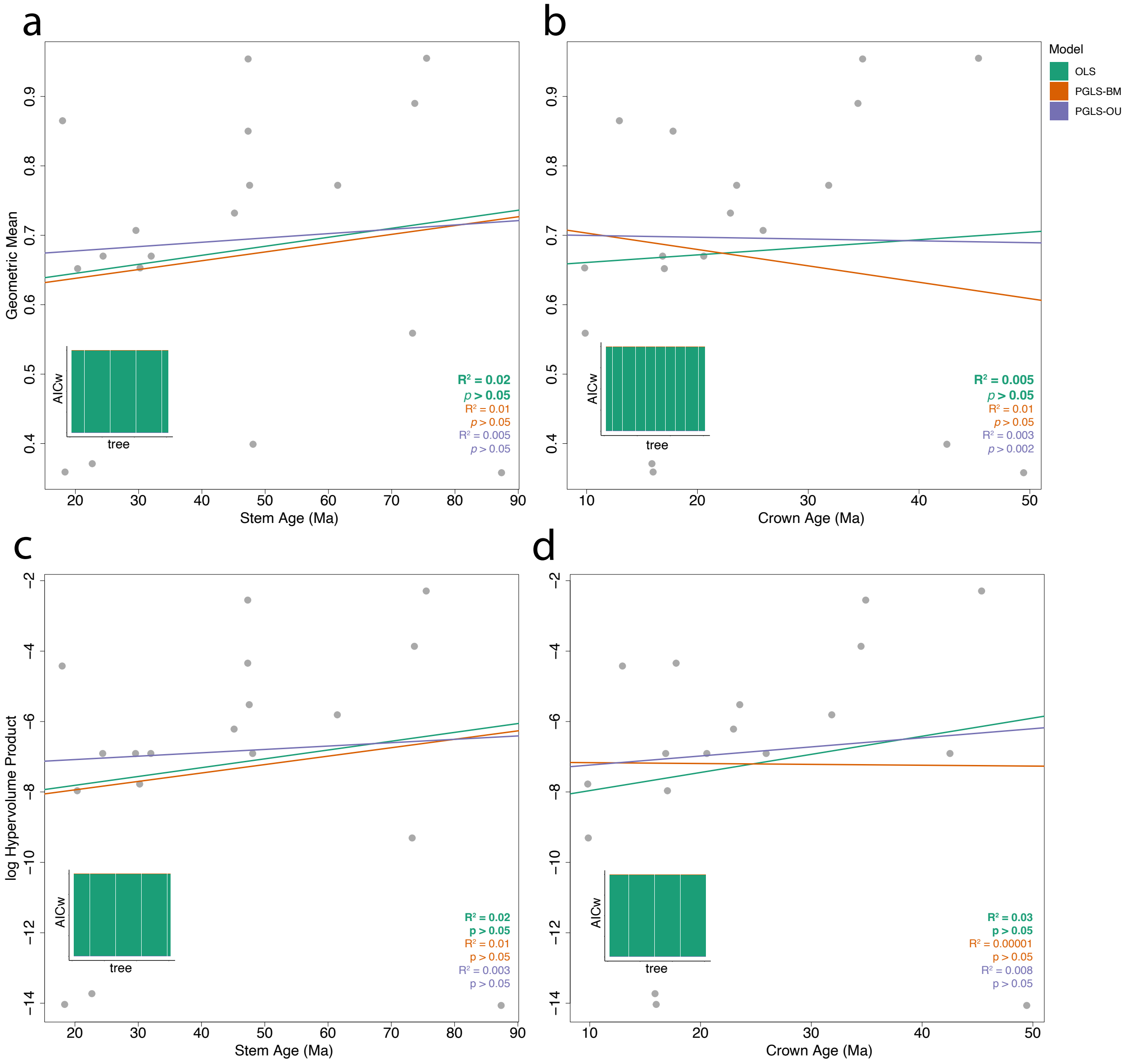

**Figure S27. Ordinary (OLS) and phylogenetic (PGLS) least-square under Brownian motion (PGLS-BM) and Ornstein–Uhlenbeck process (PGLS-OU) regressions between subclade ages and functional space using ASTRAL MT.** (a) Regression between stem age and functional space measured by the hypervolume geometric mean. (b) Regression between crown age and functional space measured by the hypervolume geometric mean. (c) Regression between stem age and functional space measured by the log-transformed hypervolume product. (d) Regression between crown age and functional space measured by the log-transformed hypervolume product. Averaged R-squared and significance (p) are provided in the lower right corner emphasizing (in bold) the best-fit model. PGLS regressions were conducted for all 30 trees.

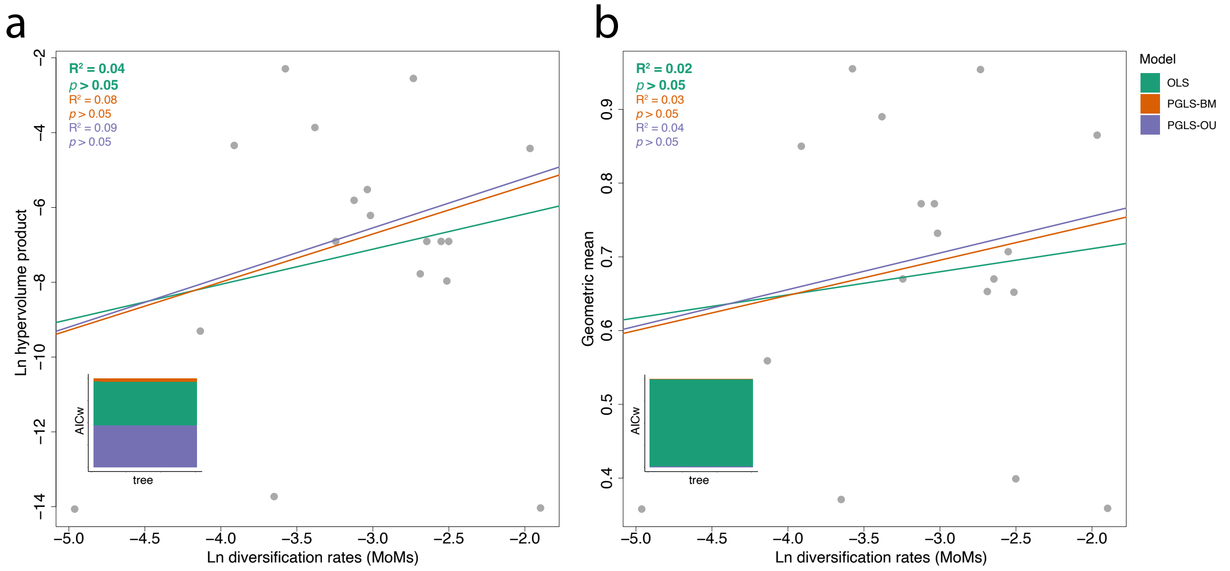

**Figure S28. Ordinary (OLS) and phylogenetic (PGLS) least-square under Brownian motion (PGLS-BM) and Ornstein–Uhlenbeck process (PGLS-OU) regressions between diversification rates and functional space for 18 subclades using ASTRAL MT.** (a) Regression between diversification rate based on Method-of-Moments (MoMs) and functional space measured by the hypervolume product. (b) Regression between diversification rate based on MoMs and functional space measured by the hypervolume geometric mean. Averaged R-squared and significance (p) are provided in the upper left corner emphasizing (in bold) the best-fit model. PGLS regressions were conducted for all 30 trees.

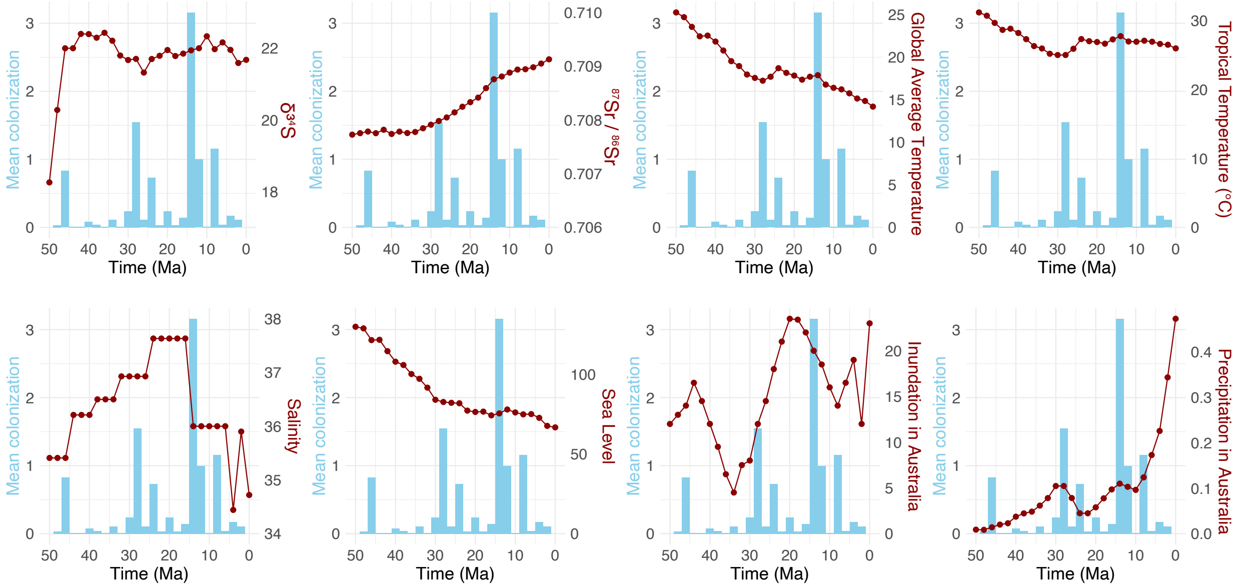

**Figure S29. Colonization events and paleoenvironmental range through time.** Blue bars show the mean number of colonization events based on the ASTRAL MT tree (see Table S19 for counts under different stochastic analyses). Red dots show the values of the eight paleoenvironmental variables used in the generalized additive models (GAMs). The x-axis is time in million years (Ma).

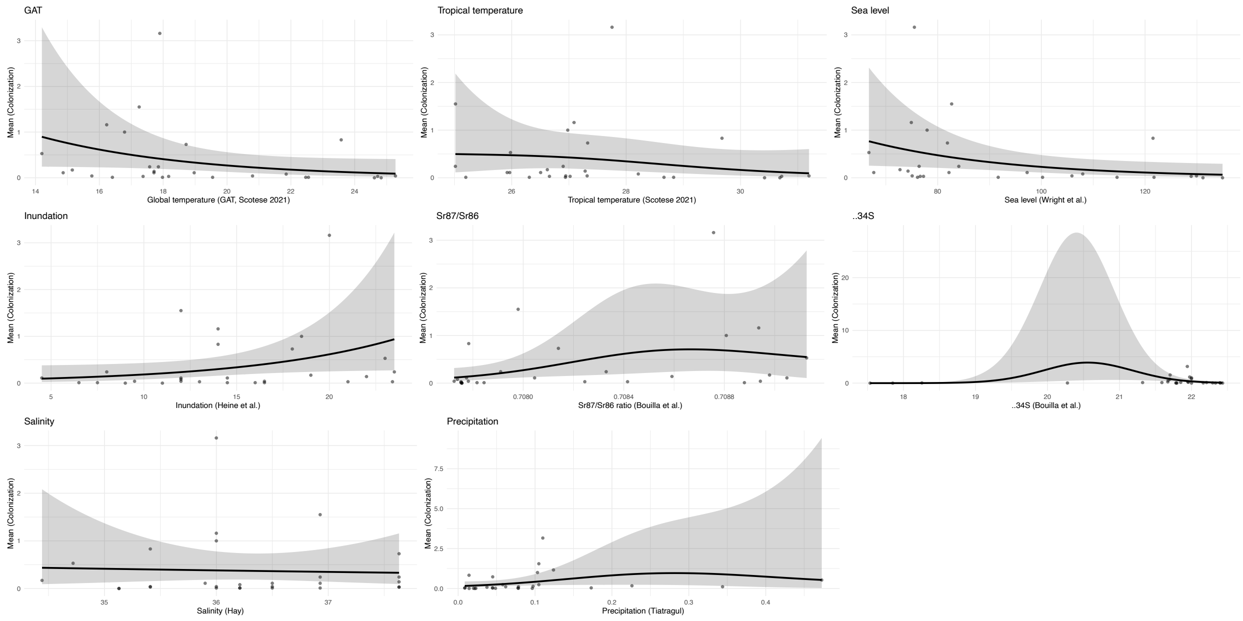

**Figure S30. Generalized Additive Models (GAM) showing the correlation between the mean number of colonization events and eight paleoenvironmental variables.** Mean numbers of colonization are retrieved from a set of 100 stochastic maps based on ASTRAL MT and Fishbase Reconciled Fresh+ dataset. Here, we show the correlations of univariate (single trait) models with no time lag. We repeated the GAM analyses with

20 different datasets, univariate and multivariate models, and different time lags. All analyses' results are available in Table S19.
